# Transcriptome analysis during fruit developmental stages in durian (*Durio zibethinus* Murr.) var. D24

**DOI:** 10.1101/2022.10.08.511399

**Authors:** Nurul Arneida Husin

**Author notes:** Corresponding author’s name: Dr. Nurul Arneida Husin, Full postal address: Biomedical Research Laboratories, School of Medicine and Health Sciences, Monash University Malaysia, Jalan Lagoon Selatan, 47500, Bandar Sunway, Selangor.

## Abstract

Durian (*Durio zibethinus* Murr.) fruits are famous for their unique aroma. This study analysed the Durian fruit transcriptome to discover the expression patterns of genes and to understand their regulation. Three developmental stages of Durian fruit, namely, early [90 days post-anthesis (DPA)], mature (120 DPA), and ripen (127 DPA), were studied. The Illumina HiSeq platform was used for sequencing. The sequence data were analysed using four different mapping aligners and statistical methods: CLC Genomic Workbench, HISAT2+DESeq2, Tophat+Cufflinks, and HISAT2+edgeR. The analyses showed that over 110 million clean reads were mapped to the Durian genome, yielding 19,976, 11,394, 17,833, and 24,351 differentially expressed genes during 90-127 days post-anthesis. Many identified differentially expressed genes were linked to the fruit ripening processes. The data analysis suggests that most genes with increased expression at the ripening stage were primarily involved in the metabolism of cofactors and vitamins, nucleotide metabolism, and carbohydrate metabolism. Significantly expressed genes from the young to mature stage were mainly associated with carbohydrate metabolism, amino acid metabolism, and cofactor and vitamin metabolism. The transcriptome data will serve as a foundation for understanding Durian fruit development-specific genes and could be helpful in fruit’s trait improvement.

## Introduction

Durian fruits are extremely popular in South-East Asia. Durian fruit is known as the “King of Fruits” because of its remarkable flavour and aroma. Both traditional and modern breeding approaches can be used to improve the quality of the fruits. In this line, understanding the gene expression patterns that occur during the development of durian fruit is critical for its further quality improvement. The texture and aroma are the two primary quality characteristics that define the market demand for durian fruits from the consumer’s perspective (Husin *et al*., 2018).

Premature fruit ripening causes significant economic losses for both farmers and consumers alike. The durian is a climacteric fruit that softens intensively during the last stages of development. The pulp softening is regulated by endogenous ethylene. However, this process involves several associated genes that are regulated mainly for fruit softening in durian. The ability to regulate fruit softening would have economic benefits for durian growers. Hence, genetic engineering tools for delaying maturation and softening processes can be used as desirable (Bouzayen *et al*., 2010; Asif *et al*., 2014).

Ripening is accompanied by increased polygalacturonase and galactosidase activity (Imsabai *et al*., 2002). Hydrolysis activity on the fruit cell wall polysaccharides leads to modifying cell wall composition (Brownleader *et al*., 1999). Cell wall polymer degradation is facilitated by various enzymes such as cellulase, polygalacturonase, β-galactosidase, pectate lyase, and xyloglucan endotransglycosylase/hydrolase (XTH) (Han *et al*., 2016).

As of January 2021, 69,566 nucleotide sequences, 44,924 genes, 65,547 proteins, and 14 SRA from *Durio zibethinus* were deposited by researchers to NCBI GenBank. These DNA and mRNA sequences are helpful in the identification of genes and or transcripts. The genome draft sequence of the Musang King durian cultivar has been made public (Teh *et al*., 2017), and most recently, long PacBio reads to sequence the Musang King var. chloroplast genome has been reported by Shearman *et al*., 2020.

Transcriptomic studies are essential in understanding multiple pathways, gene expression patterns, and secondary metabolites biosynthesis in developing fruits or tissues of interest. Several studies on gene expression patterns and gene expression profiles at the molecular level in Durian varieties have been reported (Nyffeler and Baum, 2000; Ruwaida, 2009; Vanijajiva, 2011, 2012; Indonesia *et al*., 2012; Hariyati *et al*., 2013; Santoso and Saleh, 2013; Posoongnoen *et al*., 2015; Teh *et al*., 2017; Husin *et al*., 2018). In transcriptomics of Durian, Musang King var., ripening-related gene sets included genes regulated by the MADS-BOX transcription factor family 29, SEPALLATA transcription factor family, and ethylene-related genes such as ACS (aminocyclopropane-1-carboxylic acid synthase), a key ethylene-production enzyme involved in ripening (Teh *et al*., 2017). On the other hand, high-throughput sequencing has not been employed to analyse differential gene expression during development or to compare stages of Durian growth. Hence, the research findings and data reported in this paper will contribute to a better understanding of durian fruit development.

The ultimate goal of this study was to contribute fundamental knowledge about durian fruit development and ripening from a gene expression perspective. We have investigated the transcriptomic variations in Durian by performing differential gene (DEG) discovery and gene expression characterisation in edible fruit pulp in its young, mature, and ripened stages. We identified genes involved in ethylene biosynthesis, fruit softening, and other metabolic pathways essential for the development and ripening of durian fruit. The transcriptomic data and its analysis are reported in this paper.

## Materials and Methods

### Plant Material

Fruit pulp samples from three different stages (young, mature and ripening) of Durian variety (clone D24) fruits were collected freshly in triplicate at Agriculture Park of Universiti Putra Malaysia (UPM), Selangor, Malaysia (Figure 1). Fruit pulp tissues were separated from the husk (and seeds), frozen in liquid nitrogen, and stored at □80 °C until the total RNA isolation. The young fruits were collected on the 90^th^ day after anthesis. The mature fruits were left at room temperature for seven days to let them ripe naturally. The fruit pulp tissue was separated from the ripened fruit stage.

**Figure 1.**
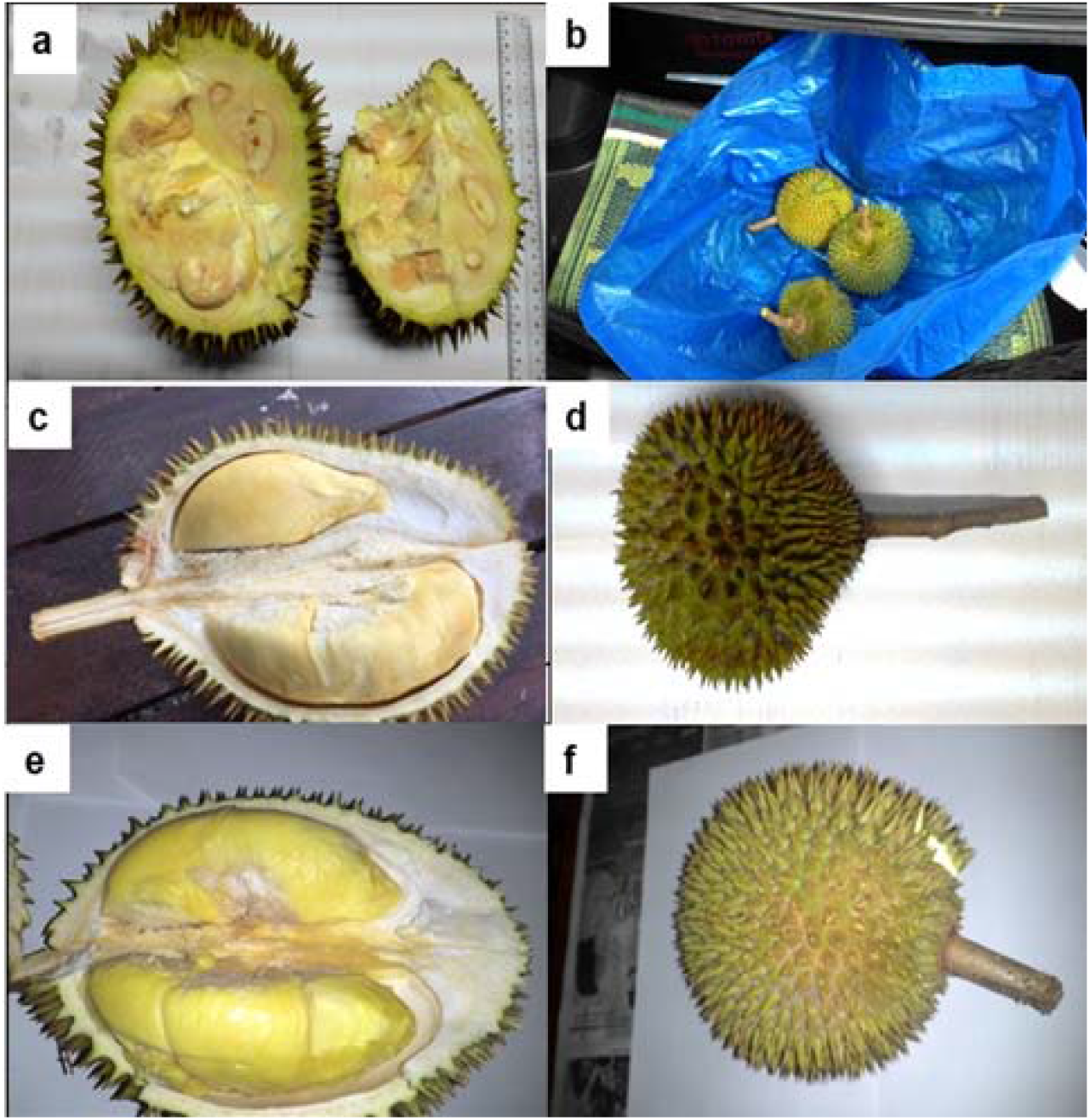
Durian fruits and pulp tissue. (a-b) Durian fruits at its young stage of growth and development (90 days); (c-d) Durian fruit at its mature stage of growth and development (120 days); and (e-f) Durian fruit at its ripening stage of growth and development (127 days).

### RNA isolation, cDNA library construction and HiSeq Illumina sequencing

An RNA extraction kit, namely the GeneAll RibospinTM Seed/Fruit RNA mini kit, was used to isolate the total RNA from young, mature, and ripened Durian pulp tissues. The RNA samples with RIN numbers more than 7.0 were considered for constructing the cDNA library as guidelines.

Poly-A mRNAs were purified from 1 ug of total RNA from respective samples using 20 μl of NEBNext Oligo d(T)_25_ beads as per illustrated in NEB Next Poly (A) mRNA magnetic isolation module. RNA mixtures were placed on a thermal cycler at 65 °C for 5 minutes and held at 4 °C to denature the RNA and facilitate binding the poly-A mRNA to the beads. The first and second strand of cDNA synthesis was performed with purified mRNA using the NEB Next Ultra RNA Library Prep Kit (Ilumina). Messenger RNAs (mRNAs) were eluted from the beads by adding the pre-prepared First-Strand Synthesis Reaction Buffer and Random Primer mix (2X). After incubation in a thermal cycler, second-strand cDNA synthesis was performed immediately by adding the mixture of second-strand synthesis reaction buffer and enzyme mix. The adaptor ligation was proceeded immediately by adding the TA ligase master mix and NEBNext adaptor. The final purification of the PCR reaction was carried out using AmPure XP beads. Before sequencing, the cDNA library’s quality was assessed using the Qubit 2.0 Fluorometer and Agilent Tape station. The anticipated data output from HiSeq Illumina sequencing was 16.7 million reads or 5 GB per sample.

### Raw read processing

After sequencing, the sequenced reads (raw reads) contain low-quality reads and adapters, which will affect the analysis quality. Therefore, it is crucial to filter the raw reads and get clean reads. Reads containing adapters, sequences with N less than 10 % (base cannot be determined), and sequences with (Qscore <= 5) base which is over 50 % of the total base, were filtered and removed. The quality of the raw data was assessed using the FastQC software. The FastQC results showed per tile sequence quality, base sequence content, sequence duplication level, overrepresented sequences, and adapter content. The pre-processing step was crucial to ensure the raw data was cleaned before downstream analysis. In addition, the removal of adapter contamination in sequence reads was a necessary step. FastQC (Andrews, 2010) and Trimmomatic (Bolger *et al*., 2014) tools were used to check and improve the quality of the raw transcriptome.

### Tools for the differential expression analysis

Four DEG/transcript identification methods were used (Figure S1). When the absolute expression log2 fold change was greater than 1.5, and the FDR-corrected p-value was less than 0.05, the analysis was considered statistically significant (Padovan *et al*., 2013; Aarhus, 2015; Zhu *et al*., 2017; Sebastian *et al*., 2018). A series of analyses, such as the number of expressed genes/transcripts, the number of up and down-regulated genes/transcripts, and the number of differentially expressed genes/transcripts, were determined.

#### CLC Genomic Workbench

Several QC trimming methods were used within the CLC Genomic workbench software before mapping to the reference *Durio zibethinus* genome. A reference-guided assembly was performed to assemble the sequencing reads by mapping the reads to the *D. zibethinus* reference genome with the defined mapping options for RNA seq, followed by the empirical analysis of differential gene expression (EDGE test), functional annotations, and categorisation of the assembled reads (Aarhus, 2015).

#### DESeq2

Principal component analysis (PCA) was performed with rlog-transformed transcripts count from htseq in DESEq2 to verify each library’s identity and suitability for differential expression (DE) analysis. The data analysis consists of pre-processing raw reads, mapping reads with HISAT2, transcript assembly using StringTie, gene counts with ht-seq, and differential analysis with DESeq2 (Love *et al*., 2014).

#### edgeR

MDS plot was constructed to assess the level of similarity between samples and their groups. This data analysis consists of pre-processing raw reads, mapping reads with HISAT2, transcript assembly using StringTie, and gene counts with htseq (Robinson *et al*., 2009; McCarthy *et al*., 2012).

#### CuffDiff

This data analysis consists of pre-processing raw reads, mapping reads with TopHat, transcript assembly using cufflinks, and differential expression analysis with CuffDiff. The up-regulated and down-regulated genes and transcripts were ranked and determined using the q value and fold change. FPKM value was used to evaluate the abundance of the genes in the differential group (Trapnell *et al*., 2012).

### Functional annotation of transcripts and pathway analysis

The Blast2GO 5 (Conesa and Götz, 2008) Basic software was installed on the computer. The differential group sequence in FASTA format was uploaded to the software. Then, the pathway analysis was conducted with the KEGG Blast2GO function. All genes that have a q-value less than 0.05 were added to the GO categories “biological process,” “molecular function,” and “cellular components.”

## Results

### Sequencing, annotation, and mapping to the Durian genome

The sequencing was carried out with the Illumina Hiseq 150 PE Run employing a commercial service provider. The data output was around 16.7 million reads per sample, equal to 5 GB. Total raw reads were 193,416,352 M, and filtered reads were 166,709,698 M, with a ≥ Q30 percentage. The quality of the raw data was assessed with the FastQC software. Trimmomatic and Fast QC were tools used to check and improve the quality of the raw transcriptome. In total, 110,351,584 M clean sequences were used for their downstream analysis. The high-throughput sequence data for nine samples were submitted to GEO (Gene Expression Omnibus), a public functional genomics data repository maintained by NCBI. Transcriptome datasets supporting this work’s conclusions are available through the NCBI under the accession number series GSE136290 (release date: March 31, 2023).

### Comparison of all mapping tools

Three different mapping methods, namely CLC Mapping, TopHat, and HISAT2 were used for mapping the reads to the Durian reference genome. The number of sequencing mapping reads using CLC is 49.82-59.42 %, TopHat 84.5-90.6 %, and HISAT2 88-95 %. Figure 2 shows the graphical representation of mapped and unmapped reads using three mapping tools. CLC mapping shows lower mapping rates, while TopHat and HISAT2 offer high and competitive mapping rates. Overall, HISAT2 outperformed TopHat significantly in aligning FASTQ reads to the Durian genome. TopHat and HISAT2 are a type of splice-aware aligners suitable for RNA reads. This spliced aligner can split reads at intron-exon boundaries. The high mapping rate could be due to the tools’ ability to handle the RNA sequences’ splicing sites. Also, HISAT2 is an improved TopHat version (Trapnell *et al*., 2009, Kate Shannon., 2016).

**Figure 2.**
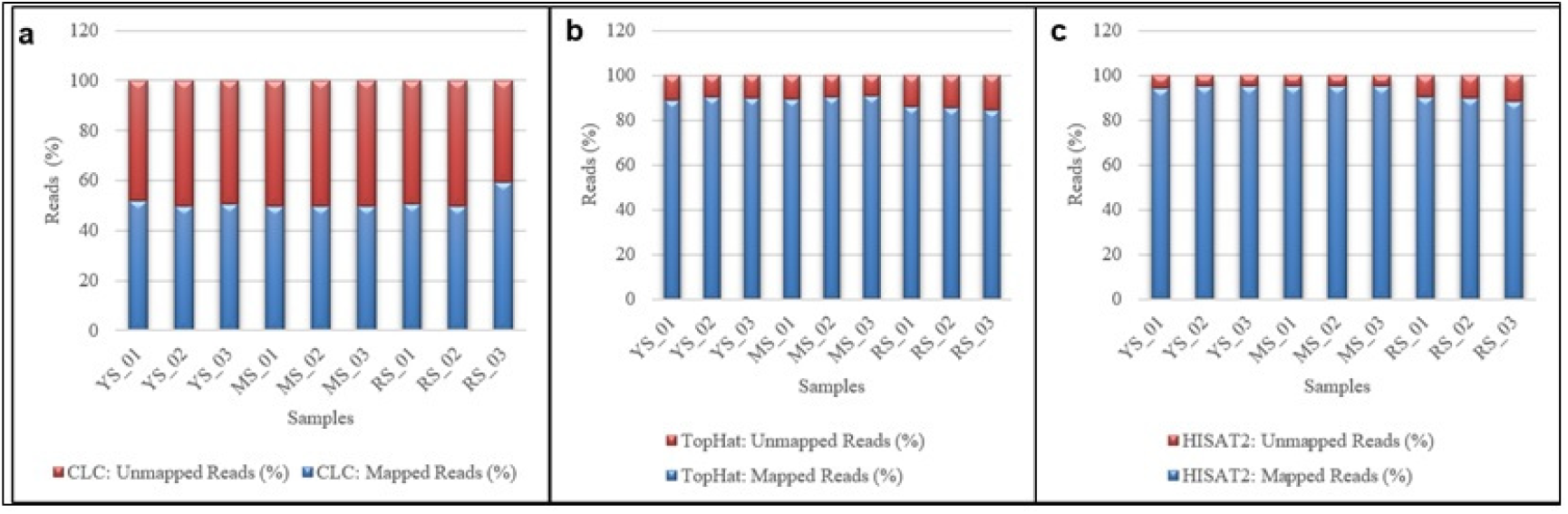
Comparison of the percentage of mapped and unmapped reads using CLC Mapping, TopHat, and HISAT2. (a) CLC Mapping, (b) TopHat Mapping, and (c) HISAT.

Compared with other research, the selection of mapping tools has not given any significant indicator to determine the correct mapping for the aligned reads. Raplee *et al*., 2019 stated that the generally low proportion of aligned reads for all input reads for HISAT2 and STAR is likely due to its quality. They also suggest that many input reads were poly(A) sequences, Illumina adapter sequences, and reads from libraries are too uninformative for accurate mapping to correspond to the very 3-end of mRNAs (Raplee *et al*., 2019). This study also contrasts with other previous studies, which reported a mapping method has a limited impact on the final analysis of DE genes (Costa-Silva *et al*., 2017). Table 1 shows the mapping tools’ performance in this study. In gene expression studies, the mapped reads’ output was used to compare the number of DEGs analyzed with four selected methods. It is also learned that selecting a suitable mapping tool relies on a variety of other variables, such as RNA quality, cDNA libraries, genome size, intron length, etc.

**Table 1.**
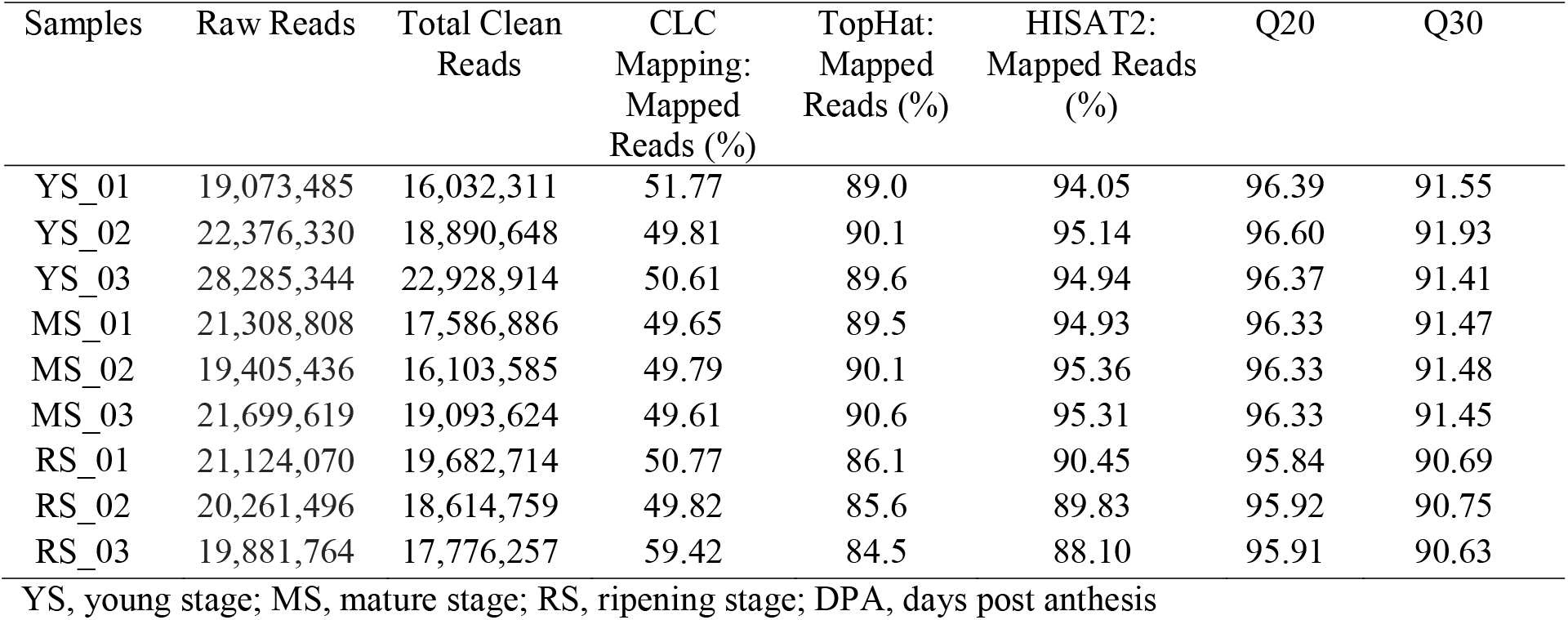
Statistics of the reads and the mapping percentage using CLC Mapping, TopHat, and HISAT2 in the present study.

### Principal component analysis (PCA) for differential expression groups

Each differential studied group was subjected to PCA analysis to identify whether the samples clustered within each group or with other groups. At the young stage (YS) vs mature stage (MS), the PCA results show that the YS and MS samples were clustered together in their group. The total variation of PC1 and PC2 for YSMS was 95 %. There was a strong correlation in their particular groups at the YS vs ripening stage (RS). The total variation of PC1 and PC2 for YSRS was 91 %. The total variation of PC1 and PC2 for MSRS was 100 %. The PCA for all samples in three different groups was calculated. Principal component 1 (PC1) and principal component 2 (PC2) were identified by variance stabilising transformation in all three groups. The percentage of variance indicates how much variance was explained by PC1 and PC2. These comparative principal component analyses demonstrate that biological replicates from all differential groups are highly correlated.

### Comparative transcriptome analysis and differential gene expression using merged DEGs

RNA-seq was used with the Illumina platform technology to explore dynamic changes in gene expression in durian fruit pulp tissues from three developmental stages, young, mature, and ripening. Various techniques are available to assess gene expression, each of which produces distinct results for rapid RNA-seq growth. However, there are no clear rules as to which technique yields the best accurate gene expression estimations. In this study, four statistical techniques were used to produce a more accurate list of DE genes. Other researchers used similar techniques to study the transcriptomic data (Rajkumar *et al*., 2015; Costa-Silva *et al*., 2017).

CLC was used to investigate the utilisation of commercial software for Durian RNA seq experiments in this study. CuffDiff was chosen in conjunction with Top Hat, as it is developed explicitly from transcripts, spliced regions, and promoters for DGE analysis. In the comparison, DESEq2 and edgeR were combined to investigate the gene-level expression, as this method work by integrating the raw reads using Ht-seq count. If the absolute value of the expression log2 fold changes > 1.5 and < −1.5, the differentially expressed genes and the transcript were considered statistically significant. Only those genes with expression identified as meaningful with q value of < 0.05 (5 % of a gene for each comparative group, false rate of discovery, FDR) were considered. After mapping the reads to the reference genome using one of CLC mapping, TopHat or HISAT2, transcripts were assembled in separate events using the CLC assembler, Cufflink, and StringTie. Table 2 highlighted the number of significant genes identified in up and down-regulation among four studied methods. Among four methods used for RNA - seq analysis, edgeR detected more DE genes in the durian transcriptome. This finding is similar to other researchers’ findings, as they reported that edgeR had identified more DEGs than other tools used (Fischer, 2003; Kvam *et al*., 2012; Seyednasrollah *et al*., 2013; Gallego Romero *et al*., 2014; Rajkumar *et al*., 2015).

**Table 2.** Total number of identified DEGs using four tools of CLC, CuffDiff, DESEq2, and edgeR.

| Differential Group | Expression | CLC | CuffDiff | DESeq2 | edgeR |
| --- | --- | --- | --- | --- | --- |
| YS/MS | Up-regulated | 2,565 | 1,461 | 5,790 | 5,117 |
|  | Down-regulated | 5,793 | 3,607 | 1,801 | 3,426 |
| YS/RS | Up-regulated | 1,611 | 1482 | 3,452 | 2,872 |
|  | Down-regulated | 3,296 | 2611 | 1,820 | 1,894 |
| MS/RS | Up-regulated | 4,141 | 2191 | 1,265 | 5,172 |
|  | Down-regulated | 2,570 | 926 | 3,705 | 5,870 |

YS, young stage; MS, mature stage; RS, ripening stage

### Differentially expressed genes in Durian during its fruit development

Genes are differentially regulated to bring about changes in durian fruit from a young to a mature stage. The differentially expressed genes were identified in the young and mature stages (Tables S1 and S2).

Pectinesterase 2-like (LOC111304600) had an 11-fold increase in expression, and the endoglucanase-like (LOC111278395) gene had a 10-fold rise in expression. Both of these genes are involved in cell wall metabolism. Glucan endo-1,3 beta-glucosidase 11-like (LOC111287487) and cellulose synthase-like protein G2 (LOC111280213), both implicated in the starch-sucrose breakdown, had a 9-fold and 6-fold expression, respectively.

The analysis suggested that the Polygalacturonase inhibitor-like (LOC111311552) gene was expressed highly (an 8-fold increase in expression) and is responsible for protein binding. Polygalacturonase inhibiting proteins (PGIPs) are cell wall proteins that inhibit the pectin-depolymerising activity of polygalacturonases and play an essential role in the plant defence system (Kalunke *et al*., 2015).

Five different genes, namely, LOB domain-containing protein 18 (LOC111304630), LOB domain-containing protein 42 (LOC111282412), LOB domain-containing protein 40 (LOC111290435), LOB domain-containing protein 1 (LOC111297059), and LOB domain-containing protein 19 (LOC111305909), were expressed during the early stage of durian fruit pulp development. These five genes are from the LOB (lateral organ boundaries)-domain gene family. Durian’s LOB gene family members exhibited a 9-fold increase in expression. Several studies have suggested that LBD genes have a broad range of functions (Luo *et al*., 2016). Genes from the plant-specific Lateral Organ Boundaries Domain (LBD) family encode transcriptional regulators with roles in various physiological and developmental processes in plants (Luo *et al*., 2016).

One of the MGL gene isoforms, namely methionine gamma-lyase-like (LOC111287425), responsible for the Volatile sulfur compounds (VSC) ‘s production, was up-regulated during the transition from the young to the mature stage. In a differential analysis of Durian, this gene is found up-regulated with a fold change of 10.163. This gene is involved in the transsulfuration metabolic pathway, where sulfur transfer from homocysteine to cysteine occurs. The pathway leads to the generation of several sulfur metabolites, including cysteine, GSH, and the gaseous signalling molecule hydrogen sulfide (H2S) (Sbodio *et al*., 2019).

ATP-binding cassette (ABC) transporters are essential for plant development in processes such as gametogenesis, seed development, seed germination, organ formation, secondary growth, protective layer formation, and phytohormone transport (Ha *et al*., 2017). In this study, four different ABC transporter gene families, namely, ABC transporter C family member 13 (LOC111274386), ABC transporter G family member 31 (LOC111301322), ABC transporter G family member 14 (LOC111286089), and ABC transporter G family member 11 (LOC111316978) were found significantly up-regulated during the early stage. Members of the ABC transporter gene family in Durian exhibited a 10-fold increase in expression.

### Differentially expressed genes in Durian during its fruit ripening

Genes that play an essential role in the Durian fruit pulps’ growth and development and are responsible for the transition from a mature to a ripening stage are depicted in Tables S3 and S4.

Four family members of xyloglucan endotransglucosylase (XTH) were present, namely, xyloglucan endotransglucosylase/ hydrolase protein 33 (LOC111317417), xyloglucan endotransglucosylase/ hydrolase 2-like (LOC111287953), xyloglucan endotransglucosylase/ hydrolase protein 6 (LOC111316166), and xyloglucan endotransglucosylase/ hydrolase protein 6 (LOC111292968). These XTH family members were directly annotated to the term cellular glucan metabolic process. The most significant increase in the expression during durian ripening was observed in this XTH gene family. Similar observations were also reported for the banana and papaya fruit softening process (Asif *et al*., 2014; Yao *et al*., 2014). Researchers’ previously reported gene families include expansins, pectate lyases, and xyloglucan endotransglycosylases responsible for softening banana fruit (Ha *et al*., 2017) Most of the genes involved in cell wall hydrolysis are also members of multigene families, and many have highly specialised functions in cell wall metabolism (Asif *et al*., 2014).

Three family members, namely, glucan endo-1,3 beta-glucosidase (LOC111285741), glucan endo-1,3 beta-glucosidase 12-like (LOC111275649), and glucan endo-1,3 beta-glucosidase (LOC111295531) were among the expressed transcripts. They are known for their hydrolysis activity (Hrmova and Fincher 2001; Singh *et al*., 2016). These genes and galactinol-sucrose galactosyltransferase 5 (LOC111316628), beta-amylase 3, and chloroplastic-like (LOC111305012) are also involved in sugar metabolism, and responsible for the degradation of starch to sugar.

Polygalacturonase (PG), LOC 111311521, was significantly expressed in Durian. It is responsible for fruit softening (García-Gago *et al*., 2009; Anand *et al*., 2018). Its high expression of 12.6-fold was noticeable while doing the data analysis. This gene is also involved in starch and sucrose metabolism. Other significant up-regulated genes involved in the hydrolysis activities were carboxylesterase SOBER1-like (LOC111275243), endoglucanase-like (LOC111278395), and beta-glucosidase 11 (LOC111305070). The research findings on *Carica papaya* L. var solo eight (8) fruits at the postharvest stage reported by Yao *et al*. (2014) reveal that the loss of the papaya’s firmness was positively related to the hydrolytic enzyme activities and the sweet taste of the presence of simple sugars such as galactose liberated from the polysaccharide complexes.

The data analysis suggests that Sulfate transporter 1.3, SULTR (LOC 111294357) is also activated during durian ripening. It is involved in sulfur metabolism (Leustek *et al*., 2000; Gigolashvili *et al*., 2014). Sulfur is required for the biosynthesis of proteins, co-enzymes, prosthetic groups, vitamins, amino acids like Cys and Met, GSH, and secondary metabolites such as GSL and sulfoflavonoids. Sulfate transporters are the most prominent S-metabolite transporters in plants, as sulfate is the primary source of sulphur taken from the soil. It is the most abundant S-containing metabolite in plant cells (Gigolashvili *et al*., 2014). The process is assisted by several sulfate transporter members (SULTR) (Pinsorn *et al*., 2018).

### Venn analysis of all differentials expressed genes

Venn analysis was carried out for all differential expressed genes to observe the specific and overlapped sequences among the three stages. Figure 3 shows the intersection among YSMS and YSRS and MSRS, with 67 up-regulated and 42 down-regulated genes overlapped in all their gene sets. The Venn diagram analysis revealed a total of 937, 504, and 1,666 of the specific differentials upregulated expressed genes, respectively, in the young, mature, and ripening stages of growth. The Venn diagram analysis showed a total of 1,965, 724, and 635 of the specific differentials downregulated expressed genes, respectively, in the young, mature, and ripening growth stage.

**Figure 3.**
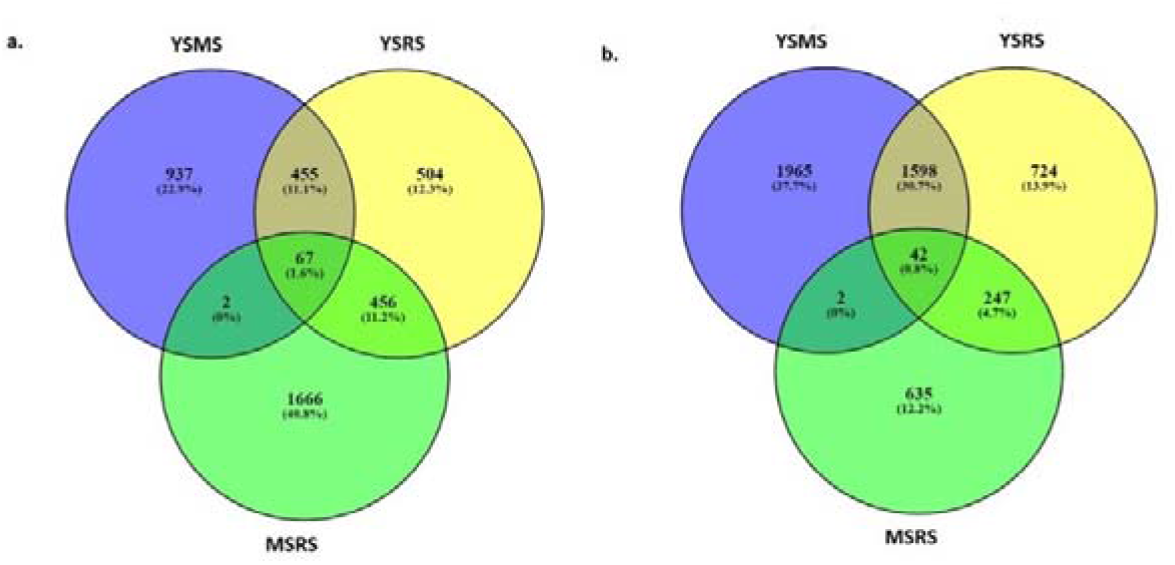
a. Venn diagram for upregulated differential expressed genes from three differential studies; namely YS/MS, YS/RS, and MS/RS. A total of 5,134 genes were expressed in all three stages. The number of tissue-specific genes are 1,965, 724, and 635, respectively. b. Venn diagram for upregulated differential expressed genes from three differential studies; namely YSMS, YSRS and MSRS. A total of 4,087 genes were expressed in all three stages. The number of tissue-specific genes are 937, 504, and 1,666, respectively.

### Functional analysis of differentially expressed genes

Blast2GO 5 Basic was downloaded to the desktop to identify the biological significance of each group’s transition stage. The entire DEGs from each group generated using CuffDiff were subjected to GO analysis to achieve a broader functional characterisation. The GO analysis was classified into three major categories, namely, biological process (BP), molecular function (MF), and cellular component (CC)). The total DEGs mapped to the three GO categories for YSMS, YSRS, and MSRS were 6,735, 3,898, and 7,310 (Table 3). All selected transcripts with q value > 0.05 were uploaded to Blast2GO for annotation to assign the putative functions to the DEGs. After running with InterProscan and Annex, the transcripts were assigned to the GO terms within three main categories (BP, MF, and CC).

**Table 3.**
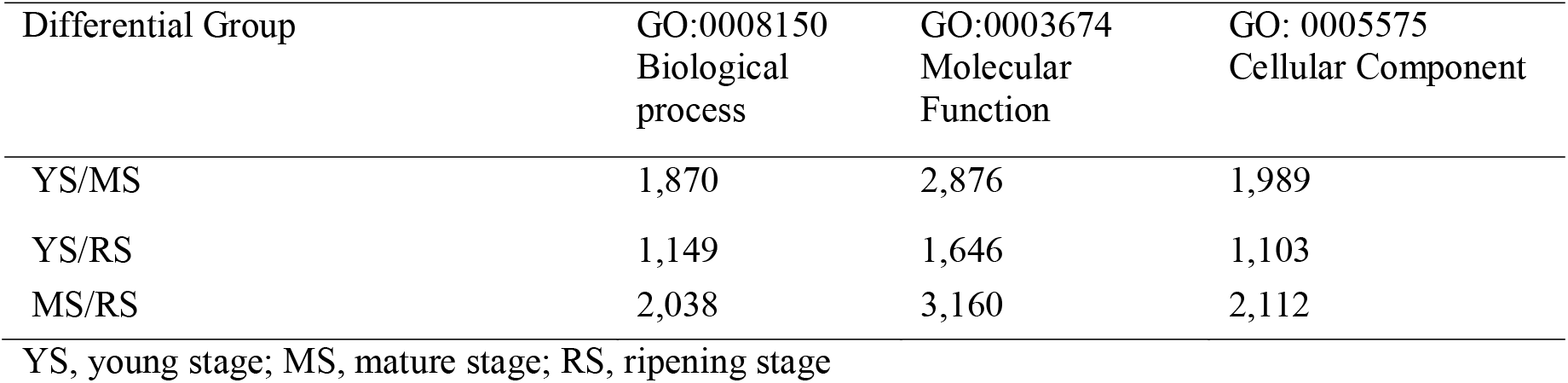
Analysis of GO terms for differently expressed sequences.

In developmental stage YS/MS, for biological process (BP), major sub-categories were 1,416 DEGs in the metabolic process (GO: 0008152), followed by 1,401 DEGs in the cellular process (GO: 0009987) and 395 DEGs in response to stimulus (GO: 0050896). As for molecular function (MF), binding (GO: 0005488) and catalytic activity (GO: 0003824) were the highest genes mapped to these GO terms, with a total number of 1,427 and 1,137, respectively. Cell (GO: 0005623), cell part (GO: 0044464), and organelle (GO: 0043226) with total DEGs of 1,297, 1,272, and 1,049 were enriched in cellular component (CC). In YS/RS, the major sub-categories for biological process (BP) were 862 for metabolic process, 806 for cellular process, and 227 for response to the stimulus. As for molecular function (MF), binding (GO: 0005488) and catalytic activity (GO: 0003824) were the genes mapped to these GO terms with a total number of 730 and 597, respectively. Cell (GO: 0005623), cell part GO: 0044464), and organelle (GO: 0043226) with total DEGs of 775, 746, and 533 were enriched in cellular component (CC). The number of DEGs for all GO terms was lower compared to YS/MS. In late-stage MS/RS, 1509, 1415, and 393 DEGs represented major sub-categories in biological process (BP) for metabolic process, cellular process, and response to the stimulus. As for molecular function (MF), binding (GO: 0005488), and catalytic activity (GO: 0003824) were the highest genes mapped to these GO terms, with a total number of 1,571 and 1,094, respectively. Cell (GO: 0005623), cell part (GO: 0044464), and organelle (GO: 0043226) with total DEGs of 1,354, 1,269, and 969 were enriched in cellular component (CC).

In the developmental stage of growth from young to mature (YS/MS), a total of 210 DEGs mapped to the developmental process (GO: 0032502) and 210 DEGs in anatomical structure development (GO: 0048856). Both GO terms were not present in the MS/RS. The DEGs are important for durian fruits’ growth and development. Several other GO terms of BP, MF, and CC that were not active in the developmental stage were localisation, catabolic process, and establishment of localisation, transport, non-membrane-bounded organelle, and intracellular non-membrane-bounded organelle. The highest DEGs of GO term associated with response to stress were in the ripening stage, MS/RS with 291, followed by YS/MS with 277 DEGs.

As shown in Figure 4, the developmental process, cellular component organization, anatomical structure development, protein modification process within the biological process, and RNA binding within the molecular function were the unique groups in YSMS. Localization, catabolic process, the establishment of localization and transport within the biological process, and non-membrane-bounded organelle, an intracellular non-membrane-bounded organelle within the cellular component, were the unique groups in MSRS. This finding revealed that the DEGs classified into the groups might be playing a tissue-specific role during fruit development (Table S5).

**Figure 4.**
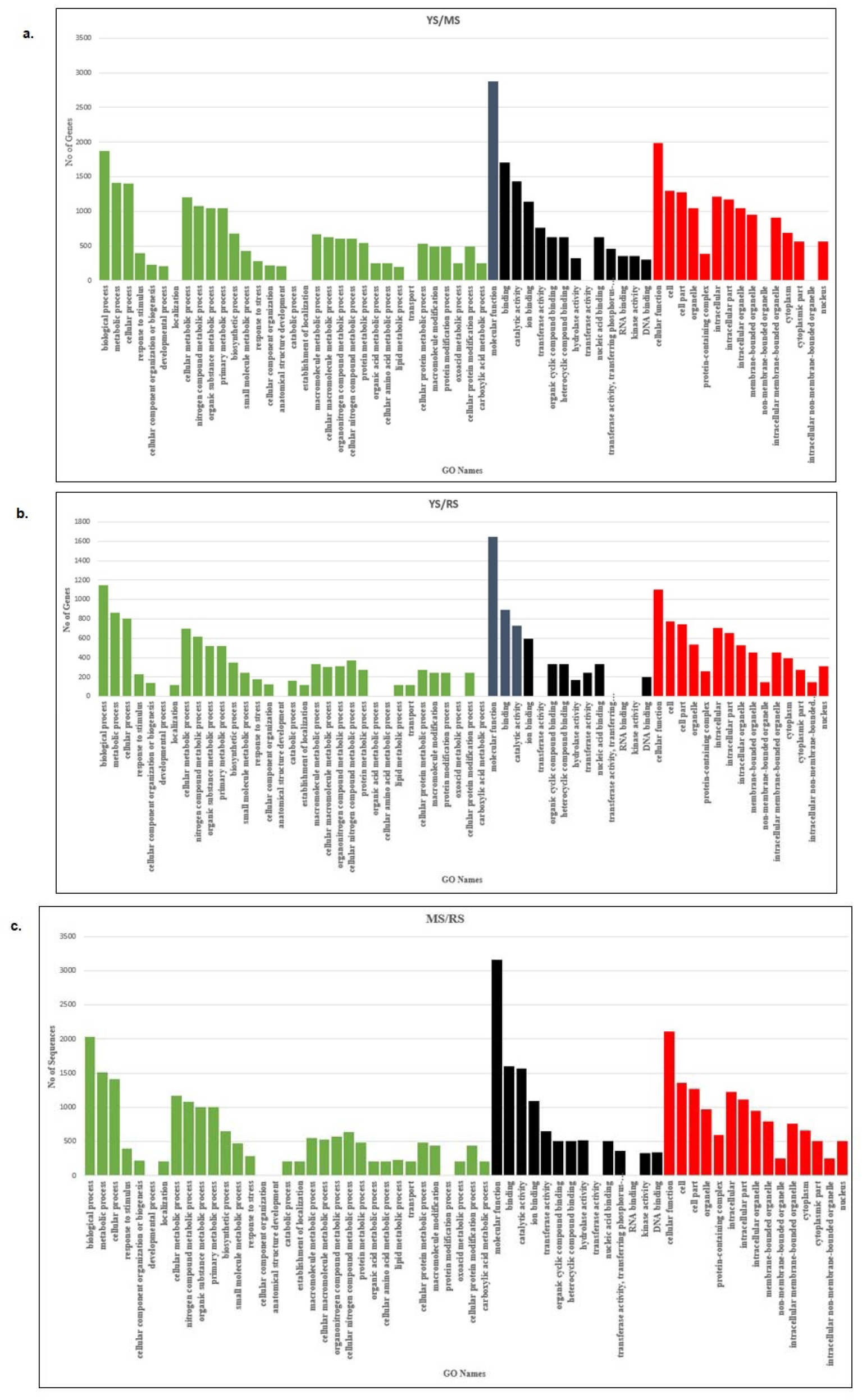
GO terms classification in a) YS/MS, b) YS/RS, and v) MS/RS

### Enrichment of the KEGG Pathway

Kyoto Encyclopedia of Genes and Genomes (KEGG) platform (https://www.genome.jp/kegg/) hosts various databases and provides tools for a systematic annotation and analysis of cell metabolic pathways and functions of the gene product. To further investigate the functional studies, all these up-regulated DEGs were enriched and annotated using the KEGG database to obtain the link to the Enzyme Commission number (EC). The EC was finally mapped to the relevant KEGG pathway to obtain the annotated pathway maps. The percentage of the mapped reads to the KEGG pathway for each differential group was 96.50 % (YS/MS), 60.20 % (YS/RS), and 35.30 % (MS/RS). Table 4 shows the number of the mapped pathway, the number of mapped sequences, and the EC number for each differential group. A total of 2,705 DEGs were assigned to 382 EC. The EC were subsequently grouped into 284 biochemical pathways.

**Table 4.**
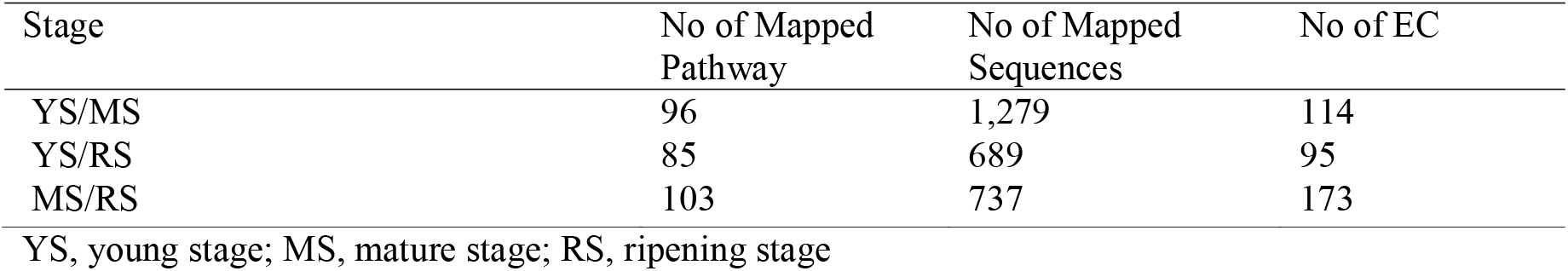
No of mapped pathways and sequences.

The top five KEGG class results from the young to mature stage data analysis indicate that most transcript encoding enzymes were linked to carbohydrate metabolism (254), metabolism of cofactors and vitamins (200), amino acid metabolism (176), nucleotide metabolism (142) and biosynthesis of other secondary metabolites (82). Carbohydrates are highly essential due to its contribution to texture, flavour, color, and nutritional value in horticultural commodities (Yahia *et al*., 2019). The classification suggests that the highest number of enzyme activities in Durian growth in the young stage were related to carbohydrate metabolism. This indicates that many pathways related to carbohydrate metabolism were activated to support the durian fruit’s growth and development. Understanding the gene expression patterns in carbohydrate metabolism will facilitate the investigation of durian fruit quality. Durian fruits development involves the accumulation of starch and sucrose. According to KEGG, 254 genes were associated with carbohydrate metabolism, and these genes were classified into several metabolic pathways. The highest number of expressed genes during the early fruit development were from starch and sucrose pathways. Starch is present in unripe durian, mainly in its pulp, which is transformed into sugars during the ripening process. These sugars contribute to the level of sweetness of the fruits. In the transition from young to ripening stage, GO analysis produced 4,154 sequences mapped with GO terms, and 2,248 transcripts encoding enzymes were annotated.

The top five KEGG class results showed that the expressed genes during the transition from young to ripening stage were mainly for metabolism of cofactors and vitamins (109), carbohydrate metabolism (102), lipid metabolism (77), nucleotide metabolism (75), and amino acid metabolism (65). The pathways for nitrogen and glutathione metabolism contained the greatest number of expressed genes.

During the transition from mature to ripening stage, GO analysis produced a total of 7,629 sequences mapped with GO terms, and of these 4,096 transcripts encoding enzymes were annotated. The top five KEGG class results were the metabolism of cofactors and vitamins (200), nucleotide metabolism (192), carbohydrate metabolism (85), lipid metabolism (61), and amino acid metabolism (56). During the late stage of fruit development, the sulphur metabolism pathway had the highest number of expressed genes, indicating the durian’s strong odour.

In contrast, the classification of transcripts suggests that the highest number of metabolism of cofactors and vitamins outperformed the other pathways in the late stage of Durian growth. In KEGG, 180 genes were identified to be associated with thiamine metabolism to support the nutritional quality and health benefit of durian fruit, mainly in the energy metabolism (Figure 5 and Table S6) (Lonsdale, 2006).

**Figure 5.**
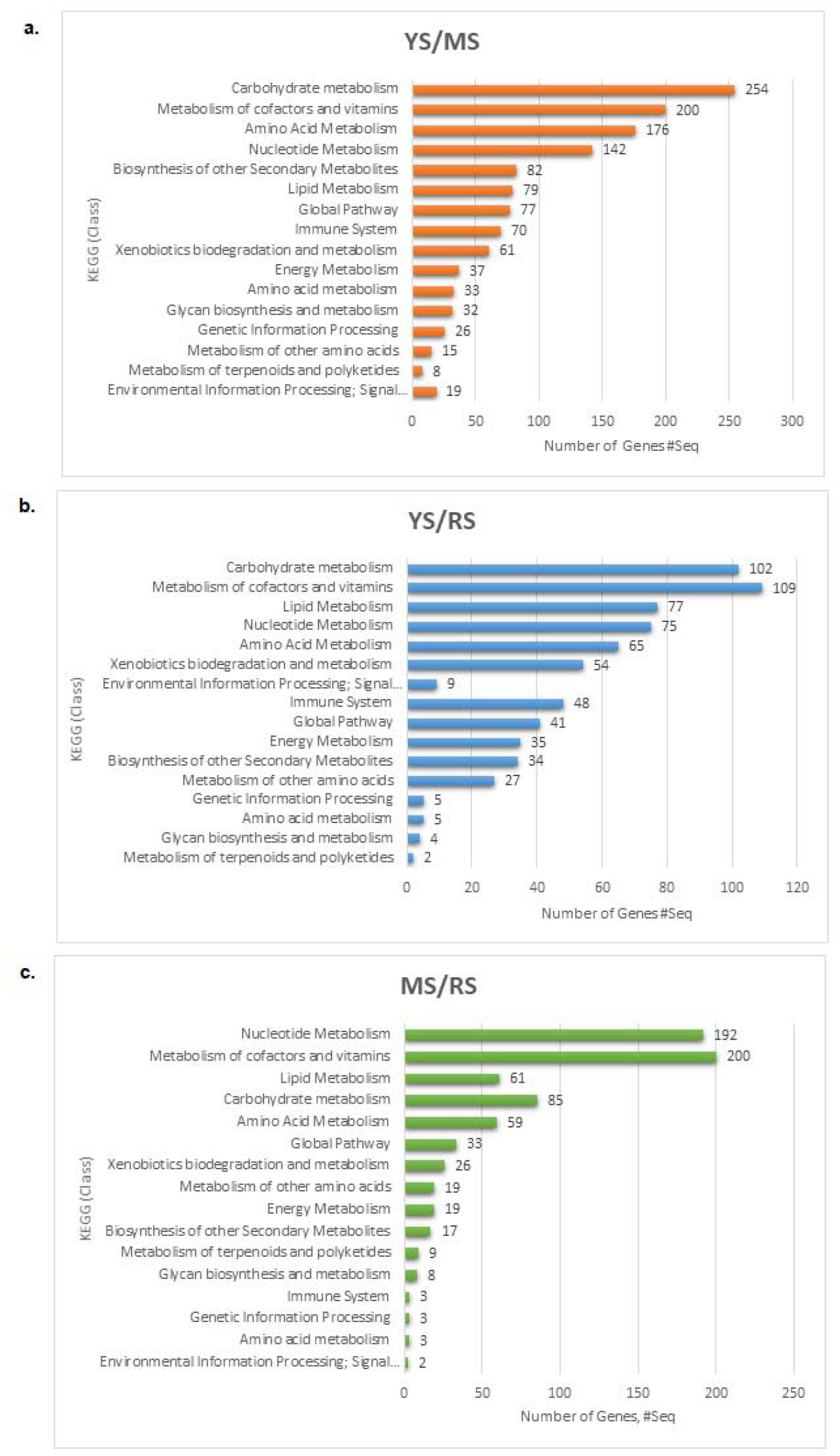
Classification of the sub-categories of KEGG pathways in a) YSMS, b) YSRS, and c) MSRS

### Genes involved in Durian fruit softening

During the fruit ripening developmental stages, a set of distinct softening genes are expressed. The significant clusters, young vs. mature, and mature vs. ripening annotation, are shown in Figure 6 and Table S7. In developmental stages, several genes that showed more than a 2-fold change in expression level were pectinesterase 3-like, U-box domain-containing protein, expansin-like B1, and probable xyloglucan. These genes coordinate to soften durian fruit in the transition from the young to the mature stage.

**Figure 6.**
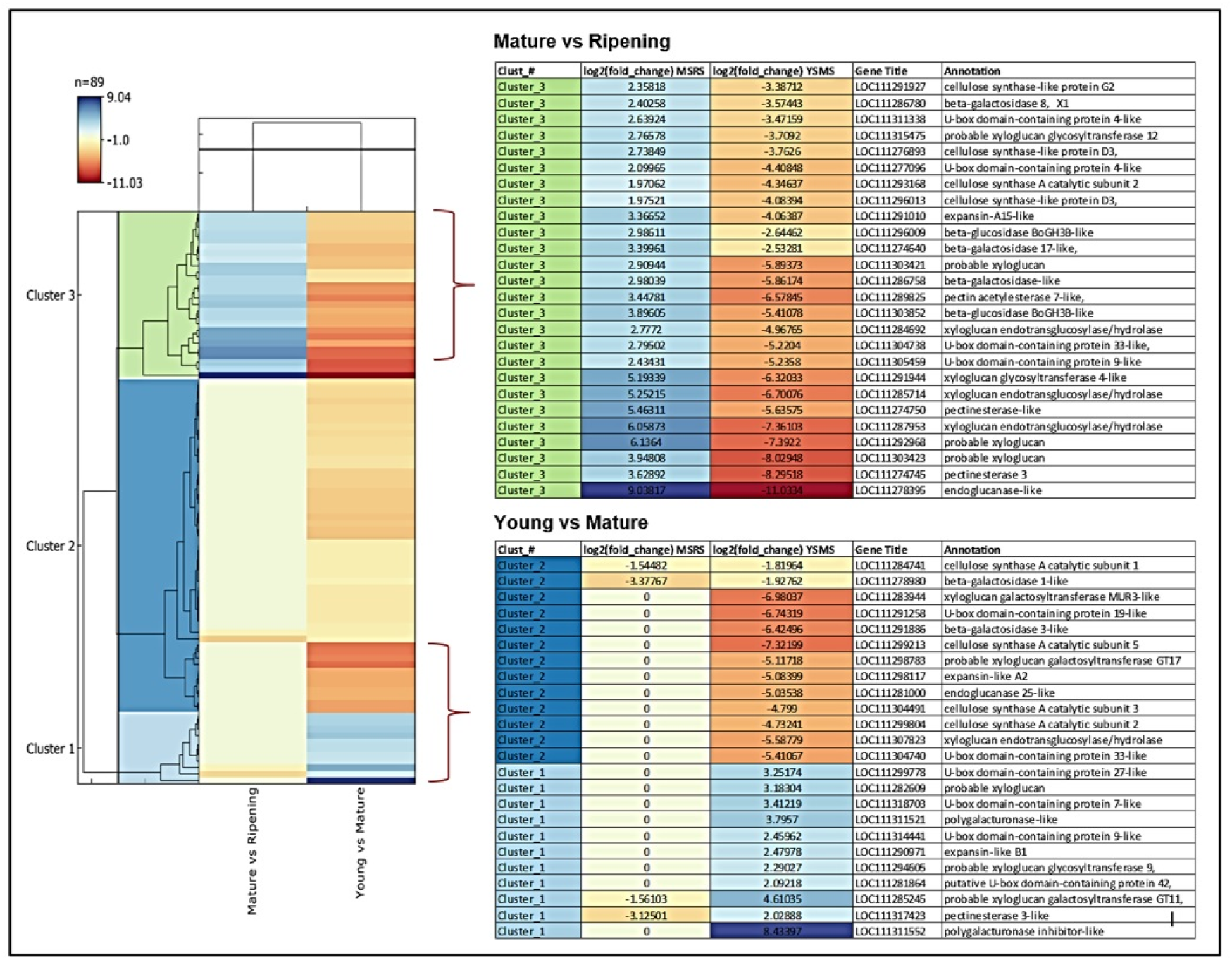
Hierarchical clustering analysis, heatmap, and key regulatory genes involved in Durian fruit softening development and ripening stage. Blue and orange bands indicate the high and low expression of genes, respectively. The color scale representing log2 fold change values is shown; young to mature stage (YSMS), young to ripening stage (YSRS), and mature to ripening stage (MSRS).

Polygalacturonase inhibitors showed the highest expression of 8-fold in the developmental stage. This plant protein can inhibit polygalacturonase (PG) enzymes in the young to mature durian growth stage. In contrast, this gene is down-regulated in the ripening stage with −8-fold expression to activate durian fruit pulp softening genes.

It is known that expansins A affect preferentially xyloglucan-rich type-I cell walls characteristic of dicotyledonous plants (Sharova, 2007; Marowa *et al*., 2016). In contrast, expansins B modify type-II cell walls rich in arabinoxylans and β-glucans (Sharova, 2007). Only proteins from the alpha-expansin and beta-expansin families have been found to weaken cell walls. The roles of expansin-like A and expansin-like B, on the other hand, remain unknown. (Palapol *et al*., 2015). While doing data analysis, we found that the expansin-B is expressed in Durian with more than a 2-fold change in expression level, indicating its role in fruit pulp softening. Endoglucanase also showed a high level (9-fold) of change in its expression.

Other softening genes that showed more than 2-fold expression change in the ripening stage were pectinesterase, U-box domain, xyloglucan endotransglucosylase/hydrolase, beta-galactosidase, cellulose synthase, endonuclease 1, expansin-A (also called α-expansins), beta-glucosidase, and pectin. It appears that more genes were expressed during the ripening stage than the developmental stage of durian fruits.

### Ethylene synthesis and signal transduction pathway in Durian

The molecular level of ethylene-controlled ripening in Durian has not been investigated in detail. Most of the studies conducted refer to a single gene or a single family of genes. The genes involved in ethylene biosynthesis and signal transduction are significantly important in the growth and development of Durian fruit. Ethylene biosynthesis is essential for the ripening of fruits; hence, the data were analysed to study the genes involved in this process. Durian shows a burst of respiration and a gaseous hormone ethylene biosynthesis at the beginning of its ripening, which regulates the fruit-maturing aspects.

The genes involved in ethylene biosynthesis, signalling pathway, and regulatory response process were identified from the transcriptome data (See Table S8). S-adenosyl-L-methionine (SAM) synthase converts methionine to SAM with a 2-fold expression during fruit ripening (Bouzayen *et al*., 2010). Several genes linked with ethylene biosynthesis have been identified with six ACS gene members. The ACS genes are involved in converting SAM to 1-aminocyclopropane-1-carboxylic acid (ACC) with 2-fold expression in both stages. Most 1-Aminocyclopropane-1-Carboxylic Acid Oxidase (ACO) genes with high expression of 4-fold increase were identified at the stage of ripening. This ACC oxidase is known to play a role in converting ACC to ethylene (Bouzayen *et al*., 2010).

In addition to this, an array of genes that are associated with ethylene signal transduction have been identified in this study. Fifteen selected gene family members of mitogen-activated protein kinase (MAPK) and genes linked to transcription factors such as EIN3, ethylene-like receptor, protein kinase constitutive triple response1 (CTR)-like were identified and grouped under the ethylene signalling pathway (Binder, 2020). The ethylene signal transduction pathway has been extensively studied in fruits as ethylene affects fruit’s post-harvest physiology and storage requirements (Binder, 2020). The ethylene response factors (ERFs) that act downstream are the last component of the ethylene signalling pathway to regulate ethylene-responsive gene’s expression. ERFs have been reported to play essential roles in plant development, flower abscission, fruit ripening, and defence responses (Gao *et al*., 2020). Fifty-six (56) ERF members expressed the ethylene signalling pathway to regulate the ethylene response for the developmental and ripening processes. ERFs mediate the ethylene-dependent ripening gene responsible for Durian fruit’s significant texture, flavour, and taste (Binder, 2020; Thirugnanasambantham *et al*., 2015). We have presented the ethylene synthesis and signal transduction pathway for a simplified explanation purpose (Figure 7).

**Figure 7.**
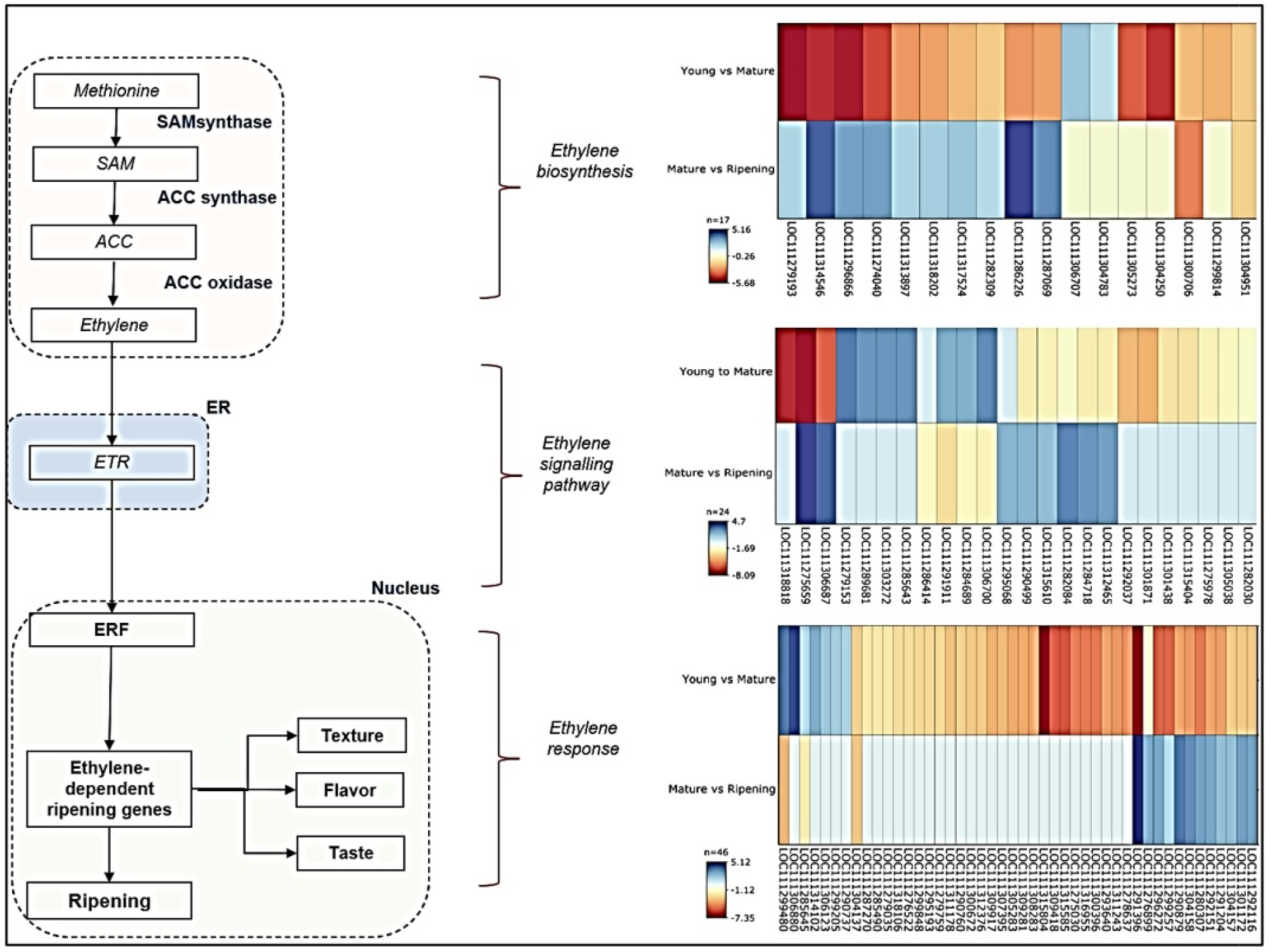
Selected members of gene families involved in ethylene biosynthesis and signal transduction in durian fruit during its developmental and ripening stages. The colour scale is representing the log2 fold change values.

### Role of MGL genes and VSC production in Durian’

To study the biological process related to Durian fruit’s strong aroma, the heatmap analysis between isoforms of MGL (methionine gamma-lyase-like) gene responsible for volatile sulphur compounds (VSC) production was carried out. The VSCs have been identified as important factors determining the degree of aroma and taste of Durian fruit pulp. MGL enzymes break methionine and cysteine into methanethiol, ethanethiol, and ethionine (Fischer and Steinhaus, 2019).

MGL genes produce VSC. Durian fruit is unique in the sense of its aroma conferred by VSCs. A study on the Durian genome reveals the presence of four copies of MGL, which contributes to making Durian fruit pulp aroma strong (Teh *et al*., 2017). There are two main groups of volatile compounds; sulfur-containing volatiles, such as thiols, disulfides, trisulfides, and esters associated with a fruity aroma. MGL genes control the VSCs level. A distinctive sulfuric aroma can be smelled in durian fruit pulp due to VSCs produced by the MGL enzyme (Teh *et al*., 2017). Seven low molecular weight sulfurs containing compounds are reported in durian fruit pulp, including hydrogen sulphide, methanethiol, ethanethiol, and propane-1-thiol (Cannon and Ho, 2018).

While annotating transcripts, we noticed that Durian fruit pulp contains five different MGL isoforms expressed in three different stages of growth, YS, MS, and RS. One outlier of the MGL gene, *MGLc* (LOC111287425), is found in the early stage of Durian fruit pulp.

Heatmap showing the gene abundance of MGL isoforms (FPKM) in YS, MS, and RS is depicted in Figure 8. This heatmap was generated using StringTie. The abundance of each reference gene was estimated with StringTie. The output of FPKM (Fragment per kilobase) and TPM (Transcript per kilo base million) value were calculated based on the normalisation of the sequencing depth and gene length. The highest FPKM or TPM value was considered the most abundant genes for the particular sample. FPKM and TPM values were used to compare the relative gene expression levels within a sample.

**Figure 8.**
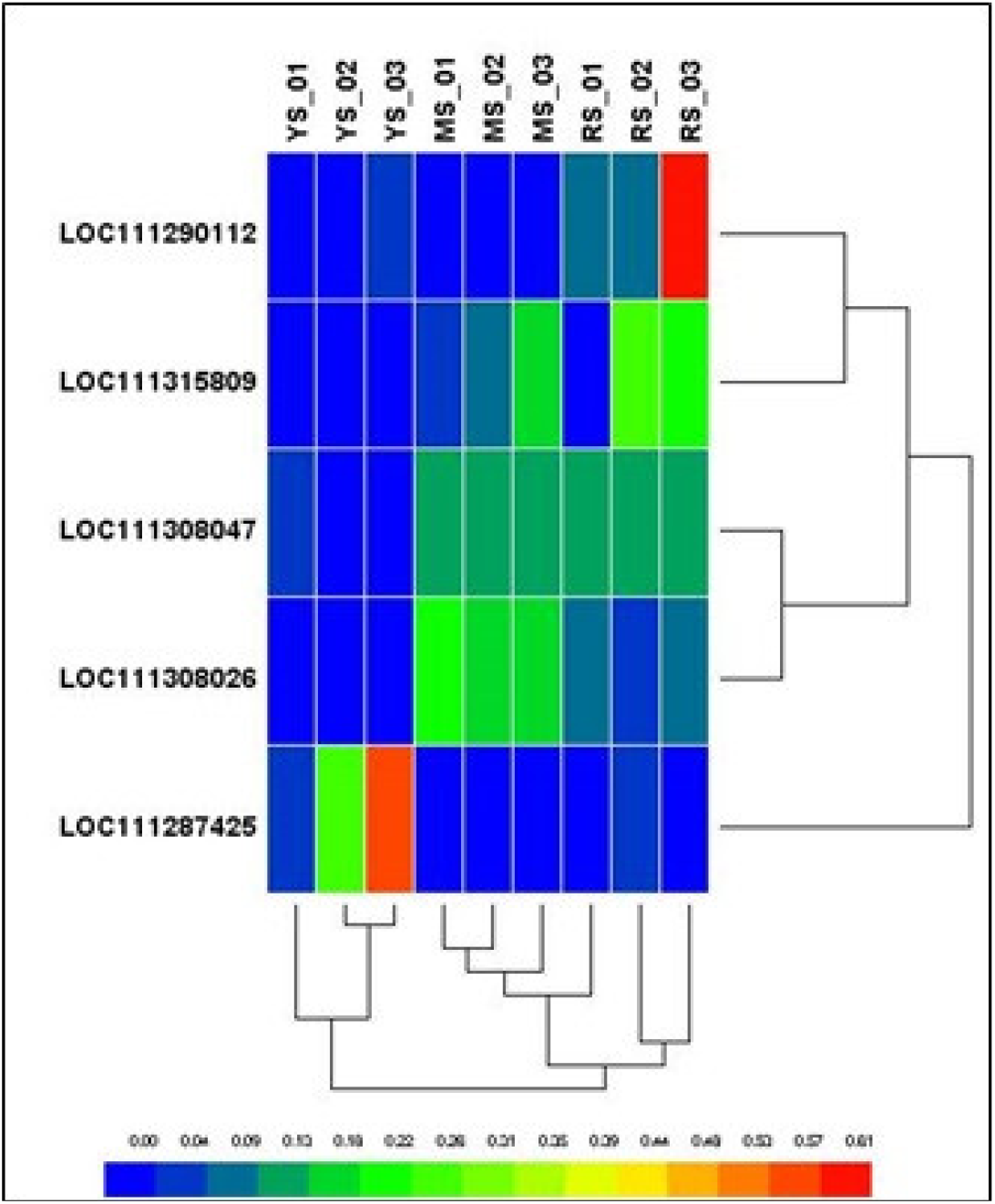
Heatmap showing the expression of MGL (Methionine gamma lyases) gene isoforms in three different growth stages of durian fruit pulp. YS represents for young stage; MS represents for the mature stage, and RS represents for ripening stage. The FPKM value and Pearson’s distance are used as the metric.

We found that durian fruits possess various isoforms of the MGL genes. The MGL gene isoforms are abundantly expressed. The transcripts were found in young, mature, and ripening fruit pulp samples. In young, mature, and ripened fruit pulp samples, *Durio zibethinus* methionine gamma-lyase-like (LOC111287425), transcript variant X1, mRNA, *D. zibethinus* methionine gamma-lyase-like (LOC111308026), transcript variant X3, misc_RNA, and *D. zibethinus* methionine gamma-lyase-like (LOC111290112), partial mRNA, were found, respectively. MGL genes abundantly expressed in the mature and ripening stage were *D. zibethinus* methionine gamma-lyase-like (LOC111308047), mRNA, and *D. zibethinus* methionine gamma-lyase-like (LOC111315809), mRNA. The correlation matrix was performed to compare the five MGL gene copies to observe its expression value in three developmental stages. There were two major clusters, major cluster 1 (C1) comprised of 2 minor clusters (C1a and C1b). Major cluster 2 comprised of C2, the outlier. C1a showed a relationship between *MGLa1* and *MGLa2*. Both MGL genes in cluster C1a showed a varying high level of expression value in the ripening stage. While C1b comprises MGLb1 and MGLb2, it showed a similar expression pattern from the mature to ripening stage. MGL genes were up-regulated from the mature to ripening stage. There were two transcripts under *MGLb2;* namely XM_022906389.1 and XM_022906396.1. Major cluster C2 was an outlier, as this MGL gene, *MGLc*, showed an uneven expression level only in the young stage of growth. This gene was downregulated in the mature to ripening stage. Under this gene, there were three transcripts; XM_022877994.1, XM_022878003.1 and XM_022877983.1. Overall, the highest-level expression value of the MGL gene was expressed in *MGLb1* in the transition stage from mature to ripening.

The structural analysis of MGL proteins published in the 2018 International Conference on Tropical Fruit Pests and Diseases suggests that all four MGL genes in Durian have 89 % to 93 % similarity at the protein sequence level. Phylogenetic analysis demonstrated that the MGL genes in Durian are evolutionary-related with high genetic diversity, separating MGLb_1 into different clades (Yusof, 2018). The active sites in MGL protein sequences are highly conserved. However, the regions proximal to the active sites in MGLb_1 showed more residue substitutions than others. Their sequence analysis also showed that LOC111315809, MGLb_1 (accession number XP_022773569) contained more residue substitutions than other MGL genes. This gene expression was highly abundant in the ripening stage, suggesting that it produces more VSCs than other isoforms.

Furthermore, they also suggested some possible residue substitutions in MGLb_1 that may affect the activity of MGL in the recognition and elimination of methionine, which eventually contributes to the pungent odour of ripening Durian fruit (Yusof, 2018). Our research findings were similar. The *MGLa2 (*LOC111315809) was expressed abundantly in the ripening fruit pulp tissue samples. These findings explain why the strong aroma of Durian is felt when fruit reaches the ripening stage.

## Discussion

This paper describes the use of next-generation sequencing (Hi-Seq Platform) to understand the molecular attributes of Durian fruit in its young, mature, and ripen stages. A better understanding of how these processes are genetically regulated will facilitate the potential capacities and development of effective strategies to improve local durian fruits by identifying key regulatory genes and their manipulation using genetic engineering tools (Husin *et al*., 2018).

We have used three different mapping tools, CLC (Aarhaus, 2015), Top Hat (Trapnell *et al*., 2009), and HISAT2 (Kim *et al*., 2015), to map the reads against the Durian Reference Genome. A comparison of these mapping tools showed that HISAT2 is a highly sensitive approach to mapping transcriptome data to the Durian Reference Genome, with the highest mapping rate of 95 % (Sutthacharoenthad, 2019). Differential expression analysis has revealed a different number of Durian DEGs for all used tools (CLC, CuffDiff, DESEq2, and edgeR) specific to their level of developmental stages in differential expression analysis. The reason is mainly due to the different statistical methods unique to each tool, and the number of mapped reads specific to the tools used is not determined by high or low mapping rates. We have learned from our results that higher rates of mapping do not indicate a higher DEG identification. Venn analysis was carried out further to find the overlapped DEGs identified by four different tools used. As a result, a comparison of the top 50 highly significant genes revealed the similarities and dissimilarities between these four pipelines. Therefore, it is advantageous to combine the results of different tools to obtain a more detailed and reliable DEGs results output.

Previous studies published by other research groups also support selecting top candidate genes to explore further using several tools for identifying DEGs (Costa-Silva *et al*., 2017; Raplee *et al*., 2019). From Schurch *et al*., 2016, it is recommended to use edgeR for samples of less than 12 replicates due to the superior combination of a truly positive and false positive. DESeq2 is recommended to use for higher replicates of samples. Only DESeq2 and edgeR (TMM) can maintain a reasonable false-positive rate in the presence of top count genes without any loss of power.

Results suggest that the total number of identified up and down-regulated expressed genes in all groups is higher using DESeq2 and edgeR compared with CLC and CuffDiff. The highest range of overlapping gene similarities between DESEq2 and edgeR is 72.1-85.4 %. Unknown genes and functions dominate most of the top DEGs associated with DESeq2 and edgeR. These obscure results need further research work using both tools in the identification of novel genes.

DESeq2 and edgeR tools identified the highest number of up-regulated genes compared to other tools in the transition of young to mature and mature to ripening stage. The edgeR has successfully identified the highest number of DEGs in up and down-regulated genes in the transition from mature to ripening compared with other tools. For comparative functional analysis and validation between four tools, the top 50 DEGs in up and down-regulated were selected from each tool. The function of the top significant genes between stages shows almost similar patterns. The most common up-regulated genes were involved in cell wall development, cell membrane and cytoplasm, and defence activation in early fruit growth. In the late stage of growth, the most common up-regulated genes were involved in the critical process of growth and development, regulated organ development, stress, defence against pathogen attack, and disease resistance.

Transcriptome analysis also revealed that genes associated with cell wall softening, such as polygalacturonase, pectin, etc., were significantly up-regulated at the ripening stage of 127 DPA. To delay durian fruit pulp softening, inhibiting PG activity was found in 90 DPA of the Durian’s developmental stage. The essential genes expressed in the ethylene biosynthesis system responsible for Durian ripening were also identified. Interestingly, the transcriptome data analysis suggests that there are five isoforms of the MGL gene, and these isoforms are expressed in Durian fruit pulp tissues during developmental and ripening stages. The MGL and ACS are essential genes. Sugar is one of the essential biochemical components that determine fruit quality. A series of enzymes control starch and sucrose metabolism during durian fruit development and maturation (See Table S9).

The Jackfruit (*Artocarpus heterophyllus* Lam.) is closely related to Durian fruit. Researchers working on Jackfruit have examined how its perianth transcriptomes and metabolites are related to sugar metabolism. They found seventeen (17) sugar metabolic genes involved in the starch and sucrose metabolism pathway in ripened Jackfruit (Hu *et al*., 2016). Some of the genes identified in Jackfruit are also expressed in Durian fruit pulp tissue. The commonly expressed genes, namely 1,4-alpha-glucan-branching enzyme 2-1, chloroplastic/amyloplastic-like, fructokinase-like 1, chloroplastic, and hexokinase-3-like, are expressed during the transition from young to mature stage. Similarly, during the transition from the mature to ripening stage, genes, namely hexokinase-3-like, sucrose-phosphatase 2-like, probable sucrose-phosphate synthase 1, glucan endo-1,3-beta-D-glucosidase-like, and fructokinase-like 1, chloroplastic are also expressed in Jackfruit (Hu *et al*., 2016).

As a whole, this study has successfully provided the transcriptome data and gene expression data specific to young, mature, and ripened developmental stages of Durian fruit pulp. The research findings reported in this paper will serve as preliminary genomics expression data, which may be helpful in developing a new strategy for genetic engineering to create a new Durian variety with desirable traits beneficial for consumers and Durian growers.

## Acknowledgement

The authors want to acknowledge that the research funding for this project was provided by AIMST University, Malaysia, under its internal research grant scheme. The authors also want to thank the Agriculture Park of Universiti Putra Malaysia (UPM), Selangor, Malaysia, for providing the Durian (clone D24) fruits samples for the study.

## Authors Contributions

Nurul Arneida Husin (NAH) collected the fruit samples, carried out the laboratory work, performed bioinformatics data analysis, validated the data analysed, and prepared the manuscript draft. SJB conceived the idea and developed the project proposal and experimental strategies; he reviewed and edited the manuscript to finalise it. SJB, SR, and RK served as the main supervisor, field-supervisor, and co-supervisor for NAH, respectively. SR and RK provided their input for manuscript improvement.

## Supplementary material - the following online material is available for this article

**Figure S1.**
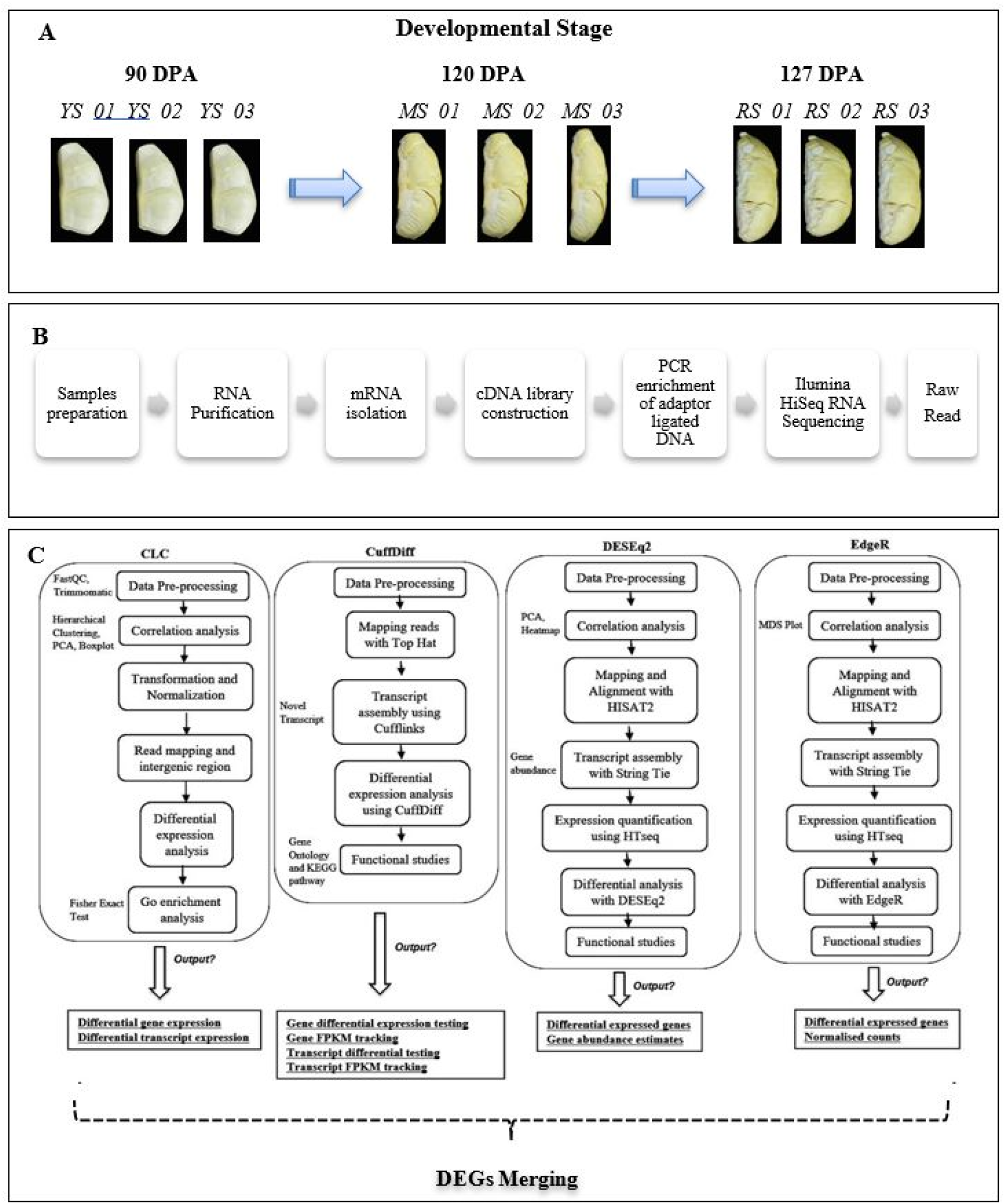
Experimental design of the study. (A) The nine samples, a biological triplicate of 90 days post-anthesis (DPA), 120 DPA, and 127 DPA, were used in the transcriptome study. (B) Wet lab experimental workflow, and (C) overview of transcriptome analysis approach using the combined methods.

**Figure S2.**
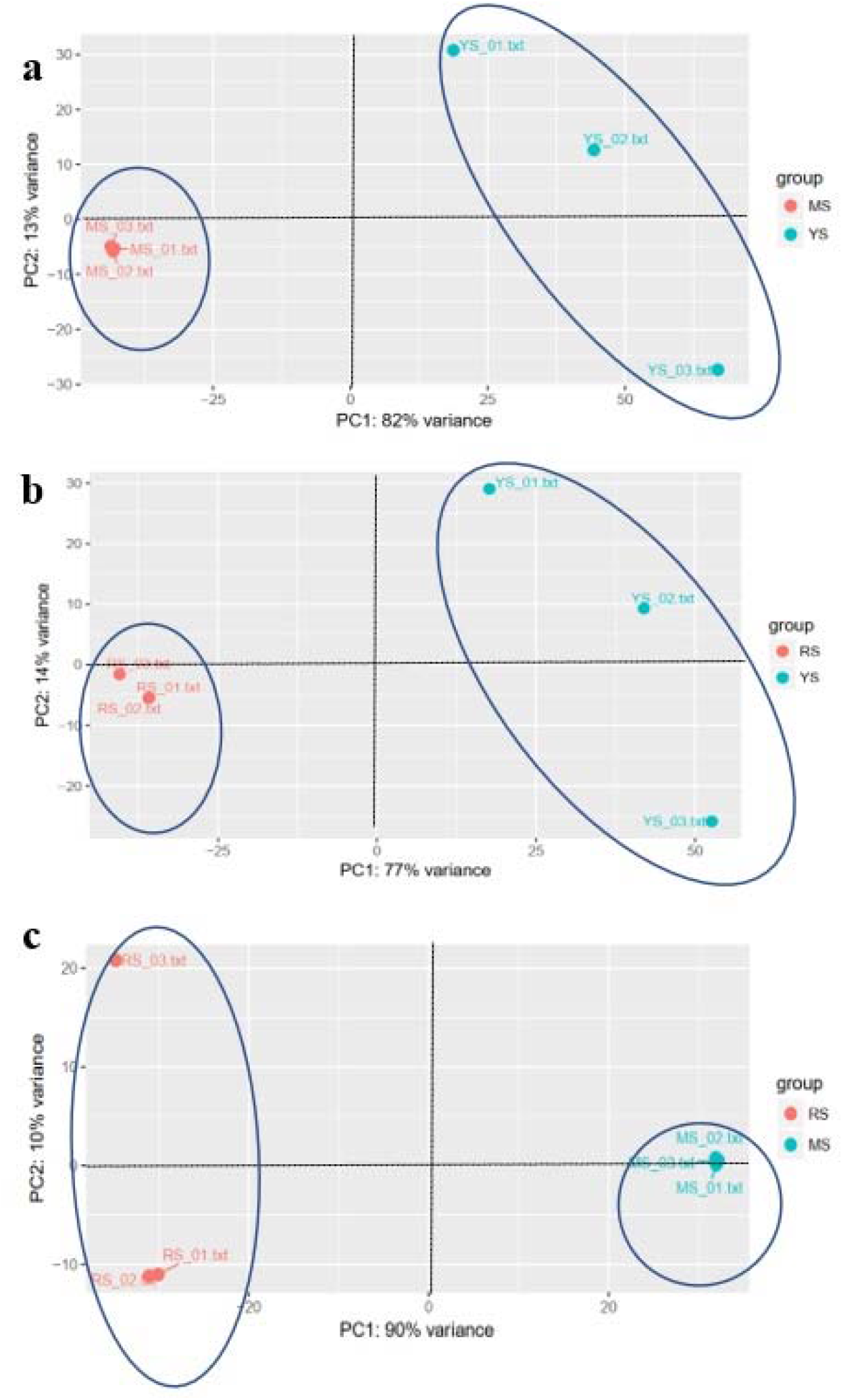
The comparative Principal component analysis (PCA) plot of the samples at young, mature, and ripening stages. Each replicate was plotted as an individual data point. This type of plot was useful for visualising the overall effect of experimental covariates and batch effects. The percentage of variance indicates how much variance was determined by PC1 and PC2. (a: Young Stage vs Mature Stage (YS/MS), b: Young Stage vs Ripening Stage (YS/RS), c: Mature Stage vs Ripening Stage (MS/RS).

**Table S1** - Differentially up-regulated expressed genes between young stage and mature stage of durian fruit pulp. We used FDR: <0.05, Log fold change >1.5 and <-1.5.

**Table S2** - Differentially down-regulated expressed genes between young stage and mature stage of durian fruit pulp. We used FDR: <0.05, Log fold change >1.5 and <-1.5.

**Table S3** - Differentially up-regulated expressed genes between mature stage and ripening stage of durian fruit pulp. We used FDR: <0.05, Log fold change >1.5 and <-1.5.

**Table S4** - Differentially down-regulated expressed genes between mature stage and ripening stage of durian fruit pulp. We used FDR: <0.05, Log fold change >1.5 and <-1.5.

**Table S5** - Categorisation of expressed genes to Gene Ontology terms in YS//MS, YS/RS and MS/RS

**Table S6** - KEGG pathway term distribution of expressed genes in YS/MS, YS/RS and MS/RS

**Table S7** - Heatmap results for softening genes.

**Table S8** - Heatmap results for signal transduction.

**Table S9** - Genes and metabolising enzymes involved in starch and sucrose metabolism pathway.

