## Supplemental Table S1 for "Transcriptome analysis during fruit developmental stages in durian (*Durio zibethinus* Murr.) var. D24"

Table S1- Differentially up-regulated expressed genes between young stage and mature stage of durian fruit pulp. We used FDR: <0.05, Log 2 fold change >1.5 and <-1.5*.*

| **Gene Symbol** | **Gene Name** | **Log2 fold change** | **FDR p-value correction** |
| --- | --- | --- | --- |
| LOC111285147 | membrane-anchored ubiquitin-fold protein 3-like | 13.3352889 | 0.04012651 |
| LOC111318243 | cytochrome b5-like | 12.5541173 | 0.02234778 |
| LOC111304600 | pectinesterase 2-like | 11.22367883 | 8.13E-26 |
| LOC111293884 | bZIP transcription factor TGA10-like, transcript | 11.20479892 | 3.42E-25 |
| LOC111298007 | probable serine/threonine-protein kinase BSK3 | 11.1190365 | 0.00487093 |
| LOC111278395 | endoglucanase-like | 10.872028 | 0.02831135 |
| LOC111277078 | uncharacterized LOC111277078 | 10.8154348 | 0.00101408 |
| LOC111274386 | ABC transporter C family member 13 | 10.7658514 | 0.01995145 |
| LOC111282227 | uncharacterized LOC111282227 | 10.6329582 | 0.00557413 |
| LOC111286089 | ABC transporter G family member 14-like | 10.5499989 | 0.00515839 |
| LOC111309500 | protein ROOT HAIR DEFECTIVE 3 homolog 2-like | 10.5240047 | 0.00034707 |
| LOC111278272 | uncharacterized LOC111278272 | 10.4641515 | 0.02272886 |
| LOC111315188 | allene oxide synthase 1, chloroplastic-like | 10.44423 | 0.000689 |
| MSTRG.31886 | Unknown sequences | 10.34963 | 0.005651 |
| LOC111306587 | LEAF RUST 10 DISEASE-RESISTANCE LOCUS RECEPTOR-LIKE PROTEIN KINASE-like 2.1 | 10.25225 | 0.011144 |
| LOC111299220 | uncharacterized LOC111299220 | 10.16312 | 1.40E-05 |
| LOC111287425 | methionine gamma-lyase-like | 10.14481 | 0.015062 |
| LOC111289620 | probable methyltransferase PMT21 | 10.1332 | 1.40E-05 |
| LOC111289620 | probable methyltransferase PMT21 | 10.07247152 | 2.00E-129 |
| LOC111283052 | ABSCISIC ACID-INSENSITIVE 5-like protein 7 | 10.06527 | 0.000829 |
| LOC111318371 | probable serine/threonine-protein kinase At1g01540 | 10.01212 | 0.000363 |
| LOC111282902 | uncharacterized LOC111282902 | 10.00101943 | 4.75E-15 |
| LOC111317091 | kinesin-like protein KIN-4A | 9.991628 | 0.001951 |
| LOC111301728 | calmodulin-binding protein 60 B-like | 9.971501 | 3.25E-05 |
| LOC111315188 | allene oxide synthase 1, chloroplastic-like | 9.941797555 | 2.40E-34 |
| LOC111301330 | uncharacterized LOC111301330 | 9.890983 | 0.009864 |
| MSTRG.16320 | Unknown sequences | 9.884381 | 0.017358 |
| LOC111287086 | probable disease resistance protein At1g58602 | 9.863055 | 0.001735 |
| MSTRG.32120 | Unknown sequences | 9.860302 | 0.001304 |
| MSTRG.11262 | Unknown sequences | 9.834361 | 0.008905 |
| LOC111291064 | meiotic recombination protein SPO11-2 | 9.774328 | 0.006075 |
| LOC111287487 | glucan endo-1,3-beta-glucosidase 11-like | 9.762352 | 0.010156 |
| LOC111297770 | DNA-directed RNA polymerase I subunit 1-like | 9.740118 | 0.001338 |
| LOC111302253 | uncharacterized membrane protein YuiD-like | 9.695723 | 0.001014 |
| LOC111306077 | RNA exonuclease 4 | 9.694658 | 0.003033 |
| LOC111299220 | uncharacterized LOC111299220 | 9.688091281 | 5.40E-32 |
| LOC111303519 | SHUGOSHIN 2-like | 9.674148 | 0.013611 |
| LOC111311329 | uncharacterized LOC111311329 | 9.664423 | 0.000407 |
| LOC111301728 | calmodulin-binding protein 60 B-like | 9.643236632 | 5.21E-41 |
| LOC111282399 | ELMO domain-containing protein C-like | 9.641626 | 0.01912 |
| LOC111316978 | ABC transporter G family member 11-like, | 9.632069514 | 1.60E-36 |
| LOC111305461 | serine/threonine-protein kinase OXI1-like | 9.61594 | 0.001961 |
| MSTRG.35109 | Unknown sequences | 9.602434 | 0.000342 |
| LOC111296692 | nudix hydrolase 26, chloroplastic | 9.565827 | 0.000763 |
| LOC111303365 | N-alpha-acetyltransferase 50-like | 9.544586 | 0.009848 |
| LOC111303521 | vacuolar protein sorting-associated protein 24 homolog 1 | 9.527588 | 0.001597 |
| LOC111283052 | ABSCISIC ACID-INSENSITIVE 5-like protein 7 | 9.52463651 | 1.04E-29 |
| MSTRG.20171 | Unknown sequences | 9.522947 | 6.50E-05 |
| LOC111318157 | uncharacterized LOC111318157 | 9.490975 | 0.000683 |
| MSTRG.2542 | Unknown sequences | 9.48254 | 0.000651 |
| LOC111287285 | histidine protein methyltransferase 1 homolog | 9.482239 | 0.024682 |
| LOC111295573 | caffeic acid 3-O-methyltransferase-like | 9.471809985 | 1.82E-11 |
| LOC111304634 | 4-hydroxy-tetrahydrodipicolinate synthase, chloroplastic | 9.448434 | 0.001676 |
| LOC111312385 | uncharacterized LOC111312385 | 9.435338 | 0.017413 |
| MSTRG.11205 | Unknown sequences | 9.413073 | 0.000812 |
| MSTRG.21045 | Unknown sequences | 9.373216 | 0.004234 |
| LOC111287086 | probable disease resistance protein At1g58602 | 9.369810051 | 1.14E-30 |
| LOC111299987 | pentatricopeptide repeat-containing protein At5g39350-like | 9.314051 | 0.002189 |
| LOC111289785 | probable NADH dehydrogenase [ubiquinone] 1 alpha subcomplex subunit 5, mitochondrial | 9.203199 | 0.037451 |
| TRNAT-UGU | Transfer RNA | 9.19047 | 0.003094 |
| LOC111295509 | probable pyridoxal 5'-phosphate synthase subunit PDX1 | 9.17956 | 0.011744 |
| MSTRG.32120 | Unknown sequences | 9.130971593 | 8.32E-21 |
| LOC111304630 | LOB domain-containing protein 18, (Lateral Organ Boundaries) | 9.1229 | 0.000312596 |
| LOC111305461 | serine/threonine-protein kinase OXI1-like | 9.118252994 | 9.81E-28 |
| LOC111304630 | LOB domain-containing protein 18 | 9.103783345 | 1.35E-12 |
| LOC111274414 | post-GPI attachment to proteins factor 3-like | 9.09872768 | 1.36E-34 |
| LOC111298007 | probable serine/threonine-protein kinase BSK3 | 9.03232922 | 4.49E-13 |
| LOC111286965 | uncharacterized LOC111286965 | 8.980394396 | 1.46E-09 |
| LOC111303521 | vacuolar protein sorting-associated protein 24 | 8.98022 | 5.24E-24 |
| LOC111318243 | cytochrome b5-like, transcript variant X1 | 8.977934 | 4.00E-10 |
| LOC111277078 | uncharacterized LOC111277078 | 8.837425 | 4.70E-13 |
| MSTRG.31886 | Unknown sequences | 8.833591 | 1.00E-12 |
| LOC111312807 | tubulin beta-6 chain-like | 8.794125 | 5.65E-29 |
| LOC111275409 | un-characterized LOC111275409 | 8.793297 | 2.21E-18 |
| LOC111283878 | uncharacterized LOC111283878 | 8.788679046 | 2.63E-42 |
| LOC111285475 | 50S ribosomal protein L9, chloroplastic-like, | 8.775158 | 5.65E-61 |
| LOC111283037 | homeobox-leucine zipper protein HOX3-like | 8.74162798 | 1.09E-08 |
| LOC111315258 | 4-coumarate--CoA ligase-like 5 | 8.649321 | 1.08E-28 |
| LOC111309500 | Protein ROOT HAIR DEFECTIVE 3 homolog 2-like | 8.642184 | 4.11E-13 |
| MSTRG.14226 | Unknown sequences | 8.590877 | 1.87E-63 |
| LOC111282227 | uncharacterized LOC111282227 | 8.577669 | 1.07E-11 |
| LOC111286089 | ABC transporter G family member 14-like | 8.500941 | 1.35E-11 |
| LOC111294556 | plant cysteine oxidase 2-like, transcript | 8.498471955 | 1.67E-07 |
| LOC111311552 | polygalacturonase inhibitor-like | 8.43397 | 0.000312596 |
| MSTRG.21045 | Unknown sequences | 8.342354 | 2.22E-13 |
| MSTRG.30203 | Unknown sequences | 8.338256 | 2.21E-29 |
| LOC111298473 | transcription activator GLK1-like | 8.3172436 | 5.17E-20 |
| LOC111289307 | probable serine/threonine-protein kinase PBL15 | 8.2843 | 1.01E-14 |
| LOC111295036 | intron-binding protein aquarius | 8.26521 | 2.62E-21 |
| MSTRG.4155 | Unknown sequences | 8.236132 | 1.24E-39 |
| LOC111274744 | uncharacterized LOC111274744 | 8.227847 | 1.38E-67 |
| LOC111317678 | protein SIEVE ELEMENT OCCLUSION B-like | 8.220546 | 1.61E-14 |
| LOC111318371 | probable serine/threonine-protein kinase At1g01540 | 8.207395 | 6.69E-12 |
| LOC111293182 | ATP-dependent RNA helicase DEAH11, chloroplastic-like | 8.196636 | 4.10E-23 |
| XLOC_006973 | novel transcript | 8.17641 | 0.000312596 |
| LOC111313563 | uncharacterized LOC111313563 | 8.14606986 | 4.12E-07 |
| LOC111301565 | beta-(1,2)-xylosyltransferase-like | 8.144911 | 2.40E-42 |
| MSTRG.19557 | Unknown sequences | 8.13941 | 7.51E-24 |
| LOC111306587 | LEAF RUST 10 DISEASE-RESISTANCE LOCUS RECEPTOR-LIKE PROTEIN KINASE-like 2.1 | 8.129383 | 4.56E-09 |
| MSTRG.31589 | Unknown sequences | 8.076908 | 8.52E-19 |
| LOC111317091 | kinesin-like protein KIN-4A | 8.05726 | 1.50E-10 |
| LOC111287425 | methionine gamma-lyase-like | 8.048428 | 1.02E-09 |
| LOC111292651 | mavicyanin-like | 8.036744907 | 7.37E-46 |
| LOC111303063 | acyl-CoA-binding protein-like | 8.035915981 | 2.99E-26 |
| LOC111311552 | polygalacturonase inhibitor-like | 8.023169229 | 9.88E-18 |
| LOC111295509 | probable pyridoxal 5'-phosphate synthase subunit PDX1 | 7.986618 | 3.10E-11 |
| LOC111281937 | aminotransferase ALD1, chloroplastic-like | 7.960043 | 9.00E-21 |
| LOC111304216 | early nodulin-93-like | 7.944972454 | 7.22E-09 |
| MSTRG.7750 | Unknown sequences | 7.94274 | 2.91E-36 |
| LOC111276436 | ribonuclease 3-like protein 2 | 7.915384 | 2.42E-34 |
| LOC111291994 | histone-lysine N-methyltransferase setd3-like | 7.912995 | 4.70E-11 |
| LOC111311329 | uncharacterized LOC111311329 | 7.905622 | 4.00E-11 |
| LOC111313836 | K(+) efflux antiporter 2, chloroplastic-like | 7.885134 | 9.14E-22 |
| MSTRG.22577 | Unknown sequences | 7.872309 | 1.85E-29 |
| MSTRG.35109 | Unknown sequences | 7.87218 | 3.05E-11 |
| LOC111304140 | hepatocyte growth factor-regulated tyrosine kinase substrate-like | 7.86719 | 4.89E-14 |
| LOC111297770 | DNA-directed RNA polymerase I subunit 1-like | 7.854016 | 4.15E-10 |
| LOC111301330 | uncharacterized LOC111301330 | 7.848045 | 2.21E-09 |
| LOC111302253 | uncharacterized membrane protein YuiD-like | 7.840544 | 2.35E-10 |
| LOC111294297 | uncharacterized LOC111294297, transcript variant X3 | 7.827264502 | 1.89E-27 |
| LOC111298508 | transcription repressor OFP12-like | 7.79210963 | 4.36E-05 |
| LOC111304275 | rust resistance kinase Lr10-like | 7.786161735 | 7.72E-06 |
| LOC111307587 | uncharacterized LOC111307587 | 7.677309697 | 2.63E-32 |
| LOC111292650 | triacylglycerol lipase 2-like | 7.55833 | 0.000312596 |
| LOC111276805 | uncharacterized LOC111276805 | 7.548785254 | 9.00E-17 |
| LOC111309908 | cytochrome P450 77A3-like | 7.19743169 | 0.000195223 |
| LOC111293555 | benzyl alcohol O-benzoyltransferase-like | 7.129697458 | 5.38E-06 |
| LOC111293555 | benzyl alcohol O-benzoyltransferase-like, transcript variant X1 | 7.10972 | 0.000312596 |
| LOC111313396 | RING-H2 finger protein ATL5-like | 6.977802898 | 5.79E-21 |
| LOC111288096 | uncharacterized LOC111288096 | 6.91513012 | 0.000315393 |
| LOC111306284 | serine/threonine-protein kinase rio2-like | 6.897779308 | 0.000624792 |
| LOC111274286 | probable polygalacturonase At3g15720 | 6.888506571 | 0.000902289 |
| LOC111276942 | non-specific phospholipase C4-like | 6.88614166 | 4.83E-23 |
| LOC111305909 | LOB domain-containing protein 19 | 6.78294307 | 3.65E-14 |
| LOC111300249 | uncharacterized LOC111300249 | 6.774395536 | 0.000970999 |
| LOC111281105 | protein DETOXIFICATION 40-like | 6.68363772 | 4.17E-38 |
| LOC111306285 | protein GLUTAMINE DUMPER 2-like | 6.668945983 | 2.33E-11 |
| LOC111280925 | uncharacterized LOC111280925 | 6.659256641 | 0.001160978 |
| LOC111290435 | LOB domain-containing protein 40-like | 6.616392391 | 2.58E-12 |
| LOC111298611 | uncharacterized LOC111298611 | 6.608186011 | 0.002060717 |
| LOC111307587 | uncharacterized LOC111307587 | 6.58076 | 0.00129227 |
| LOC111293921 | calmodulin-like | 6.571492192 | 0.000701268 |
| LOC111282509 | uncharacterized LOC111282509 | 6.55632 | 0.00343537 |
| LOC111281105 | protein DETOXIFICATION 40-like | 6.45873 | 0.000312596 |
| LOC111312958 | uncharacterized LOC111312958 | 6.438727994 | 7.11E-10 |
| LOC111315580 | synaptonemal complex protein 1-like | 6.422667564 | 1.29E-23 |
| LOC111286976 | nuclear transcription factor Y subunit C-2-like | 6.359892775 | 0.011187936 |
| LOC111301968 | uncharacterized LOC111301968 | 6.332188541 | 3.47E-13 |
| LOC111318443 | bidirectional sugar transporter N3 | 6.3118 | 0.000312596 |
| LOC111280213 | cellulose synthase-like protein G2 | 6.278284098 | 1.79E-18 |
| LOC111300836 | nuclear pore complex protein NUP205-like | 6.274587802 | 0.005580112 |
| LOC111282509 | uncharacterized LOC111282509 | 6.266850539 | 1.47E-07 |
| LOC111315580 | synaptonemal complex protein 1-like | 6.25794 | 0.000312596 |
| LOC111305909 | LOB domain-containing protein 19 | 6.20871 | 0.00598521 |
| LOC111283141 | polygalacturonase QRT3-like | 6.200765753 | 0.002915672 |
| LOC111301538 | probable nucleoredoxin 2 | 6.186988971 | 1.73E-08 |
| LOC111318443 | bidirectional sugar transporter N3 | 6.165414964 | 5.10E-06 |
| LOC111301322 | ABC transporter G family member 31 | 6.162000356 | 2.18E-09 |
| LOC111283759 | transcription factor MYB108-like | 6.160669576 | 1.56E-07 |
| LOC111317870 | proline-rich receptor-like protein kinase PERK12 | 6.107352315 | 2.22E-17 |
| LOC111286614 | putative receptor-like protein kinase At3g47110 | 6.092903782 | 0.006511795 |
| LOC111276942 | non-specific phospholipase C4-like | 6.08761 | 0.00308959 |
| LOC111285498 | transmembrane protein 45A-like | 6.06232 | 0.000312596 |
| LOC111290435 | LOB domain-containing protein 40-like | 6.03502 | 0.000312596 |
| LOC111285478 | protein NRT1/ PTR FAMILY 7.1-like, transcript | 6.01832 | 0.000582785 |
| LOC111282412 | LOB domain-containing protein 42-like | 6.0083 | 0.000312596 |
| LOC111301322 | ABC transporter G family member 31, transcript | 5.95744 | 0.000312596 |
| LOC111297059 | LOB domain-containing protein 1-like | 5.65526 | 0.000312596 |
| LOC111295041 | glucan endo-1,3-beta-glucosidase, basic isoform-like | 5.60328 | 0.00614272 |
| LOC111295977 | rop guanine nucleotide exchange factor 7-like | 5.56923 | 0.000312596 |
| LOC111316379 | aspartic proteinase Asp1-like | 5.46952 | 0.000312596 |
| LOC111296764 | probable purple acid phosphatase 20, transcript | 5.42737 | 0.000312596 |
| LOC111301815 | transmembrane protein 45A-like | 5.41779 | 0.000312596 |
| LOC111312665 | protein DETOXIFICATION 33, transcript variant | 5.37174 | 0.000312596 |
| LOC111312381 | uncharacterized methyltransferase At1g78140, | 5.3694 | 0.0214146 |
| LOC111317748 | putative SNAP25 homologous protein SNAP30 | 5.33821 | 0.000312596 |
| LOC111310483 | uncharacterized LOC111310483, transcript variant | 5.32815 | 0.000312596 |
| LOC111311747 | ABC transporter G family member 39-like | 5.29601 | 0.000312596 |
| LOC111317870 | proline-rich receptor-like protein kinase | 5.29238 | 0.000833693 |
| LOC111304064 | probable receptor-like protein kinase At1g80640, | 5.28034 | 0.000312596 |
| LOC111280882 | probable 9-cis-epoxycarotenoid dioxygenase | 5.21689 | 0.000312596 |
| LOC111298913 | uncharacterized LOC111298913 | 5.18175 | 0.000312596 |
| LOC111313976 | uncharacterized LOC111313976 | 5.17833 | 0.000312596 |
| LOC111296606 | zinc finger protein ZAT12-like | 5.17716 | 0.000312596 |
| LOC111274350 | shikimate O-hydroxycinnamoyltransferase-like | 5.03821 | 0.000312596 |
| LOC111290481 | hypersensitive-induced reaction 1 protein-like, | 5.03038 | 0.000312596 |
| LOC111287927 | cysteine synthase 2-like | 5.02663 | 0.000312596 |
| LOC111304767 | uncharacterized LOC111304767 | 4.94095 | 0.000312596 |
| LOC111283735 | NAC domain-containing protein 100-like | 4.86757 | 0.000312596 |
| LOC111299261 | heat stress transcription factor A-2-like, | 4.86241 | 0.000582785 |
| LOC111312254 | probable receptor-like serine/threonine-protein | 4.85289 | 0.000312596 |
| LOC111280497 | probable potassium transporter 13 | 4.84641 | 0.00106839 |
| LOC111309044 | UPF0496 protein At4g34320-like, transcript | 4.82114 | 0.000312596 |
| LOC111316857 | uncharacterized LOC111316857 | 4.81577 | 0.00232875 |
| LOC111312852 | tetracycline resistance protein, class H-like, | 4.80275 | 0.000312596 |
| LOC111283869 | uncharacterized LOC111283869 | 4.79708 | 0.000312596 |
| LOC111276805 | uncharacterized LOC111276805, transcript variant | 4.79352 | 0.000312596 |
| LOC111307626 | probable serine/threonine-protein kinase WNK5 | 4.77008 | 0.000312596 |
| LOC111294332 | feruloyl CoA ortho-hydroxylase 1-like | 4.75322 | 0.0338929 |
| XLOC_024860 | Novel gene | 4.71534 | 0.00129227 |
