## Supplemental Table S2 for "Transcriptome analysis during fruit developmental stages in durian (*Durio zibethinus* Murr.) var. D24"

Table S2 - Differentially down-regulated expressed genes between young stage and mature stage of durian fruit pulp. We used FDR: <0.05, Log2 fold change >1.5 and <-1.5.

| **Gene Symbol** | **Gene Name** | **Log2 fold change** | **FDR p-value correction** |
| --- | --- | --- | --- |
| MSTRG.34481 | Unknown sequences | -5.71934 | 1.26E-06 |
| LOC111301324 | iridoid synthase-like | -5.75332 | 1.08E-12 |
| LOC111286552 | protein PHOTOSYSTEM I ASSEMBLY 2, chloroplastic | -5.75397 | 1.29E-14 |
| LOC111283752 | tubby-like F-box protein 3 | -5.77035 | 6.83E-09 |
| LOC111289347 | synaptotagmin-5-like | -5.78155 | 7.58E-12 |
| MSTRG.17494 | Unknown sequences | -5.81282 | 1.68E-05 |
| MSTRG.9748 | Unknown sequences | -5.84638 | 2.47E-13 |
| LOC111276937 | probable cellulose synthase A catalytic subunit 6 [UDP-forming] | -5.85728 | 3.30E-12 |
| MSTRG.29429 | Unknown sequences | -5.89153 | 9.78E-06 |
| LOC111294072 | ubiquitin carboxyl-terminal hydrolase 2-like | -5.89682 | 1.01E-05 |
| LOC111311962 | BTB/POZ and MATH domain-containing protein 3-like | -5.90035 | 1.11E-05 |
| LOC111314105 | proline-rich protein 3-like | -5.94419 | 9.99E-19 |
| LOC111287804 | receptor-like protein 12 | -5.97622 | 3.13E-07 |
| MSTRG.17496 | Unknown sequences | -6.01054 | 5.48E-06 |
| LOC111293582 | uncharacterized LOC111293582 | -6.0596 | 4.56E-06 |
| LOC111274855 | katanin p60 ATPase-containing subunit A-like 2 | -6.08458 | 1.27E-12 |
| LOC111305908 | AT-hook motif nuclear-localized protein 1-like | -6.15924 | 3.08E-25 |
| LOC111315575 | protein ROOT PRIMORDIUM DEFECTIVE 1 | -6.21274 | 1.01E-05 |
| LOC111282083 | uncharacterized LOC111282083 | -6.2129 | 2.24E-06 |
| LOC111289814 | probable methyltransferase PMT18 | -6.25643 | 1.40E-13 |
| LOC111280314 | calcium-dependent protein kinase 21-like | -6.26744 | 2.29E-78 |
| MSTRG.4787 | Unknown sequences | -6.34128 | 4.11E-14 |
| LOC111292232 | pentatricopeptide repeat-containing protein ELI1, chloroplastic | -6.36727 | 6.84E-07 |
| LOC111292756 | UPF0548 protein At2g17695 | -6.49622 | 0.011606 |
| MSTRG.9760 | Unknown sequences | -6.53345 | 0.010832 |
| LOC111304213 | protein kinase PINOID 2-like | -6.5739 | 0.041848 |
| MSTRG.11548 | Unknown sequences | -6.60747 | 0.010324 |
| LOC111289814 | probable methyltransferase PMT18 | -6.62664 | 0.00022 |
| LOC111307584 | probable serine/threonine-protein kinase PBL8 | -6.62792 | 6.60E-31 |
| MSTRG.5729 | Unknown sequences | -6.62804 | 7.25E-07 |
| LOC111287804 | receptor-like protein 12 | -6.63194 | 0.001394 |
| LOC111291144 | protein C2-DOMAIN ABA-RELATED 4-like | -6.64067 | 0.012722 |
| MSTRG.12961 | Unknown sequences | -6.65997 | 8.47E-13 |
| MSTRG.36427 | Unknown sequences | -6.66604 | 0.010238 |
| LOC111302841 | histidine kinase 5-like | -6.68511 | 1.50E-18 |
| MSTRG.4787 | Unknown sequences | -6.69632 | 0.00031 |
| LOC111283068 | phytosulfokine receptor 1-like | -6.74285 | 2.44E-27 |
| LOC111318002 | bidirectional sugar transporter SWEET10-like | -6.74348 | 0.008228 |
| MSTRG.25731 | Unknown sequences | -6.75191 | 7.58E-10 |
| MSTRG.17492 | Unknown sequences | -6.75428 | 2.37E-13 |
| MSTRG.7090 | Unknown sequences | -6.76053 | 6.66E-08 |
| MSTRG.32710 | Unknown sequences | -6.76884 | 3.72E-27 |
| LOC111304629 | uncharacterized LOC111304629 | -6.81839 | 6.04E-08 |
| LOC111307584 | probable serine/threonine-protein kinase PBL8 | -6.86817 | 0.000252 |
| MSTRG.2851 | Unknown sequences | -6.91285 | 0.007325 |
| MSTRG.33993 | Unknown sequences | -6.95097 | 4.24E-29 |
| MSTRG.32710 | Unknown sequences | -7.01124 | 0.000202 |
| LOC111302841 | histidine kinase 5-like | -7.01672 | 0.00019 |
| LOC111283068 | phytosulfokine receptor 1-like | -7.06065 | 0.001304 |
| LOC111295507 | uncharacterized LOC111295507 | -7.07453 | 0.036571 |
| MSTRG.12961 | Unknown sequences | -7.11651 | 0.000552 |
| MSTRG.33993 | Unknown sequences | -7.20243 | 0.000174 |
| MSTRG.17492 | Unknown sequences | -7.20892 | 0.000183 |
| MSTRG.3263 | Unknown sequences | -7.22556 | 8.31E-16 |
| MSTRG.15890 | Unknown sequences | -7.34197 | 0.004234 |
| LOC111311589 | protein trichome birefringence-like | -7.34446 | 0.00901342 |
| LOC111315804 | ethylene-responsive transcription factor | -7.35398 | 0.0314026 |
| LOC111287953 | xyloglucan endotransglucosylase/hydrolase | -7.36103 | 0.000312596 |
| LOC111279922 | calmodulin-like protein 8, transcript variant | -7.3707 | 0.000312596 |
| LOC111290826 | nucleobase-ascorbate transporter 6-like | -7.37673 | 0.000312596 |
| LOC111293453 | 3-dehydrosphinganine reductase TSC10A-like | -7.37723 | 2.57E-72 |
| LOC111318818 | mitogen-activated protein kinase kinase kinase 18-like | -7.37864 | 0.00752884 |
| LOC111278288 | calcium permeable stress-gated cation channel | -7.38262 | 0.000312596 |
| LOC111281901 | transcription factor MYB62-like | -7.38509 | 0.000582785 |
| LOC111292968 | probable xyloglucan endotransglucosylase/hydrolase protein 6 | -7.3922 | 0.0202149 |
| LOC111298258 | uncharacterized LOC111298258 | -7.39728 | 0.006276 |
| MSTRG.25731 | Unknown sequences | -7.41758 | 0.000317 |
| MSTRG.17494 | Unknown sequences | -7.48989 | 0.0037 |
| LOC111275658 | aspartic proteinase-like | -7.49913 | 0.000312596 |
| LOC111274739 | subtilisin-like protease SBT1.5 | -7.53268 | 0.00397856 |
| LOC111293453 | 3-dehydrosphinganine reductase TSC10A-like | -7.54033 | 1.40E-05 |
| MSTRG.29429 | Unknown sequences | -7.54618 | 0.003054 |
| LOC111294072 | ubiquitin carboxyl-terminal hydrolase 2-like | -7.56002 | 0.003163 |
| LOC111311962 | BTB/POZ and MATH domain-containing protein 3-like | -7.57949 | 0.003426 |
| MSTRG.33951 | Unknown sequences | -7.5839 | 2.38E-10 |
| LOC111277773 | dehydration-responsive element-binding protein | -7.65162 | 0.0106799 |
| LOC111283075 | transcription factor bHLH162-like | -7.65741 | 0.0243923 |
| LOC111315575 | protein ROOT PRIMORDIUM DEFECTIVE 1 | -7.66979 | 0.00143 |
| MSTRG.17496 | Unknown sequences | -7.67053 | 0.00278 |
| LOC111318712 | phospholipase A1-IIbeta-like | -7.68886 | 0.00192383 |
| MSTRG.3263 | Unknown sequences | -7.69625 | 0.000272 |
| LOC111293487 | GEM-like protein 5 | -7.71764 | 0.000312596 |
| LOC111316856 | glucose-1-phosphate adenylyltransferase large subunit 1, chloroplastic-like | -7.72275 | 1.39E-10 |
| LOC111293582 | uncharacterized LOC111293582 | -7.72921 | 0.00278 |
| LOC111284733 | probable galactinol--sucrose | -7.73393 | 0.000312596 |
| LOC111314503 | fasciclin-like arabinogalactan protein 2 | -7.75301 | 0.012623 |
| LOC111286965 | uncharacterized LOC111286965 | -7.76867 | 7.27E-11 |
| LOC111289953 | salutaridinol 7-O-acetyltransferase-like | -7.83201 | 0.000312596 |
| LOC111295285 | uncharacterized LOC111295285 | -7.89381 | 1.47E-14 |
| LOC111282083 | uncharacterized LOC111282083 | -7.89568 | 0.002551 |
| LOC111293109 | AAA-ATPase At3g28610-like | -7.90206 | 0.00887402 |
| LOC111309924 | uncharacterized LOC111309924 | -7.916390563 | 1.92E-39 |
| LOC111309093 | protein EXORDIUM-like 2 | -7.94134 | 0.00887402 |
| LOC111292232 | pentatricopeptide repeat-containing protein ELI1, chloroplastic | -8.0167 | 0.001792 |
| LOC111303423 | probable xyloglucan endotransglucosylase/hydrolase protein 23 | -8.02948 | 0.000312596 |
| LOC111316185 | LOB domain-containing protein 38-like | -8.06646 | 0.00929446 |
| LOC111305904 | axial regulator YABBY 1-like | -8.071802227 | 2.28E-15 |
| LOC111275659 | mitogen-activated protein kinase kinase kinase | -8.08745 | 0.0106799 |
| LOC111303287 | galactosyltransferase 6 | -8.11331 | 0.000312596 |
| LOC111311549 | ras-related protein RABA4d-like | -8.114019028 | 8.45E-37 |
| MSTRG.5729 | Unknown sequences | -8.13011 | 0.014178 |
| LOC111314766 | phosphoprotein ECPP44-like | -8.13335 | 0.000312596 |
| LOC111278190 | stem-specific protein TSJT1-like | -8.16401 | 0.000312596 |
| LOC111309924 | uncharacterized LOC111309924 | -8.18371 | 0.000157 |
| LOC111298174 | uncharacterized LOC111298174 | -8.29316 | 0.000312596 |
| LOC111274745 | pectinesterase 3 | -8.29518 | 0.000312596 |
| LOC111301563,LOC111301566 | glycerophosphodiester phosphodiesterase | -8.3828 | 0.000833693 |
| MSTRG.7377 | Unknown sequences | -8.39974446 | 8.46E-23 |
| LOC111287471 | F-box/kelch-repeat protein At1g67480-like, | -8.40684 | 0.000312596 |
| MSTRG.7090 | Unknown sequences | -8.42358 | 0.001169 |
| LOC111292380 | mitochondrial pyruvate carrier 1-like, | -8.47976 | 0.000312596 |
| LOC111311549 | ras-related protein RABA4d-like | -8.4981 | 0.00118 |
| LOC111304629 | uncharacterized LOC111304629 | -8.507 | 0.001296 |
| LOC111281172 | probable WRKY transcription factor 40 | -8.51332 | 0.000312596 |
| LOC111274880 | LRR receptor-like serine/threonine-protein | -8.58511 | 0.000312596 |
| LOC111294535 | protein SLE1 | -8.58778 | 0.0361286 |
| LOC111305909 | LOB domain-containing protein 19 | -8.599375192 | 1.08E-31 |
| LOC111283076 | peroxidase N1-like | -8.62056 | 0.000312596 |
| LOC111295285 | uncharacterized LOC111295285 | -8.62972 | 8.25E-05 |
| LOC111312403 | protein EXORDIUM-like | -8.77531 | 0.000312596 |
| LOC111305904 | axial regulator YABBY 1-like | -8.82141 | 7.15E-05 |
| LOC111290894 | abscisic acid 8'-hydroxylase 1-like | -8.83601 | 0.000312596 |
| MSTRG.7377 | Unknown sequences | -8.9350345 | 3.14E-05 |
| LOC111283603 | protein RALF-like 27 | -8.992785464 | 5.35E-27 |
| LOC111316977 | uncharacterized LOC111316977 | -9.005287613 | 2.30E-20 |
| LOC111305909 | LOB domain-containing protein 19 | -9.1192627 | 0.00206985 |
| LOC111293917 | omega-3 fatty acid desaturase, | -9.13208 | 0.00212999 |
| LOC111315262 | L-ascorbate oxidase homolog | -9.13668 | 0.0130188 |
| LOC111316936 | cytochrome P450 CYP736A12-like | -9.21472 | 0.0193685 |
| LOC111284909 | asparagine synthetase [glutamine-hydrolyzing] 1 | -9.21616 | 0.00343537 |
| LOC111312815 | chaperone protein dnaJ 11, chloroplastic-like | -9.25731 | 0.038914 |
| MSTRG.33951 | Unknown sequences | -9.2749643 | 0.00041031 |
| LOC111303526 | heavy metal-associated | -9.28425 | 0.000312596 |
| LOC111281364 | dehydration-responsive element-binding protein 1A-like | -9.37327 | 0.000582785 |
| LOC111316856 | glucose-1-phosphate adenylyltransferase large subunit 1, chloroplastic-like | -9.4525362 | 0.0004438 |
| LOC111286965 | uncharacterized LOC111286965 | -9.4810942 | 0.00035447 |
| LOC111283603 | protein RALF-like 27 | -9.5559029 | 2.18E-05 |
| LOC111288324 | UDP-glucuronate 4-epimerase 6-like | -9.65161 | 0.00451342 |
| LOC111295419 | protein CDI-like | -9.7041 | 0.00999511 |
| MSTRG.31294 | Unknown sequences | -9.711706002 | 1.28E-17 |
| LOC111305464 | auxin-induced in root cultures protein 12-like | -9.74314 | 0.0108235 |
| LOC111309042 | transcription termination factor MTEF1, chloroplastic | -9.800410409 | 5.82E-25 |
| LOC111316977 | uncharacterized LOC111316977 | -9.8066878 | 3.34E-05 |
| LOC111304131 | uncharacterized LOC111304131 | -10.19820124 | 1.86E-07 |
| LOC111294115 | protein NUCLEAR FUSION DEFECTIVE 4-like | -10.21194632 | 0.000279608 |
| LOC111308224 | actin-depolymerizing factor 5 | -10.24748068 | 2.13E-10 |
| LOC111299223 | E3 ubiquitin-protein ligase ATL4-like | -10.28077601 | 1.03E-47 |
| LOC111291309 | abscisate beta-glucosyltransferase-like | -10.2951 | 0.000312596 |
| LOC111318679 | GDSL esterase/lipase At1g29670-like | -10.29761004 | 1.93E-07 |
| LOC111284187 | ACT domain-containing protein ACR2 | -10.29959505 | 8.68E-08 |
| LOC111310206 | triosephosphate isomerase, cytosolic | -10.3003751 | 1.81E-10 |
| LOC111308606 | cytochrome P450 CYP82D47-like | -10.30891723 | 1.89E-28 |
| LOC111308711 | expansin-like A2 | -10.31275919 | 7.64E-16 |
| LOC111292555 | probable galacturonosyltransferase 15 | -10.3326013 | 2.62E-16 |
| LOC111287561 | hydroquinone glucosyltransferase-like | -10.33795766 | 3.53E-09 |
| LOC111289734 | protein STICHEL-like 2 | -10.34842623 | 0.000478496 |
| LOC111305464 | auxin-induced in root cultures protein 12-like | -10.3510225 | 1.11E-26 |
| LOC111303366 | probable boron transporter 6 | -10.38437493 | 1.35E-10 |
| LOC111296264 | galactinol synthase 2-like | -10.47539103 | 2.11E-18 |
| LOC111280367 | sugar carrier protein C-like | -10.4941838 | 6.74E-05 |
| LOC111301331 | uncharacterized LOC111301331 | -10.49676028 | 1.34E-11 |
| LOC111295419 | protein CDI-like | -10.5042274 | 6.30E-60 |
| LOC111318057 | MLP-like protein 43 | -10.50850491 | 4.08E-12 |
| LOC111291687 | 3-ketoacyl-CoA synthase 11-like | -10.56048853 | 7.26E-09 |
| LOC111274387 | probable WRKY transcription factor 40 | -10.60567034 | 1.59E-11 |
| LOC111318371 | probable serine/threonine-protein kinase At1g01540 | -10.63722478 | 6.63E-17 |
| LOC111306117 | protein DETOXIFICATION 49-like | -10.65290621 | 8.57E-07 |
| LOC111285821 | 12-oxophytodienoate reductase 3-like | -10.65437297 | 4.17E-08 |
| LOC111315188 | allene oxide synthase 1, chloroplastic-like | -10.662 | 0.0496018 |
| LOC111285984 | glucomannan 4-beta-mannosyltransferase 2-like | -10.66770154 | 4.73E-07 |
| LOC111309042 | transcription termination factor MTEF1, chloroplastic | -10.673562 | 1.69E-05 |
| LOC111317093 | ethylene-responsive transcription factor RAP2-10-like | -10.69811349 | 1.43E-16 |
| LOC111311239 | WAT1-related protein At1g21890-like | -10.70063849 | 1.42E-07 |
| LOC111274892 | probable WRKY transcription factor 28 | -10.7513683 | 6.14E-05 |
| LOC111278395 | endoglucanase-like | -10.80853179 | 1.31E-10 |
| LOC111316161 | transcription factor bHLH18-like | -10.84843791 | 9.30E-08 |
| LOC111301786 | beta-amyrin 28-oxidase-like | -10.88131029 | 2.40E-07 |
| LOC111290490 | ethylene-responsive transcription factor | -10.93766998 | 2.98E-19 |
| LOC111290758 | bidirectional sugar transporter SWEET7-like | -10.995443 | 5.40E-10 |
| LOC111295502 | GATA transcription factor 8-like, transcript | -11.0041165 | 5.26E-09 |
| LOC111285778 | LOB domain-containing protein 12 | -11.01957649 | 1.50E-05 |
| LOC111278395 | endoglucanase-like | -11.0334 | 0.000833693 |
| LOC111289922 | transcription factor bHLH162-like | -11.08706906 | 1.13E-08 |
| LOC111290676 | monocopper oxidase-like protein SKU5 | -11.14908047 | 5.63E-08 |
| LOC111279835 | inorganic phosphate transporter 1-4-like | -11.16340772 | 1.34E-05 |
| LOC111314108 | probable pectinesterase/pectinesterase inhibitor | -11.1741725 | 7.75E-08 |
| LOC111314312 | inorganic phosphate transporter 1-4-like | -11.19136459 | 1.08E-05 |
| LOC111308606 | cytochrome P450 CYP82D47-like | -11.214392 | 1.69E-05 |
| LOC111309796 | CASP-like protein 4D1 | -11.24042127 | 3.13E-23 |
| LOC111315188 | allene oxide synthase 1, chloroplastic-like | -11.32095844 | 3.09E-32 |
| LOC111275649 | glucan endo-1,3-beta-glucosidase 12-like | -11.43236842 | 4.59E-10 |
| MSTRG.31294 | Unknown sequences | -11.594121 | 6.05E-05 |
| LOC111284125 | protein P21-like | -11.67915157 | 5.99E-07 |
| LOC111296083 | uncharacterized protein At1g04910-like, | -11.78699476 | 1.34E-21 |
| LOC111285489 | ethylene-responsive transcription factor TINY-like | -11.83986588 | 5.35E-09 |
| LOC111291469 | protein NRT1/ PTR FAMILY 2.11-like | -12.05371469 | 7.75E-08 |
| LOC111281964 | receptor-like protein kinase HAIKU2 | -12.54736279 | 4.72E-11 |
| LOC111318240 | protein NRT1/ PTR FAMILY 5.5-like | -13.11914601 | 1.44E-21 |
| LOC111289849 | bidirectional sugar transporter SWEET10-like | -13.95054439 | 2.49E-18 |
| LOC111275112 | sugar carrier protein C | -14.35919627 | 5.18E-08 |
