## Supplemental Table S3 for "Transcriptome analysis during fruit developmental stages in durian (*Durio zibethinus* Murr.) var. D24"

Table S3 - Differentially up-regulated expressed genes between mature stage and ripening stage of durian fruit pulp. We used FDR: <0.05, Log2 fold change >1.5 and <-1.5

| **Gene Symbol** | **Gene Name** | **Log2 fold change** | **FDR p-value correction** |
| --- | --- | --- | --- |
| LOC111295531 | glucan endo-1,3-beta-glucosidase, basic | 11.12739807 | 2.40E-13 |
| LOC111304243 | uncharacterized LOC111304243 | 10.82842026 | 3.19E-37 |
| LOC111304213 | protein kinase PINOID 2-like | 10.81391 | 2.20E-09 |
| LOC111316688 | probable leucine-rich repeat receptor-like | 10.73250792 | 3.19E-36 |
| LOC111316289 | probable leucine-rich repeat receptor-like | 10.18733519 | 1.32E-24 |
| LOC111304213 | protein kinase PINOID 2-like | 10.10328474 | 4.43E-91 |
| LOC111296590 | glutathione S-transferase U8-like | 9.819365319 | 2.58E-65 |
| LOC111283910 | uncharacterized protein At1g04910-like, | 9.624435594 | 7.93E-23 |
| LOC111299988 | ethylene-responsive transcription factor | 9.575002879 | 1.25E-20 |
| LOC111305012 | beta-amylase 3, chloroplastic-like | 9.545033365 | 7.88E-24 |
| LOC111274639 | protein indeterminate-domain 7-like, transcript | 9.538087376 | 1.35E-13 |
| LOC111303682 | 17.3 kDa class I heat shock protein-like | 9.526186953 | 4.23E-21 |
| LOC111317014 | probable mannitol dehydrogenase | 9.51845 | 0.0167276 |
| LOC111311521 | polygalacturonase-like | 9.48951 | 0.00020799 |
| MSTRG.23678 | Unknown sequences | 9.468612 | 2.00E-12 |
| LOC111312780 | probable LRR receptor-like serine/threonine-protein kinase At4g36180 | 9.419361796 | 1.65E-11 |
| LOC111318053 | WUSCHEL-related homeobox 6-like | 9.392742016 | 5.40E-20 |
| LOC111277545 | formin-like protein 16 | 9.375942033 | 6.21E-12 |
| LOC111292776 | transcription factor HEC3-like | 9.346581074 | 2.57E-10 |
| LOC111275124 | zinc finger protein CONSTANS-LIKE 4-like, | 9.272614704 | 9.57E-18 |
| LOC111282979 | auxin efflux carrier component 7-like | 9.247544358 | 2.63E-07 |
| LOC111288554 | uncharacterized LOC111288554, transcript variant X1 | 9.221445682 | 2.02E-13 |
| LOC111311996 | laccase-7-like | 9.168182881 | 1.54E-10 |
| LOC111311521 | polygalacturonase-like | 9.166818974 | 1.67E-24 |
| LOC111306867 | putative E3 ubiquitin-protein ligase LIN-1, | 9.119078894 | 1.17E-14 |
| LOC111276419 | fasciclin-like arabinogalactan protein 1 | 9.103103988 | 2.27E-13 |
| LOC111311224 | probable calcium-binding protein CML44 | 9.097227318 | 1.41E-20 |
| LOC111276733 | uncharacterized LOC111276733 | 9.064677 | 9.26E-08 |
| LOC111294224 | hevein-like preproprotein | 9.039798581 | 6.52E-11 |
| LOC111278395 | endoglucanase-like | 9.03817 | 0.00206807 |
| LOC111317014 | probable mannitol dehydrogenase | 9.018448275 | 5.03E-28 |
| LOC111291461 | putative calcium-binding protein CML19 | 9.008240216 | 6.76E-13 |
| LOC111318057 | MLP-like protein 43 | 8.970261229 | 1.38E-09 |
| LOC111300619 | uncharacterized protein LOC111300619 isoform X1 | 8.930274519 | 2.60E-08 |
| LOC111308606 | cytochrome P450 CYP82D47-like | 8.906831 | 4.60E-13 |
| LOC111282766 | transcription factor TCP8-like | 8.884562 | 1.29E-07 |
| LOC111295536 | exocyst complex component EXO70H1-like | 8.82363 | 0.00516878 |
| LOC111317417 | probable xyloglucan endotransglucosylase/hydrolase protein 33 | 8.793108152 | 1.17E-06 |
| LOC111287629 | small heat shock protein, chloroplastic-like | 8.76183 | 0.00727459 |
| LOC111278348 | uncharacterized protein LOC111278348 | 8.725124166 | 1.15E-83 |
| LOC111278395 | endoglucanase-like | 8.694660578 | 4.63E-126 |
| LOC111296235 | abscisic acid 8'-hydroxylase 4-like isoform X1 | 8.664260631 | 3.16E-25 |
| LOC111294546 | hsp70 nucleotide exchange factor fes1-like | 8.65954 | 8.86E-14 |
| LOC111306385 | LOW QUALITY PROTEIN: aluminum-activated malate transporter 10-like | 8.605369446 | 1.53E-21 |
| LOC111311549 | ras-related protein RABA4d-like | 8.605315 | 8.62E-14 |
| LOC111295536 | xocyst complex component EXO70H1-like | 8.539212156 | 8.88E-55 |
| LOC111298613 | equilibrative nucleotide transporter 3-like | 8.494437461 | 3.89E-09 |
| LOC111307281 | crocetin glucosyltransferase, chloroplastic-like | 8.492268702 | 2.16E-09 |
| LOC111283003 | Citrate regulatory gene2 protein-like isoform X1 | 8.425474473 | 5.68E-15 |
| LOC111275649 | glucan endo-1,3-beta-glucosidase 12-like | 8.393470393 | 6.00E-06 |
| LOC111285630 | copper transporter 1-like | 8.38687971 | 1.24E-07 |
| LOC111294910 | arabinogalactan peptide 13-like | 8.380379689 | 2.07E-07 |
| LOC111294546 | hsp70 nucleotide exchange factor fes1-like | 8.368659227 | 4.37E-115 |
| LOC111296858 | glutathione S-transferase U8-like | 8.363805553 | 8.88E-57 |
| LOC111300662 | stress enhanced protein 2, chloroplastic-like | 8.34376 | 0.0435014 |
| LOC111311549 | ras-related protein RABA4d-like | 8.310783961 | 1.74E-113 |
| LOC111285272 | uncharacterized protein LOC111285272 | 8.298631469 | 5.68E-101 |
| LOC111295761 | uncharacterized protein LOC111295761 | 8.276028728 | 5.77E-10 |
| LOC111304696 | cinnamoyl-CoA reductase-like SNL6 | 8.238662814 | 9.14E-08 |
| LOC111305934 | actin-depolymerizing factor 10 | 8.223795205 | 6.37E-14 |
| LOC111308606 | cytochrome P450 CYP82D47-like | 8.203264708 | 4.73E-48 |
| LOC111302013 | BURP domain protein RD22-like | 8.195761277 | 3.18E-36 |
| LOC111299025 | heptahelical transmembrane protein 1-like isoform X1 | 8.16854674 | 4.73E-36 |
| LOC111300662 | stress enhanced protein 2, chloroplastic-like | 8.140500837 | 3.42E-36 |
| LOC111316166 | probable xyloglucan endotransglucosylase/hydrolase protein 6 | 8.13317405 | 1.00E-05 |
| LOC111285251 | protein LURP-one-related 8-like | 8.127684788 | 1.90E-08 |
| LOC111274209 | uncharacterized protein LOC111274209 | 8.11798426 | 1.50E-22 |
| LOC111312913 | probable serine/threonine-protein kinase PIX13 | 8.009359 | 5.15E-13 |
| LOC111291309 | abscisate beta-glucosyltransferase-like | 7.96966 | 0.00020799 |
| LOC111280333 | pseudogene | 7.965274278 | 1.32E-27 |
| LOC111293048 | probable Histone-lysine N-methyltransferase ATXR5 | 7.899614 | 5.53E-06 |
| LOC111278572 | inorganic pyrophosphatase 2-like | 7.82673 | 0.00020799 |
| LOC111311985 | glutathione S-transferase U8-like | 7.71176 | 0.00020799 |
| LOC111285272 | uncharacterized LOC111285272 | 7.67617 | 0.00287618 |
| LOC111293717 | S-adenosylmethionine synthase 3-like | 7.630571 | 5.98E-05 |
| LOC111287154 | sodium/calcium exchanger NCL-like | 7.62475 | 0.0160447 |
| LOC111305387 | sm-like protein LSM5 | 7.466335 | 3.86E-05 |
| LOC111312192 | probable choline kinase 1, transcript variant | 7.45644 | 0.00020799 |
| LOC111312913 | probable serine/threonine-protein kinase PIX13 | 7.448223943 | 2.76E-43 |
| MSTRG.33739 | Unknown sequences | 7.375775 | 6.57E-10 |
| LOC111310012 | putative disease resistance protein At3g14460 | 7.272589 | 7.62E-05 |
| MSTRG.5729 | Unknown sequences | 7.237462 | 1.97E-12 |
| LOC111311763 | pentatricopeptide repeat-containing protein At5g12100, mitochondrial-like | 7.234828 | 6.26E-05 |
| LOC111294072 | ubiquitin carboxyl-terminal hydrolase 2-like | 7.229179 | 7.16E-05 |
| MSTRG.29429 | Unknown sequences | 7.213589 | 6.82E-05 |
| LOC111288064 | protein WVD2-like 4, transcript variant X1 | 7.19024 | 0.00926099 |
| LOC111294224 | hevein-like preproprotein | 7.13727 | 0.016141 |
| MSTRG.5729 | Unknown sequences | 7.068946481 | 3.11E-196 |
| LOC111298258 | uncharacterized LOC111298258 | 7.061663 | 0.000349 |
| LOC111299242 | 17.1 kDa class II heat shock protein-like | 7.03782 | 0.00039263 |
| MSTRG.15890 | Unknown sequences | 7.007487 | 0.000163 |
| MSTRG.36117 | Unknown sequences | 6.985456 | 1.41E-08 |
| LOC111293487 | GEM-like protein 5 | 6.91324 | 0.00020799 |
| MSTRG.18936 | Unknown sequences | 6.905658 | 0.000288 |
| LOC111311546 | OTU domain-containing protein At3g57810-like | 6.87995 | 0.000378 |
| LOC111274057 | NADH-ubiquinone oxidoreductase 20.9 kDa subunit-like | 6.868571 | 0.000225 |
| LOC111282845 | probable nucleoredoxin 1 | 6.86519 | 0.00020799 |
| LOC111303701 | 17.5 kDa class I heat shock protein-like | 6.73551 | 0.00020799 |
| LOC111295774 | zinc finger protein ZAT10-like | 6.71977 | 0.00020799 |
| LOC111287951 | NADH-ubiquinone oxidoreductase 20.9 kDa subunit-like | 6.71362 | 6.67E-07 |
| LOC111318631 | dormancy-associated protein homolog 4-like | 6.70525 | 0.0184336 |
| LOC111303683 | 7.3 kDa class I heat shock protein-like | 6.69473 | 0.028259 |
| LOC111292230 | crocetin glucosyltransferase, | 6.59706 | 0.00670448 |
| LOC111305070 | beta-glucosidase 11-like | 6.58271 | 0.0424588 |
| LOC111296619 | glutathione S-transferase U8-like | 6.53719 | 0.00020799 |
| MSTRG.12961 | Unknown sequences | 6.516409 | 7.07E-09 |
| LOC111276049 | uncharacterized LOC111287951 | 6.511039 | 0.000795 |
| LOC111287951 | uncharacterized LOC111287951 | 6.505450314 | 9.18E-90 |
| LOC111304849 | uncharacterized protein At4g00950-like | 6.488541 | 4.31E-07 |
| LOC111294289 | protein STRICTOSIDINE SYNTHASE-LIKE 10-like | 6.480273 | 0.000765 |
| MSTRG.33739 | Unknown sequences | 6.453038713 | 9.34E-20 |
| LOC111288059 | uncharacterized LOC111288059 | 6.44202 | 0.0178791 |
| LOC111318002 | bidirectional sugar transporter SWEET10-like | 6.411296 | 0.000958 |
| LOC111308755 | protein EARLY-RESPONSIVE TO DEHYDRATION 7, | 6.40309 | 0.00020799 |
| MSTRG.4785 | Unknown sequences | 6.395721 | 1.07E-12 |
| LOC111306845 | uncharacterized LOC111306845 | 6.392457 | 0.001692 |
| LOC111288524 | probable glutathione S-transferase | 6.34518 | 0.00020799 |
| TRNAG-CCC | Transfer RNA | 6.298628 | 6.36E-10 |
| LOC111277539 | protein PHR1-LIKE 1-like | 6.29326 | 3.13E-11 |
| LOC111283603 | protein RALF-like 27 | 6.292572 | 3.74E-12 |
| LOC111274403 | BRCA1-associated protein-like | 6.254206 | 0.001801 |
| LOC111279066 | ankyrin repeat-containing protein | 6.24248 | 0.00104814 |
| LOC111282160 | homeobox-leucine zipper protein HAT5-like | 6.23377 | 5.44E-11 |
| MSTRG.4785 | Unknown sequences | 6.221518293 | 1.79E-107 |
| LOC111303688 | class I heat shock protein-like | 6.20333 | 0.00020799 |
| LOC111287988 | protein SRC2-like | 6.19765 | 0.00020799 |
| LOC111286287 | uncharacterized LOC111286287 | 6.196615 | 0.001955 |
| LOC111276733 | uncharacterized LOC111276733 | 6.189345 | 8.78E-12 |
| LOC111308603 | cytochrome P450 82C4-like | 6.161448 | 0.003852 |
| MSTRG.14611 | Unknown sequences | 6.146772 | 0.002613 |
| LOC111292968 | probable xyloglucan | 6.1364 | 0.0303538 |
| LOC111304865 | pentatricopeptide repeat-containing protein At5g16420, mitochondrial | 6.10152 | 8.79E-08 |
| LOC111283809 | protein VACUOLELESS1-like | 6.101143 | 0.002969 |
| MSTRG.36117 | Unknown sequences | 6.076991 | 3.55E-17 |
| LOC111287953 | xyloglucan endotransglucosylase/hydrolase | 6.05873 | 0.00020799 |
| MSTRG.25731 | Unknown sequences | 6.051906 | 3.17E-11 |
| LOC111296917 | F-box/LRR-repeat protein 3-like | 6.04753 | 0.0069325 |
| LOC111283603 | protein RALF-like 27 | 6.033307 | 1.36E-50 |
| LOC111311009 | UDP-glycosyltransferase 73C6-like | 6.02147 | 0.0179651 |
| LOC111315694 | uncharacterized LOC111315694 | 6.021064 | 0.003436 |
| MSTRG.32862 | Unknown sequences | 6.017646 | 0.003221 |
| LOC111300451 | VQ motif-containing protein 31-like | 6.00263 | 0.0320016 |
| LOC111298889 | probable glutathione S-transferase | 6.00115 | 0.00020799 |
| LOC111300248 | uncharacterized LOC111300248 | 6.000248 | 0.003944 |
| TRNAE-CUC | Transfer RNA | 5.995896 | 1.36E-12 |
| LOC111306167 | uncharacterized LOC111306167 | 5.99026 | 0.00020799 |
| LOC111282160 | homeobox-leucine zipper protein HAT5-like | 5.971007 | 2.87E-48 |
| MSTRG.25133 | Unknown sequences | 5.967606 | 0.003794 |
| LOC111310303 | MLO-like protein 6 | 5.9592 | 0.005599 |
| LOC111285576 | chitotriosidase-1-like | 5.9544 | 0.00553144 |
| LOC111274382 | putative 12-oxophytodienoate reductase 11 | 5.94491 | 0.00860083 |
| LOC111293969 | phosphoenolpyruvate carboxylase kinase 1-like, transcript variant X1 | 5.92943 | 0.00020799 |
| LOC111303611 | zinc finger protein ZAT12-like | 5.92177 | 0.00020799 |
| LOC111277539 | protein PHR1-LIKE 1-like | 5.902129 | 4.76E-30 |
| LOC111284075 | probable disease resistance protein At4g27220 | 5.899992 | 0.004545 |
| MSTRG.22660 | LOC111278267 | 5.896968 | 1.41E-08 |
| LOC111296955 | E3 ubiquitin-protein ligase ATL31-like | 5.89603 | 0.00020799 |
| LOC111277190 | perakine reductase-like | 5.87857 | 0.0013488 |
| LOC111316515 | GEM-like protein 5 | 5.86146 | 0.00020799 |
| MSTRG.15123 | Unknown sequences | 5.852701 | 0.000146 |
| LOC111286661 | uncharacterized LOC111286661 | 5.83691 | 0.018531 |
| TRNAE-CUC | Transfer RNA | 5.830207 | 2.46E-80 |
| LOC111307193 | probable WRKY transcription factor 70 | 5.82317 | 0.00119934 |
| LOC111316628 | probable galactinol--sucrose | 5.8137 | 0.00056455 |
| LOC111313028 | BAG family molecular chaperone regulator 6-like | 5.81305 | 0.0179651 |
| LOC111312788 | uncharacterized LOC111312788 | 5.80826 | 0.00020799 |
| LOC111290564 | vignain-like | 5.7863 | 0.00020799 |
| TRNAG-CCC | Transfer RNA | 5.717676 | 1.76E-19 |
| MSTRG.12961 | Unknown sequences | 5.650746 | 4.26E-14 |
| MSTRG.15123 | Unknown sequences | 5.454912 | 1.08E-28 |
| LOC111278267 | uncharacterized LOC111278267 | 5.345298 | 1.06E-16 |
| MSTRG.34120 | Unknown sequences | 5.336981 | 5.35E-43 |
| LOC111304865 | pentatricopeptide repeat-containing protein At5g16420, mitochondrial | 5.258758 | 6.16E-12 |
| MSTRG.6601 | Unknown sequences | 5.239326 | 4.26E-25 |
| LOC111304849 | uncharacterized protein At4g00950-like | 5.239088 | 1.59E-09 |
| LOC111317018 | protein BRICK 1 | 5.206922 | 4.63E-18 |
| LOC111289594 | transcription factor MYBS1-like | 5.19537 | 4.68E-45 |
| LOC111293048 | probable Histone-lysine N-methyltransferase ATXR5 | 5.169879 | 1.05E-07 |
| LOC111301699 | thioredoxin-like protein CDSP32, chloroplastic | 5.082786 | 1.09E-12 |
| LOC111308611 | transmembrane protein 45B | 5.007227 | 4.88E-64 |
| LOC111275914 | probable L-type lectin-domain containing receptor kinase V.3 | 4.993838 | 2.39E-10 |
| LOC111275243 | probable carboxylesterase SOBER1-like | 4.946835 | 1.17E-27 |
| LOC111277239 | 14 kDa proline-rich protein DC2.15-like | 4.943081 | 2.03E-20 |
| MSTRG.31257 | Unknown sequences | 4.930831 | 5.57E-21 |
| LOC111314105 | proline-rich protein 3-like | 4.903135 | 1.90E-22 |
| LOC111294357 | sulfate transporter 1.3-like | 4.88543 | 9.62E-26 |
| LOC111287737 | G-type lectin S-receptor-like serine/threonine-protein kinase At4g27290 | 4.884873 | 7.88E-13 |
| LOC111283194 | protein STRUBBELIG-RECEPTOR FAMILY 3-like | 4.880131 | 7.65E-37 |
| LOC111286965 | uncharacterized LOC111286965 | 4.813336 | 3.98E-12 |
| MSTRG.23594 | Unknown sequences | 4.807502 | 6.05E-42 |
| MSTRG.34847 | Unknown sequences | 4.764986 | 3.74E-06 |
| MSTRG.2344 | Unknown sequences | 4.724727 | 8.10E-42 |
| LOC111305387 | sm-like protein LSM5 | 4.724291 | 3.39E-06 |
| LOC111285741 | glucan endo-1,3-beta-glucosidase | 4.603205 | 7.69E-25 |
| LOC111311763 | pentatricopeptide repeat-containing protein At5g12100, mitochondrial-like | 4.551911 | 9.29E-06 |
| LOC111310012 | putative disease resistance protein At3g14460 | 4.541002 | 1.14E-05 |
| MSTRG.29429 | Unknown sequences | 4.530304 | 1.07E-05 |
| LOC111294072 | ubiquitin carboxyl-terminal hydrolase 2-like | 4.527706 | 1.14E-05 |
| LOC111293557 | nuclear pore complex protein NUP1-like | 4.502059 | 1.66E-06 |
| MSTRG.3089 | Unknown sequences | 4.463789 | 1.43E-59 |
