## Supplemental Table S4 for "Transcriptome analysis during fruit developmental stages in durian (*Durio zibethinus* Murr.) var. D24"

Table S4 - Differentially down-regulated expressed genes between mature stage and ripening stage of durian fruit pulp. We used FDR: <0.05, Log2 fold change >1.5 and <-1.5.

| **Gene Symbol** | **Gene Name** | **Log2 fold change** | **FDR p-value correction** |
| --- | --- | --- | --- |
| LOC111309322 | sterol 14-demethylase-like | -3.95444 | 0.0005645 |
| LOC111294891 | uncharacterized LOC111294891 | -3.98419 | 0.00020799 |
| LOC111316121 | inositol-3-phosphate synthase-like | -4.03379 | 0.0369311 |
| LOC111277257 | uncharacterized LOC111277257, transcript variant | -4.04538 | 0.00020799 |
| LOC111316987 | homeobox-leucine zipper protein ATHB-6-like | -4.05321 | 0.00020799 |
| LOC111277014 | plant cysteine oxidase 2-like, transcript | -4.06902 | 0.00020799 |
| LOC111315252,  LOC111315261 | peroxisome biogenesis protein 19-2-like, protein transport protein SEC16B homolog | -4.07442 | 0.00056455 |
| LOC111302215 | zinc transporter 6, chloroplastic-like | -4.077 | 0.00020799 |
| LOC111281105 | protein DETOXIFICATION 40-like | -4.10876 | 0.00020799 |
| LOC111299368 | ras-related protein Rab7-like | -4.19375 | 0.00149434 |
| LOC111279856 | protein MARD1-like, transcript variant X1 | -4.21019 | 0.00020799 |
| LOC111275775 | uncharacterized LOC111275775 | -4.22938 | 0.00020799 |
| LOC111296642 | DExH-box ATP-dependent RNA helicase DExH12-like | -4.24855 | 0.00727459 |
| LOC111297335 | trihelix transcription factor PTL-like | -4.25548 | 0.00020799 |
| LOC111294297 | uncharacterized LOC111294297, transcript variant | -4.25692 | 0.00020799 |
| LOC111299310 | probably inactive leucine-rich repeat | -4.26403 | 0.00020799 |
| LOC111289583 | protein DOWNY MILDEW RESISTANCE 6-like | -4.31068 | 0.00020799 |
| LOC111299532 | kelch repeat-containing protein At3g27220-like, | -4.32275 | 0.00020799 |
| LOC111301337 | laccase-4-like | -4.33192 | 0.00882146 |
| LOC111297909 | uncharacterized LOC111297909 | -4.38582 | 0.00020799 |
| LOC111314020 | protein S-acyltransferase 18-like, transcript | -4.39634 | 0.00020799 |
| LOC111306227 | uncharacterized LOC111306227 | -4.47374 | 0.0369311 |
| LOC111285675 | leucine-rich repeat receptor-like protein kinase | -4.53732 | 0.00020799 |
| LOC111277427 | basic proline-rich protein-like | -4.6469 | 0.00020799 |
| LOC111289856 | traB domain-containing protein, transcript | -4.70252 | 0.00020799 |
| LOC111311845 | cytokinin riboside 5'-monophosphate | -4.79953 | 0.0153939 |
| LOC111316168 | transcription factor RAX3-like | -4.81611 | 0.00020799 |
| LOC111316158 | uncharacterized protein At5g65660-like | -5.04522 | 0.00119934 |
| LOC111282836 | glutamate formimidoyltransferase-like | -5.05711 | 0.00020799 |
| LOC111285374 | probable indole-3-pyruvate monooxygenase YUCCA4 | -5.09951 | 0.00020799 |
| LOC111274050 | uncharacterized LOC111274050 | -5.13378 | 0.00020799 |
| LOC111294421 | aquaporin SIP1-1-like | -5.21543 | 0.00020799 |
| LOC111288008 | uncharacterized LOC111288008 | -5.32271 | 0.0153939 |
| LOC111313328 | uncharacterized LOC111313328 | -5.3437 | 0.00020799 |
| LOC111280755 | uncharacterized LOC111280755 | -5.36599 | 0.00020799 |
| LOC111313396 | RING-H RING-H2 finger protein ATL5-like2 finger protein ATL5-like | -5.42314 | 0.00149434 |
| LOC111294357 | sulfate transporter 1.3-like, transcript variant | -5.4384 | 0.00020799 |
| LOC111301699 | thioredoxin-like protein CDSP32, chloroplastic | -5.46366 | 0.0444584 |
| LOC111277239 | 14 14 kDa proline-rich protein DC2.15-like kDa proline-rich protein DC2.15-like | -5.55792 | 0.00491837 |
| LOC111302352 | zinc finger protein 2-like | -5.82013 | 0.00020799 |
| LOC111296030 | protein PIN-LIKES 3-like | -5.915014811 | 0.001385095 |
| LOC111299239 | solute carrier family 25 member 44-like | -5.92814 | 0.000208 |
| LOC111296829 | uncharacterized protein LOC111296829 | -5.951010534 | 0.00347361 |
| LOC111297280 | fructose-bisphosphate aldolase 1, chloroplastic-like | -5.95271 | 0.000208 |
| LOC111307200 | non-specific lipid-transfer protein 3-like | -5.957471163 | 0.000736369 |
| LOC111296871 | serine carboxypeptidase-like 27 isoform X1 | -5.97343927 | 0.002488342 |
| LOC111311845 | cytokinin riboside 5'-monophosphate phosphoribohydrolase LOG3-like | -5.98316846 | 1.71E-23 |
| LOC111311241 | WAT1-related protein At4g08300-like | -6.004389786 | 0.000324017 |
| LOC111276733 | uncharacterized protein LOC111276733 | -6.033095468 | 5.32E-22 |
| LOC111298691 | classical arabinogalactan protein 9-like | -6.039672988 | 0.00050363 |
| LOC111277115 | uncharacterized LOC111277115 | -6.04314 | 0.000208 |
| LOC111286976 | nuclear transcription factor Y subunit C-2-like | -6.056477887 | 0.005249413 |
| LOC111297709 | uncharacterized LOC111297709 | -6.06598 | 0.0431503 |
| LOC111283878 | uncharacterized protein LOC111283878 | -6.155812456 | 9.12E-67 |
| LOC111313342 | myb-related protein 308-like | -6.15936 | 0.0211934 |
| LOC111299239 | solute carrier family 25 member 44-like | -6.160903654 | 2.22E-69 |
| LOC111281093 | uncharacterized protein LOC111281093 | -6.218754239 | 0.003594237 |
| LOC111288040 | ethylene receptor-like | -6.21884 | 3.05E-164 |
| LOC111302663 | 17.3 kDa class I heat shock protein-like | -6.22236 | 4.59E-26 |
| LOC111289805 | mavicyanin-like | -6.24476 | 2.96E-13 |
| LOC111310679 | uncharacterized protein LOC111310679 | -6.254454744 | 0.000407877 |
| MSTRG.20794 | Unknown sequences | -6.25521 | 5.49E-13 |
| LOC111299987 | pentatricopeptide repeat-containing protein At5g39350-like | -6.29017 | 2.05E-11 |
| LOC111296451 | uncharacterized LOC111296451 | -6.29958 | 5.27E-25 |
| LOC111298611 | uncharacterized protein LOC111298611 | -6.311471128 | 0.000469177 |
| MSTRG.33394 | Unknown sequences | -6.31791 | 1.11E-95 |
| LOC111294066 | transcription factor CSA-like | -6.318003633 | 3.40E-39 |
| LOC111283878 | uncharacterized LOC111283878 | -6.31947 | 0.000208 |
| MSTRG.9353 | Unknown sequences | -6.32033 | 8.79E-26 |
| MSTRG.16875 | Unknown sequences | -6.32179 | 1.38E-19 |
| MSTRG.29631 | Unknown sequences | -6.32458 | 5.43E-27 |
| MSTRG.24765 | Unknown sequences | -6.35591 | 1.40E-24 |
| LOC111305971 | LOB domain-containing protein 1-like | -6.375603275 | 7.06E-18 |
| LOC111292205 | U-box domain-containing protein 27-like | -6.378154499 | 1.76E-11 |
| LOC111295167 | serine/arginine-rich splicing factor SC35-like | -6.38201 | 8.53E-18 |
| MSTRG.10291 | Unknown sequences | -6.40003 | 7.84E-22 |
| LOC111304854 | WD repeat-containing protein 44-like | -6.40018 | 3.76E-13 |
| LOC111309528 | uncharacterized protein LOC111309528 | -6.408745624 | 1.86E-05 |
| LOC111310912 | 3,9-dihydroxypterocarpan 6A-monooxygenase-like | -6.409674566 | 8.04E-05 |
| LOC111312349 | uncharacterized LOC111312349 | -6.41738 | 1.20E-47 |
| LOC111282412 | LOB domain-containing protein 42-like | -6.42836 | 0.000208 |
| LOC111292392 | protein trichome birefringence-like 43 | -6.437495032 | 2.19E-05 |
| LOC111277115 | uncharacterized protein LOC111277115 | -6.459764 | 9.03E-57 |
| MSTRG.32718 | Unknown sequences | -6.47771 | 1.60E-41 |
| LOC111289856 | traB domain-containing protein isoform X1 | -6.495710116 | 1.50E-29 |
| LOC111316286 | probable leucine-rich repeat receptor-like protein kinase At1g35710 | -6.49974 | 1.46E-12 |
| LOC111293859 | ubiquitin-conjugating enzyme E2 2 | -6.50177 | 6.27E-15 |
| MSTRG.21056 | Unknown sequences | -6.52476 | 1.93E-13 |
| LOC111295759 | zinc finger protein-like 1 homolog | -6.52661 | 4.08E-107 |
| LOC111305380 | alcohol dehydrogenase 1-like | -6.529963658 | 0.000148747 |
| LOC111313342 | myb-related protein 308-like | -6.531272691 | 6.70E-40 |
| MSTRG.13315 | Unknown sequences | -6.53683 | 1.44E-18 |
| MSTRG.36145 | Unknown sequences | -6.54518 | 3.74E-20 |
| LOC111277239 | 14 kDa proline-rich protein DC2.15-like | -6.558372723 | 6.07E-42 |
| LOC111282412 | protein=LOB domain-containing protein 42-like | -6.570419784 | 9.08E-72 |
| LOC111274856 | calcyclin-binding protein-like | -6.57107 | 6.64E-16 |
| LOC111287652 | stem-specific protein TSJT1-like | -6.584753109 | 7.68E-95 |
| LOC111297851 | eukaryotic translation initiation factor 2D | -6.58581 | 4.00E-154 |
| MSTRG.649 | Unknown sequences | -6.60992 | 7.23E-27 |
| LOC111317381 | uncharacterized protein LOC111317381 | -6.649742902 | 4.33E-06 |
| LOC111312151 | eukaryotic translation initiation factor 2D | -6.66415 | 3.83E-12 |
| LOC111284093 | putative clathrin assembly protein At1g33340 | -6.67899582 | 4.20E-06 |
| MSTRG.24788 | Unknown sequences | -6.72541 | 3.34E-161 |
| LOC111304242 | probable indole-3-pyruvate monooxygenase YUCCA4 | -6.73315992 | 1.99E-05 |
| MSTRG.4788 | Unknown sequences | -6.74233 | 8.39E-51 |
| MSTRG.13935 | Unknown sequences | -6.78338 | 9.04E-46 |
| MSTRG.32418 | Unknown sequences | -6.83039 | 6.56E-22 |
| LOC111309529 | putative F-box protein At1g47765 | -6.832700784 | 1.60E-06 |
| LOC111312958 | uncharacterized LOC111312958 | -6.846234496 | 1.68E-22 |
| LOC111314256 | protein LIFEGUARD 2-like | -6.8786 | 1.67E-14 |
| LOC111280983 | embryo-specific protein ATS3B-like | -6.89428 | 2.42E-35 |
| LOC111292167 | protein SCAI-like | -6.92068 | 1.92E-22 |
| LOC111287086 | probable disease resistance protein At1g58602 | -7.0144 | 1.10E-44 |
| LOC111316151 | probable glycosyltransferase At5g03795 | -7.0346 | 1.64E-13 |
| LOC111303853 | indole-3-acetaldehyde oxidase-like | -7.151661676 | 7.02E-07 |
| MSTRG.34919 | Unknown sequences | -7.18357 | 1.71E-16 |
| LOC111276612 | LOB domain-containing protein 4-like | -7.243266574 | 3.36E-07 |
| MSTRG.5465 | Unknown sequences | -7.24605 | 5.07E-28 |
| LOC111318518 | uncharacterized LOC111318518 | -7.319323386 | 6.13E-07 |
| MSTRG.27668 | Unknown sequences | -7.41324 | 4.31E-56 |
| LOC111298508 | transcription repressor OFP12-like | -7.485007744 | 1.78E-06 |
| LOC111299024 | calmodulin-binding receptor-like cytoplasmic kinase 3 | -7.53485 | 1.50E-20 |
| LOC111296791 | uncharacterized LOC111296791 | -7.57246 | 2.46E-21 |
| LOC111290228 | leucine-rich repeat receptor protein kinase | -7.574169595 | 6.34E-09 |
| LOC111311518 | uncharacterized LOC111311518 | -7.582043405 | 2.08E-18 |
| LOC111317678 | protein SIEVE ELEMENT OCCLUSION B-like | -7.5867 | 4.24E-05 |
| LOC111274060 | probable stress-associated endoplasmic reticulum | -7.600592562 | 2.30E-08 |
| LOC111289620 | probable methyltransferase PMT21 | -7.601359298 | 2.45E-78 |
| LOC111294291 | uncharacterized LOC111294291, transcript variant | -7.613542517 | 2.65E-114 |
| LOC111276730 | uncharacterized LOC111276730 | -7.6196 | 0.011557 |
| LOC111314621 | putative pentatricopeptide repeat-containing protein At1g19290 | -7.63118 | 0.00268 |
| LOC111274856 | calcyclin-binding protein-like | -7.63181 | 0.003399 |
| LOC111285266 | ATP-dependent 6-phosphofructokinase 3-like | -7.6341 | 0.000101 |
| MSTRG.15958 | Unknown sequences | -7.63997 | 1.13E-05 |
| LOC111281786 | probable ubiquitin-conjugating enzyme E2 23 | -7.64286 | 8.40E-06 |
| MSTRG.32418 | Unknown sequences | -7.64693 | 2.86E-05 |
| LOC111278971 | ubiquitin carboxyl-terminal hydrolase 6-like | -7.650372783 | 8.39E-45 |
| MSTRG.27668 | Unknown sequences | -7.65553 | 1.12E-12 |
| LOC111298394 | ferredoxin--nitrite reductase, chloroplastic-like | -7.663576408 | 6.59E-18 |
| LOC111275121 | uncharacterized LOC111275121 | -7.67414 | 0.047556 |
| LOC111318002 | bidirectional sugar transporter SWEET10-like | -7.674514843 | 2.48E-09 |
| LOC111295458 | acyl carrier protein 1, chloroplastic-like | -7.68122 | 0.01566 |
| MSTRG.13583 | Unknown sequences | -7.690612198 | 4.31E-90 |
| MSTRG.2695 | Unknown sequences | -7.711985892 | 4.64E-56 |
| LOC111292167 | protein SCAI-like | -7.71438 | 8.82E-10 |
| LOC111274446 | probable disease resistance protein At5g63020 | -7.72995 | 1.00E-05 |
| LOC111311438 | transport and Golgi organization 2 homolog | -7.73797 | 0.0112713 |
| LOC111289620 | probable methyltransferase PMT21 | -7.75255 | 1.48E-09 |
| LOC111318703 | U-box domain-containing protein 7-like | -7.78136874 | 2.13E-09 |
| LOC111306326 | cysteine-rich repeat secretory protein 60 | -7.78927 | 0.006774 |
| LOC111294291 | uncharacterized LOC111294291 | -7.81045 | 0.000208 |
| MSTRG.13583 | Unknown sequences | -7.81791 | 2.88E-10 |
| MSTRG.13316 | Unknown sequences | -7.823461033 | 4.38E-42 |
| LOC111293859 | ubiquitin-conjugating enzyme E2 2 | -7.82351 | 5.84E-07 |
| LOC111313563 | uncharacterized LOC111313563 | -7.841984559 | 3.24E-10 |
| LOC111304854 | WD repeat-containing protein 44-like | -7.85156 | 0.00259 |
| MSTRG.5465 | Unknown sequences | -7.86892 | 0.000975 |
| MSTRG.2710 | Unknown sequences | -7.87387 | 2.49E-05 |
| MSTRG.5906 | Unknown sequences | -7.87783 | 1.60E-05 |
| LOC111275643 | 65-kDa microtubule-associated protein 1-like | -7.88051 | 0.0153 |
| LOC111311769 | nucleolar transcription factor 1-B-like, | -7.883230096 | 5.49E-11 |
| LOC111291655 | probable magnesium transporter NIPA6 | -7.88339 | 1.11E-05 |
| LOC111292600 | uncharacterized LOC111292600 | -7.88564 | 3.17E-06 |
| LOC111309666 | uncharacterized LOC111309666 | -7.889628613 | 8.25E-10 |
| MSTRG.31953 | Unknown sequences | -7.92144 | 4.06E-06 |
| LOC111291600 | 1,4-alpha-glucan-branching enzyme 1, chloroplastic/amyloplastic-like | -7.93143 | 1.30E-05 |
| LOC111304214 | early nodulin-93-like | -7.957374581 | 1.07E-109 |
| MSTRG.2695 | Unknown sequences | -8.00671 | 1.44E-11 |
| LOC111274241 | transcription factor VOZ1 | -8.010812073 | 3.89E-31 |
| MSTRG.21056 | Unknown sequences | -8.04361 | 0.008704 |
| LOC111278971 | ubiquitin carboxyl-terminal hydrolase 6-like | -8.04697 | 0.000845 |
| MSTRG.5349 | Unknown sequences | -8.05037 | 0.013354 |
| MSTRG.31361 | LOC111290245 | -8.07112 | 4.66E-05 |
| LOC111302666 | UDP-glycosyltransferase 76E2-like | -8.23364 | 2.47E-06 |
| MSTRG.13316 | Unknown sequences | -8.29971 | 7.85E-08 |
| MSTRG.14039 | Unknown sequences | -8.34373 | 6.99E-07 |
| LOC111311552 | polygalacturonase inhibitor-like | -8.365194624 | 1.14E-139 |
| MSTRG.5908 | Unknown sequences | -8.37008 | 1.03E-06 |
| MSTRG.29215 | Unknown sequences | -8.41169 | 0.001064 |
| LOC111311552 | polygalacturonase inhibitor-like | -8.47809 | 0.000208 |
| LOC111287629 | small heat shock protein, chloroplastic-like | -8.502309642 | 6.15E-56 |
| LOC111316686 | probable leucine-rich repeat receptor-like protein kinase At1g35710 | -8.51171 | 6.23E-05 |
| LOC111314600 | GDP-mannose transporter GONST3-like | -8.55671 | 1.18E-06 |
| LOC111278395 | endoglucanase-like | -8.594383006 | 4.03E-100 |
| LOC111299987 | pentatricopeptide repeat-containing protein At5g39350-like | -8.60895 | 3.34E-07 |
| LOC111292881 | uncharacterized protein At1g04910 | -8.71075 | 0.036341 |
| MSTRG.34919 | Unknown sequences | -8.73673 | 0.005391 |
| LOC111316286 | probable leucine-rich repeat receptor-like protein kinase At1g35710 | -8.76366 | 9.38E-08 |
| LOC111311438 | transport and Golgi organization 2 homolog | -8.767848142 | 1.02E-87 |
| LOC111278395 | endoglucanase-like | -8.79611 | 4.36E-13 |
| LOC111287629 | small heat shock protein, chloroplastic-like | -8.96268 | 4.60E-13 |
| LOC111274241 | transcription factor VOZ1 | -8.99109 | 4.23E-06 |
| LOC111296791 | uncharacterized LOC111296791 | -8.99909 | 6.79E-05 |
| LOC111299024 | calmodulin-binding receptor-like cytoplasmic kinase 3 | -9.01691 | 0.000796 |
| LOC111309530 | putative F-box protein At3g16210 | -9.058104145 | 6.66E-20 |
| LOC111314256 | protein LIFEGUARD 2-like | -9.21108 | 2.10E-08 |
| LOC111311518 | uncharacterized LOC111311518 | -9.2998 | 0.019308 |
| LOC111316151 | probable glycosyltransferase At5g03795 | -9.3103 | 0.01446 |
| LOC111282902 | uncharacterized LOC111282902 | -9.697481644 | 1.92E-26 |
| LOC111304216 | early nodulin-93-like | -11.0197445 | 9.60E-139 |
