## Supplemental Table S5 for "Transcriptome analysis during fruit developmental stages in durian (*Durio zibethinus* Murr.) var. D24"

Table S5**:** Categorisation of expressed genes to Gene Ontology terms in YS/MS, YS/RS and MS/RS.

a. Biological Process [P]

| Level | GO ID | GO Name [Biological Process] | YS/MS | YS/RS | MS/RS |
| --- | --- | --- | --- | --- | --- |
| 1 | GO:0008150 | biological process | 1870 | 1149 | 2038 |
| 2 | GO:0008152 | metabolic process | 1416 | 862 | 1509 |
| 2 | GO:0009987 | cellular process | 1401 | 806 | 1415 |
| 2 | GO:0050896 | response to stimulus | 395 | 227 | 393 |
| 2 | GO:0071840 | cellular component organization or biogenesis | 228 | 139 | 219 |
| 2 | GO:0032502 | developmental process | 210 | 0 | 0 |
| 2 | GO:0051179 | localization | 0 | 116 | 208 |
| 3 | GO:0044237 | cellular metabolic process | 1204 | 700 | 1169 |
| 3 | GO:0006807 | nitrogen compound metabolic process | 1075 | 620 | 1084 |
| 3 | GO:0071704 | organic substance metabolic process | 1042 | 520 | 1009 |
| 3 | GO:0044238 | primary metabolic process | 1042 | 520 | 1009 |
| 3 | GO:0009058 | biosynthetic process | 674 | 346 | 652 |
| 3 | GO:0044281 | small molecule metabolic process | 432 | 245 | 471 |
| 3 | GO:0006950 | response to stress | 277 | 174 | 291 |
| 3 | GO:0016043 | cellular component organization | 215 | 128 | 0 |
| 3 | GO:0048856 | anatomical structure development | 210 | 0 | 0 |
| 3 | GO:0009056 | catabolic process | 0 | 161 | 209 |
| 3 | GO:0051234 | establishment of localization | 0 | 116 | 208 |
| 4 | GO:0043170 | macromolecule metabolic process | 673 | 333 | 555 |
| 4 | GO:0044260 | cellular macromolecule metabolic process | 621 | 306 | 534 |
| 4 | GO:1901564 | organonitrogen compound metabolic process | 610 | 308 | 573 |
| 4 | GO:0034641 | cellular nitrogen compound metabolic process | 608 | 374 | 639 |
| 4 | GO:0019538 | protein metabolic process | 541 | 273 | 484 |
| 4 | GO:0006082 | organic acid metabolic process | 248 | 0 | 205 |
| 4 | GO:0006520 | cellular amino acid metabolic process | 248 | 0 | 205 |
| 4 | GO:0006629 | lipid metabolic process | 196 | 120 | 231 |
| 4 | GO:0006810 | transport | 0 | 116 | 208 |
| 5 | GO:0044267 | cellular protein metabolic process | 533 | 271 | 480 |
| 5 | GO:0043412 | macromolecule modification | 490 | 246 | 444 |
| 5 | GO:0036211 | protein modification process | 490 | 246 | 0 |
| 5 | GO:0043436 | oxoacid metabolic process | 248 | 0 | 205 |
| 6 | GO:0006464 | cellular protein modification process | 490 | 246 | 444 |
| 6 | GO:0019752 | carboxylic acid metabolic process | 248 | 0 | 205 |

b. Molecular Function [F]

| Level | GO ID | GO Name [molecular_function] | YS/MS | YS/RS | MS/RS |
| --- | --- | --- | --- | --- | --- |
| 1 | GO:0003674 | molecular function | 2876 | 1646 | 3160 |
| 2 | GO:0005488 | binding | 1699 | 894 | 1601 |
| 2 | GO:0003824 | catalytic activity | 1427 | 730 | 1571 |
| 3 | GO:0043167 | ion binding | 1137 | 597 | 1094 |
| 3 | GO:0016740 | transferase activity | 764 | 0 | 649 |
| 3 | GO:0097159 | organic cyclic compound binding | 631 | 335 | 504 |
| 3 | GO:1901363 | heterocyclic compound binding | 631 | 335 | 504 |
| 3 | GO:0016787 | hydrolase activity | 323 | 166 | 519 |
| 3 | GO:0016740 | transferase activity | 0 | 247 | 0 |
| 4 | GO:0003676 | nucleic acid binding | 631 | 335 | 504 |
| 4 | GO:0016772 | transferase activity, transferring phosphorus-containing groups | 459 | 0 | 358 |
| 5 | GO:0003723 | RNA binding | 359 | 0 | 0 |
| 5 | GO:0016301 | kinase activity | 354 | 0 | 329 |
| 5 | GO:0003677 | DNA binding | 301 | 196 | 344 |

c. Cellular Component [C]

| Level | GO ID | GO Name [cellular_component] | YS/MS | YS/RS | MS/RS |
| --- | --- | --- | --- | --- | --- |
| 1 | GO:0005575 | cellular function | 1989 | 1103 | 2112 |
| 2 | GO:0005623 | cell | 1297 | 775 | 1354 |
| 2 | GO:0044464 | cell part | 1273 | 746 | 1269 |
| 2 | GO:0043226 | organelle | 1049 | 533 | 969 |
| 2 | GO:0032991 | protein-containing complex | 388 | 258 | 595 |
| 3 | GO:0005622 | intracellular | 1208 | 709 | 1220 |
| 3 | GO:0044424 | intracellular part | 1166 | 653 | 1114 |
| 3 | GO:0043229 | intracellular organelle | 1042 | 527 | 952 |
| 3 | GO:0043227 | membrane-bounded organelle | 951 | 454 | 796 |
| 3 | GO:0043228 | non-membrane-bounded organelle | 0 | 149 | 256 |
| 4 | GO:0043231 | intracellular membrane-bounded organelle | 911 | 452 | 757 |
| 4 | GO:0005737 | cytoplasm | 690 | 395 | 663 |
| 4 | GO:0044444 | cytoplasmic part | 558 | 270 | 508 |
| 4 | GO:0043232 | intracellular non-membrane-bounded organelle | 0 | 149 | 256 |
| 5 | GO:0005634 | nucleus | 563 | 308 | 504 |
