## Supplemental Table S6 for "Transcriptome analysis during fruit developmental stages in durian (*Durio zibethinus* Murr.) var. D24"

Table S6**:** KEGG pathway term distribution of expressed genes in YS/MS, YS/RS and MS/RS.

| KEGG pathway terms | Map | YS/MS | YS/RS | MS/RS |
| --- | --- | --- | --- | --- |
| Nucleotide Metabolism |  |  |  |  |
| Purine metabolism | map00230 | 138 | 68 | 185 |
| Pyrimidine metabolism | map00240 | 4 | 7 | 7 |
| Metabolism of cofactors and vitamins |  |  |  |  |
| Thiamine metabolism | map00730 | 138 | 70 | 180 |
| Nicotinate and nicotinamide metabolism | map00760 | 18 | 6 | 5 |
| Ubiquinone and other terpenoid-quinone biosynthesis | map00130 | 16 | 13 | 2 |
| One carbon pool by folate | map00670 | 6 | 0 | 1 |
| Pantothenate and CoA biosynthesis | map00770 | 5 | 3 | 1 |
| Riboflavin metabolism | map00740 | 5 | 2 | 1 |
| Porphyrin and chlorophyll metabolism | map00860 | 4 | 7 | 3 |
| Vitamin B6 metabolism | map00750 | 3 | 0 | 1 |
| Folate biosynthesis | map00790 | 1 | 4 | 2 |
| Biotin metabolism | map00780 | 0 | 1 | 1 |
| Lipoic acid metabolism | map00785 | 0 | 0 | 1 |
| Retinol metabolism | map00830 | 4 | 3 | 2 |
| Carbohydrate metabolism |  |  |  |  |
| **Starch and sucrose metabolism** | **map00500** | **85** | **8** | **15** |
| Amino sugar and nucleotide sugar metabolism | map00520 | 57 | 19 | 15 |
| Glycolysis / Gluconeogenesis | map00010 | 24 | 15 | 11 |
| Galactose metabolism | map00052 | 16 | 15 | 11 |
| Pentose and glucuronate interconversions | map00040 | 14 | 4 | 4 |
| Pentose phosphate pathway | map00030 | 13 | 1 | 2 |
| Inositol phosphate metabolism | map00562 | 9 | 3 | 1 |
| Glyoxylate and dicarboxylate metabolism | map00630 | 8 | 3 | 2 |
| Fructose and mannose metabolism | map00051 | 7 | 6 | 3 |
| Citrate cycle (TCA cycle) | map00020 | 4 | 11 | 4 |
| Butanoate metabolism | map00650 | 2 | 5 | 4 |
| Ascorbate and aldarate metabolism | map00053 | 0 | 2 | 5 |
| C5-Branched dibasic acid metabolism | map00660 | 0 | 0 | 1 |
| Pyruvate metabolism | map00620 | 15 | 10 | 7 |
| Genetic Information Processing |  |  |  |  |
| Aminoacyl-tRNA biosynthesis | map00970 | 26 | 7 | 3 |
| Lipid Metabolism |  |  |  |  |
| Glycerophospholipid metabolism | map00564 | 25 | 24 | 9 |
| Glycerolipid metabolism | map00561 | 16 | 14 | 14 |
| Ether lipid metabolism | map00565 | 14 | 15 | 6 |
| Sphingolipid metabolism | map00600 | 9 | 1 | 2 |
| Arachidonic acid metabolism | map00590 | 5 | 10 | 2 |
| Fatty acid biosynthesis | map00061 | 3 | 0 | 4 |
| Synthesis and degradation of ketone bodies | map00072 | 2 | 2 | 2 |
| Fatty acid degradation | map00071 | 1 | 2 | 3 |
| Linoleic acid metabolism | map00591 | 1 | 0 | 0 |
| alpha-Linolenic acid metabolism | map00592 | 0 | 5 | 6 |
| Cutin, suberine and wax biosynthesis | map00073 | 0 | 0 | 7 |
| Linoleic acid metabolism | map00591 | 0 | 4 | 2 |
| Steroid biosynthesis | map00100 | 0 | 0 | 2 |
| Steroid hormone biosynthesis | map00140 | 3 | 0 | 2 |
| Biosynthesis of other Secondary Metabolites |  |  |  |  |
| Phenylpropanoid biosynthesis | map00940 | 25 | 9 | 5 |
| Caffeine metabolism | map00232 | 14 | 12 | 2 |
| Isoquinoline alkaloid biosynthesis | map00950 | 9 | 0 | 3 |
| Tropane, piperidine and pyridine alkaloid biosynthesis | map00960 | 9 | 1 | 2 |
| Glucosinolate biosynthesis | map00966 | 4 | 3 | 0 |
| Flavonoid biosynthesis | map00941 | 1 | 0 | 0 |
| Biosynthesis of secondary metabolites - unclassified | map00999 | 0 | 0 | 1 |
| Indole alkaloid biosynthesis | map00901 | 1 | 1 | 1 |
| Novobiocin biosynthesis | map00401 | 9 | 0 | 0 |
| Neomycin, kanamycin and gentamicin biosynthesis | map00524 | 5 | 4 | 1 |
| Streptomycin biosynthesis | map00521 | 5 | 4 | 1 |
| Phenazine biosynthesis | map00405 | 0 | 0 | 1 |
| Amino Acid Metabolism |  |  |  |  |
| Cysteine and methionine metabolism | map00270 | 20 | 11 | 9 |
| Arginine and proline metabolism | map00330 | 18 | 7 | 4 |
| Glycine, serine and threonine metabolism | map00260 | 14 | 6 | 4 |
| Valine, leucine and isoleucine biosynthesis | map00290 | 13 | 6 | 2 |
| Alanine, aspartate and glutamate metabolism | map00250 | 12 | 6 | 4 |
| Arginine biosynthesis | map00220 | 12 | 4 | 2 |
| Tyrosine metabolism | map00350 | 12 | 2 | 11 |
| Phenylalanine metabolism | map00360 | 10 | 1 | 0 |
| Phenylalanine, tyrosine and tryptophan biosynthesis | map00400 | 9 | 2 | 5 |
| Valine, leucine and isoleucine degradation | map00280 | 8 | 5 | 2 |
| Histidine metabolism | map00340 | 3 | 1 | 2 |
| Tryptophan metabolism | map00380 | 3 | 1 | 2 |
| Lysine biosynthesis | map00300 | 2 | 2 | 0 |
| Phenylalanine metabolism | map00360 | 0 | 0 | 4 |
| Methane metabolism | map00680 | 7 | 6 | 5 |
| Lysine degradation | map00310 | 33 | 5 | 3 |
| Environmental Information Processing |  |  |  |  |
| Phosphatidylinositol signaling system | map04070 | 19 | 9 | 2 |
| Energy Metabolism |  |  |  |  |
| Carbon fixation in photosynthetic organisms | map00710 | 18 | 9 | 3 |
| Oxidative phosphorylation | map00190 | 10 | 0 | 6 |
| **Sulphur metabolism** | **map00920** | **6** | **2** | **13** |
| Biosynthesis of unsaturated fatty acids | map01040 | 1 | 0 | 1 |
| **Nitrogen metabolism** | **map00910** | **1** | **16** | **2** |
| Carbon fixation pathways in prokaryotes | map00720 | 1 | 8 | 2 |
| Metabolism of other amino acids |  |  |  |  |
| Cyanoamino acid metabolism | map00460 | 6 | 0 | 2 |
| **Glutathione metabolism** | **map00480** | **5** | **18** | **5** |
| beta-Alanine metabolism | map00410 | 2 | 4 | 5 |
| Selenocompound metabolism | map00450 | 1 | 1 | 3 |
| Phosphonate and phosphinate metabolism | map00440 | 0 | 0 | 1 |
| Taurine and hypotaurine metabolism | map00430 | 0 | 3 | 1 |
| D-Glutamine and D-glutamate metabolism | map00471 | 1 | 1 | 2 |
| Glycan biosynthesis and metabolism |  |  |  |  |
| Glycosphingolipid biosynthesis - globo and isoglobo series | map00603 | 6 | 1 | 0 |
| Glycosylphosphatidylinositol (GPI)-anchor biosynthesis | map00563 | 5 | 0 | 0 |
| Glycosphingolipid biosynthesis - ganglio series | map00604 | 4 | 0 | 1 |
| Other glycan degradation | map00511 | 3 | 0 | 2 |
| Glycosaminoglycan degradation | map00531 | 2 | 0 | 1 |
| Glycosphingolipid biosynthesis - lacto and neolacto series | map00601 | 2 | 0 | 0 |
| N-Glycan biosynthesis | map00510 | 0 | 0 | 1 |
| Mucin type O-glycan biosynthesis | map00512 | 2 | 0 | 0 |
| Various types of N-glycan biosynthesis | map00513 | 2 | 0 | 0 |
| Lipopolysaccharide biosynthesis | map00540 | 2 | 1 | 0 |
| Glycosaminoglycan biosynthesis - heparan sulfate / heparin | map00534 | 2 | 2 | 0 |
| Glycosaminoglycan biosynthesis - keratan sulfate | map00533 | 2 | 0 | 0 |
| Lipopolysaccharide biosynthesis | map00540 | 0 | 0 | 2 |
| Peptidoglycan biosynthesis | map00550 | 0 | 0 | 1 |
| Metabolism of terpenoids and polyketides |  |  |  |  |
| Diterpenoid biosynthesis | map00904 | 3 | 0 | 0 |
| Carotenoid biosynthesis | map00906 | 1 | 0 | 1 |
| Terpenoid backbone biosynthesis | map00900 | 2 | 0 | 5 |
| Limonene and pinene degradation | map00903 | 0 | 0 | 1 |
| Sesquiterpenoid and triterpenoid biosynthesis | map00909 | 0 | 0 | 1 |
| Geraniol degradation | map00281 | 2 | 2 | 0 |
| Insect hormone biosynthesis | map00981 | 0 | 0 | 1 |
| Global Pathway |  |  |  |  |
| Biosynthesis of antibiotics | map01130 | 77 | 41 | 33 |
| Aminobenzoate degradation | map00627 | 1 | 2 | 0 |
| Naphthalene degradation | map00626 | 1 | 0 | 1 |
| Steroid degradation | map00984 | 1 | 0 | 2 |
| Drug metabolism - cytochrome P450 | map00982 | 11 | 7 | 4 |
| Drug metabolism - other enzymes | map00983 | 29 | 24 | 7 |
| Nitrotoluene degradation | map00633 | 13 | 11 | 1 |
| Metabolism of xenobiotics by cytochrome P450 | map00980 | 5 | 7 | 3 |
| Styrene degradation | map00643 | 0 | 0 | 5 |
| Caprolactam degradation | map00930 | 0 | 2 | 1 |
| Chloroalkane and chloroalkene degradation | map00625 | 0 | 1 | 2 |
| Environmental Information Processing; Signal transduction |  |  |  |  |
| mTOR signaling pathway | map04150 | 1 | 0 | 1 |
| Immune System |  |  |  |  |
| Th1 and Th2 cell differentiation | map04658 | 33 | 24 | 1 |
| T cell receptor signaling pathway | map04660 | 37 | 24 | 2 |
