## Supplemental Table S7 for "Transcriptome analysis during fruit developmental stages in durian (*Durio zibethinus* Murr.) var. D24"

Table S7 - Heatmap results for softening genes.

| Clust_# | Log2 fold change (Transition from mature stage to ripening stage) | Log2 fold change (Transition from young stage to mature stage) | Gene Symbol | Gene Name |
| --- | --- | --- | --- | --- |
| Cluster_3 | 2.35818 | -3.38712 | LOC111291927 | cellulose synthase-like protein G2 |
| Cluster_3 | 2.40258 | -3.57443 | LOC111286780 | beta-galactosidase 8, X1 |
| Cluster_3 | 2.63924 | -3.47159 | LOC111311338 | U-box domain-containing protein 4-like |
| Cluster_3 | 2.76578 | -3.7092 | LOC111315475 | probable xyloglucan glycosyltransferase 12 |
| Cluster_3 | 2.73849 | -3.7626 | LOC111276893 | cellulose synthase-like protein D3, |
| Cluster_3 | 2.09965 | -4.40848 | LOC111277096 | U-box domain-containing protein 4-like |
| Cluster_3 | 1.97062 | -4.34637 | LOC111293168 | cellulose synthase A catalytic subunit 2 |
| Cluster_3 | 1.97521 | -4.08394 | LOC111296013 | cellulose synthase-like protein D3, |
| Cluster_3 | 3.36652 | -4.06387 | LOC111291010 | expansin-A15-like |
| Cluster_3 | 2.98611 | -2.64462 | LOC111296009 | beta-glucosidase BoGH3B-like |
| Cluster_3 | 3.39961 | -2.53281 | LOC111274640 | beta-galactosidase 17-like, |
| Cluster_3 | 2.90944 | -5.89373 | LOC111303421 | probable xyloglucan |
| Cluster_3 | 2.98039 | -5.86174 | LOC111286758 | beta-galactosidase-like |
| Cluster_3 | 3.44781 | -6.57845 | LOC111289825 | pectin acetylesterase 7-like, |
| Cluster_3 | 3.89605 | -5.41078 | LOC111303852 | beta-glucosidase BoGH3B-like |
| Cluster_3 | 2.7772 | -4.96765 | LOC111284692 | xyloglucan endotransglucosylase/hydrolase |
| Cluster_3 | 2.79502 | -5.2204 | LOC111304738 | U-box domain-containing protein 33-like, |
| Cluster_3 | 2.43431 | -5.2358 | LOC111305459 | U-box domain-containing protein 9-like |
| Cluster_3 | 5.19339 | -6.32033 | LOC111291944 | xyloglucan glycosyltransferase 4-like |
| Cluster_3 | 5.25215 | -6.70076 | LOC111285714 | xyloglucan endotransglucosylase/hydrolase |
| Cluster_3 | 5.46311 | -5.63575 | LOC111274750 | pectinesterase-like |
| Cluster_3 | 6.05873 | -7.36103 | LOC111287953 | xyloglucan endotransglucosylase/hydrolase |
| Cluster_3 | 6.1364 | -7.3922 | LOC111292968 | probable xyloglucan |
| Cluster_3 | 3.94808 | -8.02948 | LOC111303423 | probable xyloglucan |
| Cluster_3 | 3.62892 | -8.29518 | LOC111274745 | pectinesterase 3 |
| Cluster_3 | 9.03817 | -11.0334 | LOC111278395 | endoglucanase-like |
| Cluster_2 | 0 | -3.28599 | LOC111318030 | U-box domain-containing protein 11-like, |
| Cluster_2 | 0 | -3.23435 | LOC111312181 | cellulose synthase A catalytic subunit 5 |
| Cluster_2 | 0 | -3.33978 | LOC111304673 | xyloglucan endotransglycosylase/hydrolase |
| Cluster_2 | 0 | -3.31337 | LOC111296397 | polygalacturonase inhibitor 1 |
| Cluster_2 | 0 | -3.18168 | LOC111280747 | U-box domain-containing protein 4-like |
| Cluster_2 | 0 | -3.18452 | LOC111274630 | beta-galactosidase 1 |
| Cluster_2 | 0 | -3.14884 | LOC111311190 | U-box domain-containing protein 29-like |
| Cluster_2 | 0 | -3.02487 | LOC111317984 | U-box domain-containing protein 6-like, |
| Cluster_2 | 0 | -2.99743 | LOC111279254 | probable xyloglucan glycosyltransferase 12, |
| Cluster_2 | 0 | -2.91779 | LOC111298977 | cellulose synthase A catalytic subunit 1 |
| Cluster_2 | 0 | -2.78119 | LOC111278693 | U-box domain-containing protein 4-like |
| Cluster_2 | 0 | -2.74939 | LOC111309977 | beta-galactosidase, X1 |
| Cluster_2 | 0 | -2.61443 | LOC111306340 | U-box domain-containing protein 21 |
| Cluster_2 | 0 | -2.59038 | LOC111315722 | cellulose synthase A catalytic subunit 5 |
| Cluster_2 | 0 | -3.57817 | LOC111315695 | U-box domain-containing protein 29-like |
| Cluster_2 | 0 | -3.52013 | LOC111288407 | probable xyloglucan |
| Cluster_2 | 0 | -3.46012 | LOC111312526 | probable xyloglucan glycosyltransferase 12 |
| Cluster_2 | 0 | -3.78156 | LOC111312616 | probable xyloglucan 6-xylosyltransferase 5 |
| Cluster_2 | 0 | -3.77137 | LOC111316883 | beta-galactosidase 10-like |
| Cluster_2 | 0 | -3.69578 | LOC111289968 | U-box domain-containing protein 33-like, |
| Cluster_2 | 0 | -4.02578 | LOC111289648 | expansin-like B1, X1 |
| Cluster_2 | 0 | -3.98731 | LOC111275321 | endoglucanase 25-like |
| Cluster_2 | 0 | -3.89443 | LOC111303111 | beta-galactosidase-like |
| Cluster_2 | 0 | -4.18466 | LOC111275267 | U-box domain-containing protein 4-like, |
| Cluster_2 | 0 | -4.0887 | LOC111288463 | U-box domain-containing protein 18 |
| Cluster_2 | 0 | -1.71824 | LOC111317186 | U-box domain-containing protein 5-like, |
| Cluster_2 | 0 | -1.71082 | LOC111278000 | U-box domain-containing protein 30-like |
| Cluster_2 | 0 | -1.68645 | LOC111317549 | U-box domain-containing protein 21-like |
| Cluster_2 | 0 | -1.78624 | LOC111295948 | U-box domain-containing protein 14-like |
| Cluster_2 | 0 | -1.78937 | LOC111280838 | beta-galactosidase 17-like, |
| Cluster_2 | 0 | -1.76237 | LOC111318270 | beta-glucosidase 40-like, X1 |
| Cluster_2 | 0 | -1.52835 | LOC111313678 | cellulose synthase A catalytic subunit 3 |
| Cluster_2 | 0 | -2.04035 | LOC111304739 | U-box domain-containing protein 33-like, |
| Cluster_2 | 0 | -2.06938 | LOC111283356 | probable xyloglucan glycosyltransferase 6, |
| Cluster_2 | 0 | -2.11865 | LOC111314351 | xyloglucan galactosyltransferase MUR3-like |
| Cluster_2 | 0 | -2.10863 | LOC111286079 | pectinesterase/pectinesterase inhibitor |
| Cluster_2 | 0 | -2.0059 | LOC111313303 | xyloglucan endotransglycosylase/hydrolase |
| Cluster_2 | 0 | -2.0191 | LOC111276820 | beta-glucosidase 42-like, X1 |
| Cluster_2 | 0 | -1.9904 | LOC111277138 | pectin acetylesterase 12-like, |
| Cluster_2 | -1.54482 | -1.81964 | LOC111284741 | cellulose synthase A catalytic subunit 1 |
| Cluster_2 | -3.37767 | -1.92762 | LOC111278980 | beta-galactosidase 1-like |
| Cluster_2 | 0 | -6.98037 | LOC111283944 | xyloglucan galactosyltransferase MUR3-like |
| Cluster_2 | 0 | -6.74319 | LOC111291258 | U-box domain-containing protein 19-like |
| Cluster_2 | 0 | -6.42496 | LOC111291886 | beta-galactosidase 3-like |
| Cluster_2 | 0 | -7.32199 | LOC111299213 | cellulose synthase A catalytic subunit 5 |
| Cluster_2 | 0 | -5.11718 | LOC111298783 | probable xyloglucan galactosyltransferase GT17 |
| Cluster_2 | 0 | -5.08399 | LOC111298117 | expansin-like A2 |
| Cluster_2 | 0 | -5.03538 | LOC111281000 | endoglucanase 25-like |
| Cluster_2 | 0 | -4.799 | LOC111304491 | cellulose synthase A catalytic subunit 3 |
| Cluster_2 | 0 | -4.73241 | LOC111299804 | cellulose synthase A catalytic subunit 2 |
| Cluster_2 | 0 | -5.58779 | LOC111307823 | xyloglucan endotransglucosylase/hydrolase |
| Cluster_2 | 0 | -5.41067 | LOC111304740 | U-box domain-containing protein 33-like |
| Cluster_1 | 0 | 3.25174 | LOC111299778 | U-box domain-containing protein 27-like |
| Cluster_1 | 0 | 3.18304 | LOC111282609 | probable xyloglucan |
| Cluster_1 | 0 | 3.41219 | LOC111318703 | U-box domain-containing protein 7-like |
| Cluster_1 | 0 | 3.7957 | LOC111311521 | polygalacturonase-like |
| Cluster_1 | 0 | 2.45962 | LOC111314441 | U-box domain-containing protein 9-like |
| Cluster_1 | 0 | 2.47978 | LOC111290971 | expansin-like B1 |
| Cluster_1 | 0 | 2.29027 | LOC111294605 | probable xyloglucan glycosyltransferase 9, |
| Cluster_1 | 0 | 2.09218 | LOC111281864 | putative U-box domain-containing protein 42, |
| Cluster_1 | -1.56103 | 4.61035 | LOC111285245 | probable xyloglucan galactosyltransferase GT11, |
| Cluster_1 | -3.12501 | 2.02888 | LOC111317423 | pectinesterase 3-like |
| Cluster_1 | 0 | 8.43397 | LOC111311552 | polygalacturonase inhibitor-like |
