## Supplemental Table S8 for "Transcriptome analysis during fruit developmental stages in durian (*Durio zibethinus* Murr.) var. D24"

Table S8 - Heatmap results for signal transduction

1. Genes involved in ethylene biosynthesis

| Gene Symbol | Gene Name | Log2 fold change (Transition from young stage to mature stage) | Log2 fold change (Transition from mature stage to ripening stage) |
| --- | --- | --- | --- |
| LOC111314546 | 1-aminocyclopropane-1-carboxylate oxidase | -5.10515 | 4.57221 |
| LOC111296866 | 1-aminocyclopropane-1-carboxylate oxidase | -5.42301 | 3.30056 |
| LOC111317524 | 1-aminocyclopropane-1-carboxylate oxidase | -2.24663 | 2.43102 |
| LOC111306707 | 1-aminocyclopropane-1-carboxylate oxidase | 2.41753 | 0 |
| LOC111305273 | 1-aminocyclopropane-1-carboxylate oxidase | -3.95841 | 0 |
| LOC111274040 | 1-aminocyclopropane-1-carboxylate oxidase-like | -4.43793 | 3.24625 |
| LOC111300706 | 1-aminocyclopropane-1-carboxylate synthase | -2.09305 | -3.28987 |
| LOC111313897 | 1-aminocyclopropane-1-carboxylate synthase-like | -2.80886 | 1.94419 |
| LOC111304783 | 1-aminocyclopropane-1-carboxylate synthase-like | 1.61627 | 0 |
| LOC111299814 | S-adenosylmethionine carrier 1 | -2.27598 | 0 |
| LOC111286226 | S-adenosylmethionine decarboxylase | -2.96624 | 5.16461 |
| LOC111287069 | S-adenosylmethionine decarboxylase | -2.65215 | 3.66747 |
| LOC111318202 | S-adenosylmethionine decarboxylase | -2.65431 | 2.18832 |
| LOC111282309 | S-adenosylmethionine decarboxylase | -1.74857 | 1.8788 |
| LOC111304250 | S-adenosylmethionine decarboxylase | -4.8777 | 0 |
| LOC111304951 | S-adenosylmethionine synthase 2 | -1.72133 | -1.5831 |
| LOC111279193 | S-adenosylmethionine synthase 2 | -5.6817 | 2.0695 |

2. Genes involved in ethylene signalling pathway

| Gene Symbol | Gene Name | Log2 fold change (Transition from young stage to mature stage) | Log2 fold change (Transition from mature stage to ripening stage) |
| --- | --- | --- | --- |
| LOC111284689 | EIN3-binding F-box protein 1-like | 1.96413 | -1.74249 |
| LOC111303272 | EIN3-binding F-box protein 1-like | 2.7235 | 0 |
| LOC111279153 | ethylene receptor 2-like | 3.36653 | 0 |
| LOC111306700 | ethylene receptor 2-like | 2.8125 | -2.07699 |
| LOC111291911 | ethylene receptor 2-like | 2.05944 | -2.76811 |
| LOC111305038 | ethylene response sensor 1-like | -1.73411 | 0 |
| LOC111295068 | ethylene-overproduction protein 1-like | 0 | 1.82714 |
| LOC111286414 | ethylene-overproduction protein 1-like | 0 | -1.80587 |
| LOC111282084 | mitogen-activated protein kinase 17-like | -1.65344 | 3.15983 |
| LOC111275978 | mitogen-activated protein kinase 19-like | -1.95352 | 0 |
| LOC111284718 | mitogen-activated protein kinase 3 | -2.2638 | 2.62908 |
| LOC111282030 | mitogen-activated protein kinase 9-like | -1.67659 | 0 |
| LOC111289681 | mitogen-activated protein kinase homolog | 2.62611 | 0 |
| LOC111301438 | mitogen-activated protein kinase homolog NTF6 | -2.62893 | 0 |
| LOC111275659 | mitogen-activated protein kinase kinase kinase | -8.08745 | 4.69845 |
| LOC111306687 | mitogen-activated protein kinase kinase kinase | -6.06221 | 3.81307 |
| LOC111290499 | mitogen-activated protein kinase kinase kinase | -2.24909 | 1.81575 |
| LOC111315610 | mitogen-activated protein kinase kinase kinase | -2.04864 | 1.60796 |
| LOC111285643 | mitogen-activated protein kinase kinase kinase | 2.69438 | 0 |
| LOC111315404 | mitogen-activated protein kinase kinase kinase | -2.69059 | 0 |
| LOC111292037 | mitogen-activated protein kinase kinase kinase | -3.94503 | 0 |
| LOC111301871 | mitogen-activated protein kinase kinase kinase | -4.09542 | 0 |
| LOC111318818 | mitogen-activated protein kinase kinase kinase | -7.37864 | 0 |
| LOC111312465 | serine/threonine-protein kinase CTR1-like | -2.18159 | 2.47455 |

3. Genes involved in ethylene response

| Gene Symbol | Gene Name | Log2 fold change (Transition from young stage to mature stage) | Log2 fold change (Transition from mature stage to ripening stage) |
| --- | --- | --- | --- |
| LOC111290879 | ethylene-responsive transcription factor | -3.9508 | 3.77611 |
| LOC111304158 | ethylene-responsive transcription factor | -4.23178 | 3.28427 |
| LOC111280307 | ethylene-responsive transcription factor | -4.65325 | 3.01434 |
| LOC111301172 | ethylene-responsive transcription factor | -2.91794 | 2.85424 |
| LOC111292151 | ethylene-responsive transcription factor | -3.99649 | 2.64843 |
| LOC111292116 | ethylene-responsive transcription factor | -2.92381 | 2.64577 |
| LOC111296272 | ethylene-responsive transcription factor | -5.15796 | 2.63686 |
| LOC111304157 | ethylene-responsive transcription factor | -2.9678 | 2.0609 |
| LOC111299257 | ethylene-responsive transcription factor | -5.07856 | 1.59693 |
| LOC111306880 | ethylene-responsive transcription factor | 4.59281 | 0 |
| LOC111314102 | ethylene-responsive transcription factor | 2.58359 | 0 |
| LOC111306123 | ethylene-responsive transcription factor | 1.93404 | 0 |
| LOC111299205 | ethylene-responsive transcription factor | 1.69509 | 0 |
| LOC111285490 | ethylene-responsive transcription factor | -1.82564 | 0 |
| LOC111279035 | ethylene-responsive transcription factor | -1.85751 | 0 |
| LOC111287270 | ethylene-responsive transcription factor | -1.9127 | 0 |
| LOC111318106 | ethylene-responsive transcription factor | -2.21131 | 0 |
| LOC111295193 | ethylene-responsive transcription factor | -2.38553 | 0 |
| LOC111279759 | ethylene-responsive transcription factor | -2.45319 | 0 |
| LOC111299848 | ethylene-responsive transcription factor | -2.52998 | 0 |
| LOC111300672 | ethylene-responsive transcription factor | -2.83074 | 0 |
| LOC111312376 | ethylene-responsive transcription factor | -2.83595 | 0 |
| LOC111311178 | ethylene-responsive transcription factor | -3.07038 | 0 |
| LOC111309917 | ethylene-responsive transcription factor | -3.4791 | 0 |
| LOC111305283 | ethylene-responsive transcription factor | -3.66669 | 0 |
| LOC111309281 | ethylene-responsive transcription factor | -3.80244 | 0 |
| LOC111308283 | ethylene-responsive transcription factor | -3.85505 | 0 |
| LOC111293640 | ethylene-responsive transcription factor | -4.00151 | 0 |
| LOC111311243 | ethylene-responsive transcription factor | -4.20813 | 0 |
| LOC111316955 | ethylene-responsive transcription factor | -4.80753 | 0 |
| LOC111300396 | ethylene-responsive transcription factor | -4.84218 | 0 |
| LOC111309418 | ethylene-responsive transcription factor | -5.50105 | 0 |
| LOC111315585 | ethylene-responsive transcription factor | -5.68808 | 0 |
| LOC111315804 | ethylene-responsive transcription factor | -7.35398 | 0 |
| LOC111285645 | ethylene-responsive transcription factor | 1.59995 | -1.5414 |
| LOC111304137 | ethylene-responsive transcription factor | -2.61087 | -2.96681 |
| LOC111299480 | ethylene-responsive transcription factor | 3.77712 | -3.3807 |
| LOC111307395 | ethylene-responsive transcription factor 1B | -3.30071 | 0 |
| LOC111276899 | ethylene-responsive transcription factor 2-like | -1.86055 | 2.74019 |
| LOC111291204 | ethylene-responsive transcription factor 2-like | -3.92165 | 2.13609 |
| LOC111290760 | ethylene-responsive transcription factor 2-like | -2.68887 | 0 |
| LOC111276522 | ethylene-responsive transcription factor 4-like | -2.1634 | 0 |
| LOC111278637 | ethylene-responsive transcription factor 4-like | -4.33223 | 0 |
| LOC111275030 | ethylene-responsive transcription factor 4-like | -4.74314 | 0 |
| LOC111291396 | ethylene-responsive transcription factor ERF061 | -7.03836 | 5.12252 |
| LOC111290737 | ethylene-responsive transcription factor ERF113 | 1.57019 | 0 |
