## Supplemental Table S9 for "Transcriptome analysis during fruit developmental stages in durian (*Durio zibethinus* Murr.) var. D24"

Table S9 - Genes and metabolizing enzymes involved in starch and sucrose metabolism pathway.

1. Identified genes in starch and sucrose metabolism pathway

a. Transition from young to mature stage

| Locus (Chr:Start-End) | Metabolizing enzymes | Gene symbol | Description |
| --- | --- | --- | --- |
| NW_019167937.1:31150242-31160456 | EC:5.3.1.9 - isomerase | LOC111312441 | *uncharacterized LOC111312441* |
| NW_019167960.1:7870010-7870999 | EC:2.4.1.34 - synthase | LOC111316836 | *uncharacterized LOC111316836* |
| NW_019168015.1:14014159-14021413 | EC:2.4.1.34 - synthase | LOC111278181 | callose synthase 11-like |
| NW_019167938.1:15505567-15524846 | EC:2.4.1.18 - branching enzyme | LOC111314672 | 1,4-alpha-glucan-branching enzyme 2-1,chloroplastic/amyloplastic-like |
| NW_019168426.1:6282-21213 | EC:2.4.1.18 - branching enzyme | LOC111291600 | 1,4-alpha-glucan-branching enzyme 1,chloroplastic/amyloplastic-like |
| NW_019167960.1:9302267-9305462 | EC:2.7.7.27 - adenylyltransferase | LOC111316856 | glucose-1-phosphate adenylyltransferase large subunit 1, chloroplastic-like |
| NW_019168481.1:25094521-25097000 | EC:2.7.1.4 - fructokinase (phosphorylating) | LOC111293926 | fructokinase-like 1, chloroplastic |
| NW_019167882.1:22167487-22174212 | EC:2.7.1.1 - hexokinase type IV glucokinase | LOC111303941 | hexokinase-3-like |

b. Transition from young to ripening stage

| Locus (Chr:Start-End) | Metabolizing enzymes | Gene symbol | Description |
| --- | --- | --- | --- |
| NW_019167849.1:10546605-10554032 | EC:2.4.1.34 - synthase | LOC1112911135 | callose synthase 12-like |
| NW_019167882.1:22167487-22174212 | EC:2.7.1.1 - hexokinase type IV glucokinase | LOC111303941 | hexokinase-3-like |
| NW_019168481.1:21661427-21668986 | EC:2.4.1.12 - synthase (UDP-forming) | LOC111293564 | probable cellulose synthase A catalytic subunit 8 [UDP-forming] |
| NW_019168481.1:8521803-8523746 | EC:2.4.1.12 - synthase (UDP-forming) | LOC111293942 | 4-alpha-glucanotransferase DPE2 |

c. Transition from mature to ripening stage

| Locus (Chr:Start-End) | Metabolizing enzymes | Gene symbol | Description |
| --- | --- | --- | --- |
| NW_019167937.1:31150242-31160456 | EC:3.2.1.39 - endo-1,3-beta-D-glucosidase | LOC111312441 | *uncharacterized LOC111312441* |
| NW_019168015.1:14014159-14021413 | EC:2.4.1.34 - synthase | LOC111278181 | callose synthase 11-like |
| NW_019167960.1: 9302267-9305462 | EC:2.7.7.27 - adenylyltransferase | LOC111316856 | glucose-1-phosphate adenylyltransferase large subunit 1, chloroplastic-like |
| NW_019167882.1:22167487-22174212 | EC:2.7.1.1 - hexokinase type IV glucokinase | LOC111303941 | hexokinase-3-like |
| NW_019168481.1:25094521-25097000 | EC:2.7.1.4 - fructokinase (phosphorylating) | LOC111293926 | fructokinase-like 1, chloroplastic |
| NW_019167960.1:7870010-7870999 | EC:2.7.7.27 - adenylyltransferase | LOC111316836 | *uncharacterized LOC111316836* |
| NW_019167937.1:2,192,345-2,196,476 | EC:3.1.3.24 - phosphatase | LOC111311629 | sucrose-phosphatase 2-like |
| NW_019168159.1:25,486,523-25,495,125 | EC:2.4.1.13 - synthase | LOC111286073 | probable sucrose-phosphate synthase 1 |
| NW_019167937.1:33,919,794-33,922,417 | EC:3.2.1.39 - endo-1,3-beta-D-glucosidase | LOC111312703 | glucan endo-1,3-beta-D-glucosidase-like |
| NW_019167838.1:4,158,516-4,160,995 | EC:3.2.1.4 - endo-1,4-beta-D-glucanase | LOC111278395 | endoglucanase-like |
| NW_019167838.1:6,756,338-6,763,193 | EC:2.4.1.12 - synthase (UDP-forming) | LOC111287763 | probable trehalose-phosphate phosphatase D |
| NW_019168381.1:26,223,140-26,228,064 | EC:2.4.1.12 - synthase (UDP-forming) | LOC111290775 | acid beta-fructofuranosidase 1, vacuolar-like |
| NW_019167904.1:18,187,419-18,190,194 | EC:3.2.1.39 - endo-1,3-beta-D-glucosidase | LOC111306779 | beta-amylase 1, chloroplastic-like |
| NW_019168026.1:2,088,092-2,091,258 | EC:3.2.1.2 - saccharogen amylase | LOC111279622 | beta-amylase 1, chloroplastic-like |
| NW_019167904.1:20,121,034-20,122,845 | EC:3.2.1.39 - endo-1,3-beta-D-glucosidase | LOC111306922 | glucan endo-1,3-beta-glucosidase, basic vacuolar  isoform |
| NW_019167937.1_33173237_33175372_plus | EC:3.2.1.1 - glycogenase | LOC111312611 | alpha-amylase like |

2. No of sequences in metabolizing enzymes of starch and sucrose metabolism pathway.

| Enzyme | No of seqs in enzyme | Stage of Growth |
| --- | --- | --- |
| EC:5.3.1.9 - isomerase | 9 | young to mature stage |
| EC:2.4.1.34 - synthase | 4 | young to mature stage |
| EC:2.4.1.18 - branching enzyme | 52 | young to mature stage |
| EC:2.7.7.27 - adenylyltransferase | 15 | young to mature stage |
| EC:2.7.1.1 - hexokinase type IV glucokinase | 5 | young to mature stage |
| EC:2.7.1.4 - fructokinase (phosphorylating) | 1 | young to mature stage |
| Enzyme | No of seqs in enzyme | Stage of Growth |
| EC:2.4.1.34 - synthase | 1 | young to ripening stage |
| EC:2.7.1.1 - hexokinase type IV glucokinase | 4 | young to ripening stage |
| EC:2.4.1.12 - synthase (UDP-forming) | 1 | young to ripening stage |
| EC:2.4.1.25 - disproportioning enzyme | 2 | young to ripening stage |
| Enzyme | No of seqs in enzyme | Stage of Growth |
| EC:5.3.1.9 - isomerase | 9 | mature to ripening stage |
| EC:2.4.1.34 - synthase | 4 | mature to ripening stage |
| EC:2.7.7.27 - adenylyltransferase | 15 | mature to ripening stage |
| EC:2.7.1.1 - hexokinase type IV glucokinase | 5 | mature to ripening stage |
| EC:2.7.1.4 - fructokinase (phosphorylating) | 1 | mature to ripening stage |
| EC:3.1.3.24 - phosphatase | 2 | mature to ripening stage |
| EC:3.2.1.21 - gentiobiase | 2 | mature to ripening stage |
| EC:3.2.1.1 - glycogenase, | 1 | mature to ripening stage |
| EC:3.2.1.4 - endo-1,4-beta-D-glucanase, | 2 | mature to ripening stage |
| EC:2.4.1.13 - synthase | 5 | mature to ripening stage |
| EC:3.1.3.12 - trehalose 6-phosphatase, | 1 | mature to ripening stage |
| EC:3.2.1.26 - invertase, | 1 | mature to ripening stage |
| EC:3.2.1.2 - saccharogen amylase, | 3 | mature to ripening stage |
| EC:3.2.1.39 - endo-1,3-beta-D-glucosidase | 2 | mature to ripening stage |
| EC:3.2.1.48 - alpha-glucosidase | 1 | mature to ripening stage |
| EC:2.4.1.14 - synthase | 4 | mature to ripening stage |
| EC:3.2.1.20 - maltase | 1 | mature to ripening stage |
| EC:2.4.1.12 - synthase (UDP-forming) | 14 | mature to ripening stage |
| EC:2.4.1.1 - phosphorylase | 4 | mature to ripening stage |
